# Evolution of cross-tolerance to metals in yeast

**DOI:** 10.1101/2025.02.12.637395

**Authors:** Anna L. Bazzicalupo, Penelope C. Kahn, Eully Ao, Joel Campbell, Sarah P. Otto

## Abstract

Organisms often face multiple selective pressures simultaneously (e.g., mine tailings with multiple heavy metal contaminants), yet we know little about when adaptation to one stressor provides cross-tolerance or cross-intolerance to other stressors. To explore the potential for cross-tolerance, we first adapted *Saccharomyces cerevisiae* to high concentrations of six single metals in a short-term evolutionary rescue experiment. We then measured the cross-tolerance of each metal-adapted line in the other five metals. We generated and tested three predictors for the degree of cross-tolerance, based on the similarity between pairs of metal environments in (1) their physiochemical properties, (2) the overlap in genes known to impact tolerance to both metals, and (3) their co-occurrence in the environment. None of these predictors explained significant variation in cross-tolerance. Instead, we observed that adapted lines in one metal were frequently cross-tolerant to certain metals (manganese and nickel) and intolerant to others (cobalt and zinc). Furthermore, cross-tolerance between pairs of metals was not reciprocal, with mutations accumulating in one metal (e.g., copper) providing adaptation to another metal (e.g., manganese), but not *vice versa*. Evolved lines also differed in their degree of specialization, with lines evolved in manganese or copper more specialized to that metal, but lines evolved in cobalt or zinc more generally tolerant. To determine the genetic basis of these metal adaptations, we sequenced the genomes of 109 metal-adapted yeast lines. The SNP mutation spectrum was significantly different in cadmium, cobalt, and manganese than expected in a mutation accumulation experiment in *S. cerevisiae*. In addition, two lines were highly mutated, bearing defects in DNA repair genes (both in manganese). Thirteen genes exhibited parallel adaptation to different metals; three of these genes generated broad cross-tolerance. Several mutations were found in vacuolar transporter genes, suggesting an important role for vacuolar proteins in adapting to metal stress. Our results with these metal-adapted lines indicate that cross-tolerance is challenging to predict, depending on the combined stressors experienced and the nature of the mutations involved.

## Introduction

Organisms evolving in a particular environment accumulate pleiotropic genetic responses in other environments that remain unrealized and mostly untested^1^. As the world changes rapidly around us, predicting when cross-tolerance or cross-intolerance are expected would be an important tool for conservation and ecosystem management. The results and analyses of Jerison et al.^2^ suggest that there is a relationship between the degree of environmental similarity and the amount of cross-tolerance expected. Tracking yeast undergoing experimental evolution in 11 environments, Jerison et al.^2^ found that chance played a major role in the probability of generating specialists and generalists, depending on the frequency and fitness costs of mutations providing narrow versus broad tolerance.

Industrialization and technological innovation over the past century have led to a massive increase in heavy metal pollution^34^. Heavy metals induce a multitude of stresses that affect a variety of physiological responses, and exposure can be lethal^5^. Humans frequently take advantage of the toxic properties of metals, using elevated concentrations for microbial control (e.g., in wine production)^6^. These pollutants, especially lead and mercury, have been recognized for their distinctly negative effects on human health, leading to global efforts to reduce their rates of use and release into the environment^7,8^. Other metals continue to be emitted to the biosphere via mining, smelting, leaching from plastics and electronics in landfills, agricultural runoff, automobiles, and roadwork^3^.

Several metals, such as zinc, cobalt, nickel, copper, and manganese, act as micronutrients in the eukaryotic cell that are essential in trace amounts for proper cell functioning but are toxic at high concentrations^9^. Metal ions induce the production of reactive oxygen species (ROS), which have the potential to damage membranes, proteins, and DNA, becoming lethal when concentrations reach high levels^3^. Evolution of metal resistance can occur rapidly, with adaptations involving several gene pathways and physiological mechanisms^10–13^. For example, strains of *E. coli* evolved in a high concentration of silver nanoparticles gain mutations impacting ion sensing, nucleotide biosynthesis, and transcription^10^.

Micronutrients cannot be completely excluded from cells^14^, and fine-tuned regulatory systems ensure a balance between heavy metal excess and deficiency. These systems include transporters for import and export, chelators for metal binding, and antioxidants to counteract ROS production. Despite some similarities in their effects (e.g., ROS production), exposure to different metals can induce dramatically different consequences. For example, contact with metallic copper induces extensive cytoplasmic membrane damage in both *Saccharomyces cerevisiae* and *Candida albicans*^15^, while other metals are known to be mutagenic. Cadmium toxicity leads to recombinagenic activity, induces deletion mutations via oxidative stress^16^, and inhibits mismatch repair of small misalignments^17^. Cobalt is mutagenic to mitochondrial DNA but less so to nuclear DNA^18–20^. Nickel can cause changes in chromatin structure that alter gene expression^21^. Differences in the targets of these metals suggest that adaptive mutations selected in one environment may not improve tolerance to other metals. On the other hand, yeast have evolved general stress response strategies that may facilitate cross-tolerance of adaptative mutations (e.g., chelation or compartmentalization).

To investigate the cross-tolerance of mutations evolved in the presence of different metals, we generated rapidly evolved lines via evolutionary rescue experiments. We exposed replicate populations of *S. cerevisiae* to five essential metal co-factors (cobalt, copper, manganese, nickel, and zinc) and one toxic metal (cadmium) at concentrations that caused most populations to go extinct but allowed for evolutionary rescue through rare, large-effect mutations (Fig. 1). By studying these adapted lines, we document the prevalence and genetic basis of cross-tolerance and cross-intolerant adaptive mutations and seek to better predict the scope for evolutionary rescue when organisms face additional stressors (Fig. 1 and Fig. S1).

**Figure 1.**
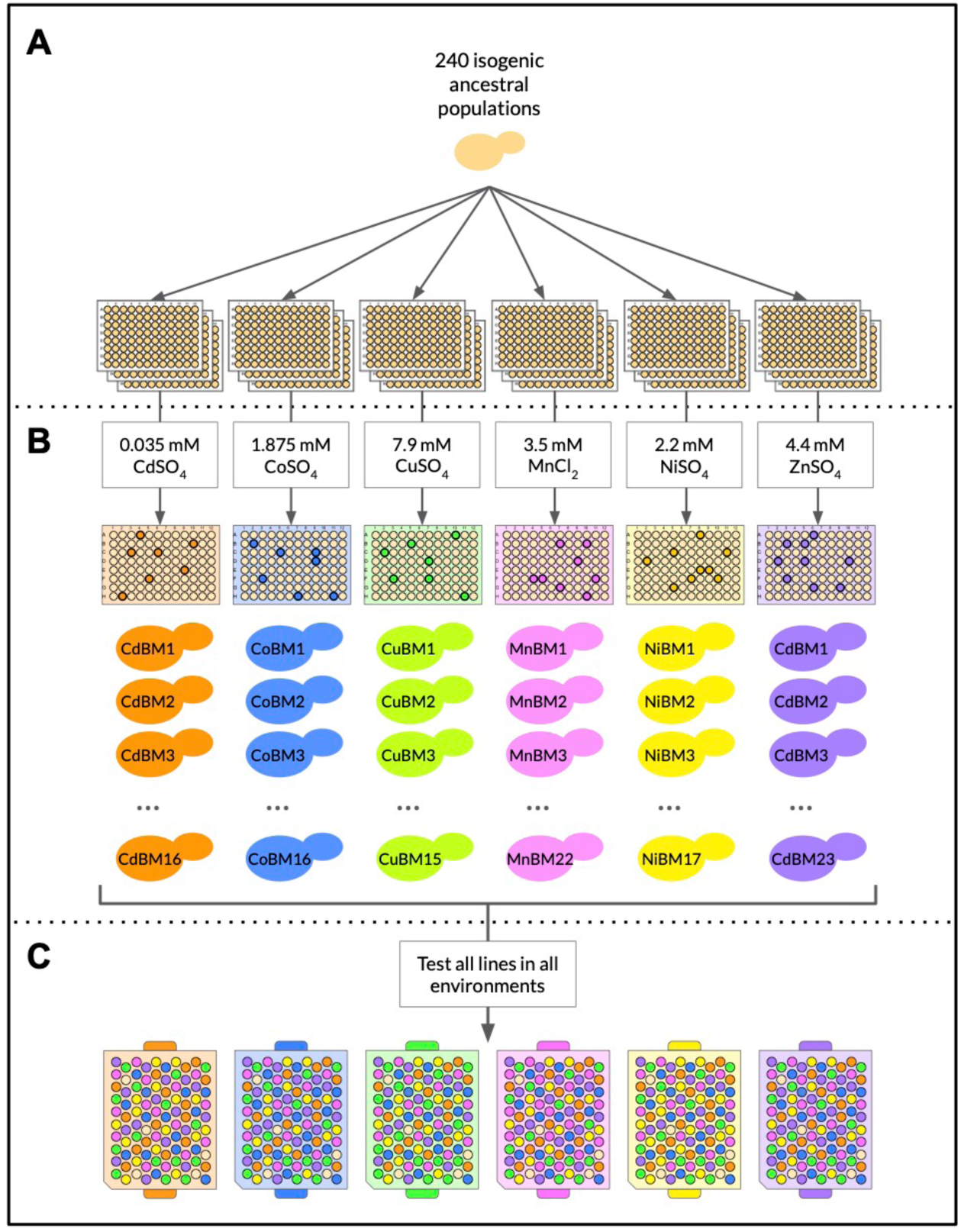
Diagram of experimental design for evolution experiment and fitness tests. (A) Two-hundred and forty single colony isolates of strain W303 were inoculated into liquid YPAD in three 96-well plates, yielding 240 ancestral lines. See figure S1 for details of the block design. (B) The 240 ancestral lines were exposed to a lethal dose of CdSO_4_, CoSO_4_, CuSO_4_, MnCl_2_, NiSO_4_, and ZnSO_4_, prompting evolutionary rescue, and yielding between 15 and 23 evolved lines for each metal. (C) The growth rates of all evolved lines were measured in each metal environment using a Bioscreen C automatic plate reader.

We developed three *a priori* predictions for the degree of cross-tolerance between pairs of metals:

### Prediction I: Similarity in reactivity of metals predicts cross-tolerance

We predict that cross-tolerance would be observed more often for pairs of metals that have more similar oxidizing and reducing potential.

### Prediction II: Similarity in the sets of genes known to affect metal sensitivity predicts cross-tolerance

We predict that cross-tolerance would be observed more often for pairs of metals that have a greater overlap in the set of genes affecting tolerance to those metals, as reported in the *Saccharomyces Genome Database*^22^.

### Prediction III: Similarity in metal abundance across environments predicts cross-tolerance

We predict that cross-tolerance would be observed more often for metal pairs that commonly co-occur in natural environments, reflecting an evolutionary history of greater exposure to these combinations.

Cross-tolerance was not well predicted by any of these measures. Instead, evolution in certain metals led frequently to generalists that better tolerated many metals (cross-tolerance), while evolution in other metals lead to specialists that performed poorly across metals (cross-intolerance). Furthermore, mutations were observed repeatedly in thirteen genes among lines adapted to different metals, despite arising frequently, only three of these lines were more broad-scale cross-tolerant on average. Thus, the environmental conditions first encountered and the adaptive steps first taken matter disproportionately in predicting tolerance to multiple metal stressors.

## Materials and Methods

### Strains and Media

Yeast populations used in this experiment were derived from haploid lines of the lab strain W303. A sample of frozen W303 stock was inoculated in liquid rich medium (YPD (RPI Research Products Y20090) + adenine, referred to as YPAD onward) and grown overnight at 30°C. A 1:1000 dilution was performed and spread on YPAD plates in order to bottleneck the initial population down to a single cell (one cell founding each replicate well). After two days of growth at 30°C, 240 single colonies were isolated and each used to inoculate one well of a 96-deepwell plate containing 1 mL of YPAD (three plates in total with 80 inoculated wells each). The plates were then allowed to grow overnight at 30°C for use in the evolutionary rescue experiments.

To obtain a tolerance profile and determine concentrations of each metal beyond which growth is not typically observed, a whole population sample of W303 was subdivided and exposed to an increasing series of metal concentrations. The tolerance profiles were obtained using a Bioscreen C, a microbiology workstation that automatically tracks growth trajectories by measuring turbidity of liquid cultures in 100-well Bioscreen plates at regular intervals using wide band filters (Oy Growth Curves Ab Ltd., 2009). Throughout, the fitness measure that we use is the maximum mitotic growth rate obtained using a loess fit to the log of the growth curves obtained from one well in the Bioscreen C (see details in Gerstein et al.^11^ and https://github.com/joelkcamp/crossTolerance_bazzicalupoEtAl for code). Maximum growth rates were measured across ten metal concentrations (chosen to span the transition from non-lethal to lethal concentrations based on pilot experiments), and tolerance curves were determined for the lab yeast strains W303 and BY4741, which was the ancestral background of the lines evolved in Gerstein et al.^11^. Specifically, W303 single colony cultures were diluted 1:11 and grown overnight in 96 well plates to obtain stock cultures. Bioscreen plates were then inoculated with a 1:100 dilution of these stocks along with a single metal at different concentrations (mM) to determine the lowest concentration of “no growth”, i.e., where the maximum growth rate was similar to that observed in blank wells (see Fig. S2). Further testing around the observed point of no growth was conducted to narrow down the lowest concentration that conferred no growth, which was used as the metal concentration in the rescue experiments (see Table S1 for concentrations used in the experiment).

To gain further insight into the genetic mechanisms conferring tolerance, we also assayed metal tolerance of lines evolved from other experiments. To estimate the impact of mitochondrial function on metal tolerance, we assayed ten random lines that were respiration deficient due to a loss of mitochondrial function (“petite”, so-called due to small colony size when grown on solid growth media) and ten normal (“grande”) lines that had arisen in another experiment within the lab (selecting on outcrossing rates, not metal media)^23^. We also used Copper Beneficial Mutation (CBM) lines from Gerstein et al.^11^ (evolved from the same ancestral stock) to test if whole-chromosome duplications provided cross-tolerance to other metals.

### Evolutionary rescue experimental design

We measured the optical density of each of the 240 cultures grown in the 96-deepwell plates with YPAD and normalized culture concentrations by diluting all to the lowest concentration recorded. For each of the six metals used, metal stock solutions were added to YPAD in 240 wells across three 96-deepwell plates to obtain the minimum lethal concentration (Table S1). Each well was then inoculated with starting culture from one of the 240 isolated colonies, using the same block design for all metals (Fig. 1 and Fig. S1). Identical adaptive mutations observed across metals from the same well and plate position are thus considered to have arisen during the growth of that colony (non-independent) and discussed below. The plates were shaken (200 rpm) at 30°C for 14 days. All boxes were checked daily by visual examination, and growth was recorded when we saw settling of cells at the bottom of a well. Because this initial growth is often faint, we then allowed 24 hours to confirm growth, after which the well was manually mixed, and two sample tubes were frozen at -80°C by placing 400 µL of culture in 15% glycerol. In order to screen for genetic adaptations (as opposed to physiological or plastic responses), we started each evolved line from frozen in liquid YPAD and allowed them to grow for 2 or 3 days until growth could be observed. We then inoculated 10 μL of this culture into 1 mL YPAD + metal at the lethal concentrations used in the experiment. Wells that did not exhibit growth within 4 days were eliminated from the experiment. We also tested each of the evolved lines for respiratory mutations by spot plating them onto glycerol plates (which require respiration) and YPAD agar plates to confirm petite phenotypes.

### Fitness measurements and correlates

The fitness of the metal-adapted lines was assayed in YPAD and in each of the metal environments at concentrations determined to be lethal for the ancestor. We have observed that yeast adapted to a certain concentration of metal in the 96-deepwell plates (1 mL per well) are not able to grow at the same concentration in the Bioscreen C plates (130 µL per well), as also seen by Gerstein et al.^11^, potentially due to differences in aeration. In order to determine lethal concentrations in the Bioscreen C plates, we randomly chose three adapted lines for each metal and tested them in five concentrations of that metal spanning the range between the last concentration of high growth in the tolerance profile (Fig. S2) and the concentration used in the evolutionary rescue experiments. The highest concentration that allowed growth in all three adapted lines was chosen for use in fitness measurement assays (Table S1). We then measured maximum growth rate of adaptive mutations in the Bioscreen C for each metal in all the other metals, yielding a data-set that allows measurement of the pleiotropic effects of adaptive mutations to different metal environments *in vivo*. Relative growth rate of each metal-adapted line in a given environment was calculated by taking the difference between the maximum growth rate of the evolved line and that of the ancestral strain.

For a pair of metals, we define a narrow-scale cross-tolerance given by CT_(X,Y)_ = (r_Y|X_/r_X|X_ + r_X|Y_/r_Y|Y_)/2, where r_Y|X_ is growth rate assayed in metal Y given an evolutionary history in X. Broad-scale cross-tolerance CT_(X)_ was calculated by averaging the relative growth rate of a line across all metal environments Y except that line’s evolutionary environment X. If a genotype performs well, on average, in other environments, we consider it to be highly cross-tolerant and a generalist. Conversely, if it performs poorly, it is cross-intolerant and a specialist.

### Testing predictors of cross-tolerance

We expected that narrow-scale cross-tolerance CT_(X,Y)_ would be higher for pairs of metal environments that are more similar. We thus developed predictions for cross-tolerance using three different measures of similarity between pairs of metals, testing these predictions of CT_(X,Y)_ using regression models in R.

#### Prediction I: Similarity in reactivity of metals predicts cross-tolerance

Metal ions have different oxidizing and reducing potentials in solution. We first quantified the amount of available ions in the filtered metal media by Inductively Coupled Plasma Optical Emission spectroscopy (ICP-OES) analysis. Five mL of each metal media used for the experiment was filtered with a 0.22 µm filter and preserved with 2 drops of TraceMetal Grade Nitric Acid.

Oxidation-reduction potential (Eh) values quantify the potential of a medium to transfer electrons^24^. We used this as a proxy for the physicochemical impact of each metal. Redox readings were recorded with a YSI Professional Series Instrument Pro Plus sonde equipped with Ag/AgCl reference electrodes calibrated by Zobell solution 4M KCl. We converted the ORP readings into Eh values by adding the offset voltage of 200mV. We then used the absolute difference between the Eh values of metal pairs as a prediction of the amount of cross-tolerance.

#### Prediction II: Similarity in the sets of genes known to affect metal sensitivity predicts cross-tolerance

In this analysis we predict pleiotropy from gene annotations in the *Saccharomyces* Genome Database (SGD)^22^. A database query for the observable trait “metal resistance” yields a table of annotations from classical genetic studies and systematic mutation sets describing the effects of different mutations on the fitness of yeast in different metals. The data was manipulated in R to obtain a list of genes altering resistance for each metal.

SGD also includes information about the sign of the mutational effect. The original SGD data was filtered such that each row contained a gene, its mutation information (null or overexpression), the metal associated, and its phenotype (increased or decreased resistance to that metal). Each row was assigned a “gene activity” of – or + depending on if the mutation was described as null (knockout or loss of function) or overexpression, respectively. Each row was assigned a “direction of change” of + or – depending on if increased or decreased resistance resulted, respectively. Each row was assigned a “sign product” of + if the product of these two signs was positive (i.e., gene activity and direction of change are the same) or – if the product was negative. If the sign product of a mutation was the same for a pair of metals, then previously studied mutations have similar phenotypic effects in the two metals (positive pleiotropy). If the sign product differs, known mutations tend to have opposite effects when exposed to the two metals (negative pleiotropy). The proportion of positive pleiotropy was calculated by dividing the number of mutations with positive pleiotropy by the total number of common genes (i.e., genes with data for both metals, to control for different numbers of studies with the different metals). We expect a positive relationship between the proportion of genes exhibiting positive pleiotropy for two metals in SGD and the cross-tolerance C_(X|Y)_ for these two metals.

#### Prediction III: Similarity in metal abundance across environments predicts cross-tolerance

To measure environmental correlations between metal pairs, we downloaded data from the United States Geological Survey (USGS), specifically, the National Geochemical Database (https://mrdata.usgs.gov/#, accessed September 1, 2021, file: geochem, based on soil and sediment samples). From the whole database we retained only the metals used in our experiments. We extracted six columns (CD_ICP40, CO_ICP40, CU_ICP40, MN_ICP40, NI_ICP40, ZN_ICP40) reporting the metal concentrations in parts per million (ppm) as measured by Inductively Coupled Plasma (https://mrdata.usgs.gov/#geochemistry). We consider these data to represent a plausible set of environments experienced over the full evolutionary history of yeast (not the specific habitats of *S. cerevisiae*).

We converted all negative values to zeros, as these are considered measurements below detection levels from spectrometry after acid dissolution (USGS, further documentation is at http://mrdata.usgs.gov/geochem/doc/analysis.htm). We then added a value of one to all data points (to avoid zeroes) and log-transformed the data. Because metal pairs that co-occur often in the environment would more frequently have induced selection for mechanisms that confer tolerance to both metals in their evolutionary past^25^, we predicted that cross-tolerance would be observed more often for metals that positively covary in the environment.

### DNA extractions, library prep and sequencing

We extracted DNA and sequenced the genomes of 110 yeast lines, One-hundred and nine experimentally evolved, metal adapted yeast lines (16 cadmium-evolved, 16 cobalt-evolved, 15 copper-evolved, 22 manganese-evolved, 17 nickel-evolved, and 23 zinc-evolved) plus their ancestral strain were taken from freezer stock, plated on YPAD to isolate a single cell and cultured overnight in liquid YPAD media in a 30°C shaking incubator. Cells were then rinsed and resuspended in 1 M sorbitol. Samples were incubated in a solution composed of 1 M sorbitol, 0.5 M EDTA, and Zymolyase (5U/µL, Zymo Research E1005) to break the cell walls, followed by incubation in a buffer (1 M Tris-HCl, 5 M NaCl, 10% SDS, 0.5 M EDTA, 100% Triton-X and double-distilled water) with Proteinase K (20mg/mL, Luna Nanotech GPK-020) to lyse the cells. Samples were purified with phenol-chloroform-isoamyl alcohol (25:24:1, Sigma-Aldrich 77617-100), followed by precipitation using 100% isopropanol and washing with 70% ethanol. DNA pellets were air dried and then rehydrated in nuclease-free water. After the pellets had fully dissolved, samples were then treated with RNAse (10 mg/mL, Thermo Scientific FEREN0531) to remove any RNA contamination. Genomic DNA samples were further purified with silica columns and a binding buffer (guanidine-HCl, Tween-20, isopropanol, double-distilled water) followed by elution with nuclease-free water. The Qubit DsDNA Broad Range Assay Kit (Invitrogen Q32853) was used for quantification, following the standard protocol. Whole genomes were sequenced with Illumina technology on a NextSeq 550 System, 150-bp paired-end performed at the Sequencing + Bioinformatics Consortium at the University of British Columbia, Vancouver, British Columbia, Canada. The raw reads are available from the National Center for Biotechnology Information Short Read Archive (project number PRJNA994147).

### Sequence data analysis, variant calling and chromosome duplication

Raw reads were demultiplexed with bcl2fastq v2.20.0.422, without any modification and then cleaned with cutadapt using default settings for read quality and adapter removal. Paired-end reads were mapped to the *Saccharomyces cerevisiae* reference genome S288C in BWA^26^. Each mapped genome was then converted to a sam file, sorted, converted to a BAM file, and finally indexed in SAMTools^27^. We marked duplicate reads, estimated genome coverage, called variants for each individual and then combined all the variants in the experiment with *GATK4*^28^ (see S1 Appendix and GitHub repository for code). We focused on variant calls assuming haploid genomes, but we also called variants assuming diploid genomes. Depth of coverage was high for each sample (mean depth per site of 30.1 across all samples, with a minimum of 16.5, Table S2).

We extracted, cleaned and filtered genomic SNP information from the VCF in R using the tidyverse package^29^. We removed sites based on the distributions of quality measures, keeping only those with QD > 4, FS < 50, SOR < 4, MQ > 50. We then removed multiple alternates by removing anything with a “,” in “ALT”. For each metal sample, the genotype call was set to “.” whenever the depth of coverage was less five (DB<5). We removed any sites where all lines had the same genotype (ancestral to W303). We used SnpEff^30^ with the snpEff_v5_0_Saccharomyces_cerevisiae database to determine the effect of each mutation; we then removed non-protein coding mutations (variants considered as “MODIFIER”) and dubious genes. When there were multiple overlapping coding genes, we report the effect in the verified gene (no case of a mutation in two verified overlapping genes occurred). Sites with unidentified genotypes (“.”) for more than 5 of the 110 sampled sequences were removed; these often clustered and were consistent with alignment difficulties caused by divergent gene copies (*BSC1*, *DAN4*, *FLO1, FLO9, HAP1, MSS11, PIR3*, YHL008C, YKR073C), including flocculation genes known to exhibit copy number variation. We then re-called the variants assuming the samples were diploid and removed those sites with only heterozygous calls for the alternate allele (likely errors in alignment). A total of 414 unique mutations were observed, the majority in only one or two lines (384 and 18, respectively). Mutations in five genes were found in a large number of lines (Table S3B): 10 lines for a mutation in *NGG1*, 6 in *TFG1*, 8 in YLL066W-B, 23 in YGR130C, and 8, 8, 17, and 9 lines at four sites in YMR317W). Of these, YMR317W and YGR130C had nearly double the depth of coverage, consistent with a gene duplication, while YMR317W and YLL066W-B had mapping qualities (MQ) in the bottom 2% of genotype calls, consistent with alignment issues. These genes were removed from the data set (YMR317W, YGR130C, YLL066W-B). The SNPs at *NGG1* and *TFG1* were found together in multiple copper lines, suggesting contamination (discussed below). These sites were not removed from the analysis but were only counted once. After filtering, we confirmed the validity of each mutation by viewing the read alignments of the BAM files in IGV.

We calculated total depth of coverage for each chromosome to estimate and detect chromosomal duplication. We first created a coverage file for each site in each yeast line using the SAMTools pileup program. The files were then processed with a custom perl script, which adds together the depth of coverage in windows of 1000 bp. We used the same script to estimate the coverage for the mitochondrial genome of the petite lines in windows of 100 bp.

As *CUP1* duplicates fail to align, we estimated a relative copy-number of the *CUP1* genes by following the “*in silico*” protocol^11^. The protocol uses the unix command “grep” to count the instances of *CUP1* sequences directly in the unaligned fastq files, compared to reference loci (*RIX1*, *DED81*, *DUR3*) also on chromosome VIII. We report estimates of *CUP1* compared to the estimated copy-number of our sequenced ancestral line W303 (18.1 copies relative to the three reference loci).

We summarized the underlying genetic mechanisms for metal tolerance by running a GO term analysis for biological processes, molecular functions, and cellular components. We used the GO Slim Mapper tool (https://www.yeastgenome.org/goSlimMapper) in SGD (see Table S4 panel A for GO Terms used in analyses downstream). To formally test overrepresentation of GO terms, we used the GO Term Finder tool in SGD for all evolved lines except MnBM42 (102 SNPs) and MnBM14 (36 SNPs), which both had unusually high numbers of SNPs possibly due to mutations in DNA repair genes (e.g., *IRC20*, *FYV6*) and DNA damage response genes (e.g., *RAD17*, *MKT1*). Briefly, GO Term Finder compares the frequency of GO annotations in a given set of genes to those in the entire genome, and calculates a p-value using an algorithm based on a hypergeometric distribution (detailed description in Boyle et al.^31^). GO Term Finder performs multiple test correction by calculating a corrected p-value using the Bonferroni method and reporting a False Discovery Rate. The output includes a visualization of the GO term hierarchy, which was used to structure the terms presented in Figure 4. To investigate if certain GO terms were associated with broad cross tolerance, we ran parallel tests with GO Term Finder using gene lists from the lines with broad cross tolerance values in the top 25% and bottom 25%. We used the same methods to investigate if certain GO terms were associated with evolution in each environment using gene lists from the lines evolved in a given metal.

We used SnpEff^30^ to categorize each genic mutation as a “LOW” impact (synonymous), “MEDIUM” impact (non-synonymous), or “HIGH” impact (stop gained or frameshift). Using chi-squared goodness of fit tests, we compared the distribution of these effects in the whole mutation set to the distribution of effects in each significantly overrepresented term, allowing us to observe, for example, terms with significantly more loss of function mutations than expected (Fig. S5). A term was considered redundant and removed if its gene list had greater than 90% overlap with another term’s gene list. In these cases, the term with a lower FDR was retained, unless there were multiple cases of overlap with a term, in which case judgement was made to maximize representation across the GO term hierarchical tree.

### Parallel adaptation

Adaptation to a given heavy metal often involved the same gene (S2 Appendix). We performed randomisations using the logic of Gerstein et al.^11^ to calculate the probability of seeing so many genes hit multiple times, given the total number of mutations observed in each metal and the 6607 genes in the yeast genome. Specifically, we randomized the observed number of mutations across genes and compared the mean number of mutations per gene bearing a mutation to the means from 1000 randomized datasets (see the Supplementary S2 Appendix *Mathematica* file for detailed methodology). To be conservative, within each test, identical mutations appearing in multiple lines were ignored, as we cannot exclude some amount of cross-contamination. These calculations allowed us to test if there was significant parallel use of the same gene(s) during evolutionary rescue in each metal and across all of the metals combined.

## Results

### Evolutionary rescue, fitness profiles and cross-tolerance

Our goal was to isolate adaptive mutations to each metal by minimizing the time over which evolutionary rescue could occur, thereby avoiding the accumulation of non-adaptive mutations that would obscure the results (e.g., confusing linkage effects with cross-tolerance effects). On average, the lines described in this experiment were detected and isolated after 9.3 days of exposure to high metal concentrations (*SD* = 2.9).

Out of 1,440 total populations (240 replicates × 6 metals), rescue was observed a total of 152 times. Of these, 120 evolved lines were confirmed to grow in the presence of their evolution metal (Fig. 1 and Table S1) and were subject to further experimentation (21 cadmium, 16 cobalt, 15 copper, 22 manganese, 17 nickel, and 29 zinc lines). The average rate of rescue was 8% (*SD* = 2%) and was similar across metal environments.

Fitness assays (as measured by maximal growth rate) of all evolved lines and the ancestor in different environments reveal complex patterns (Fig. 2 and Fig. S3). Figure 2A-F shows the difference in growth rate of each evolved line relative to the ancestor when tested in each metal environment and ancestral conditions (YPAD). When assayed in permissive media (YPAD), a large proportion of metal-evolved lines (98 of 120, 82%) had significantly lower growth rate (Fisher’s exact test; P < 0.05) than the ancestor (left-most points in Fig. 2A-F), indicating the existence of trade-offs between tolerance to high versus standard metal concentrations. In general, replicate populations evolved in one environment have distinct patterns in their cross-tolerance to other metals, as demonstrated by the criss-crossing lines of the fitness profiles, possibly pointing to diverse adaptive mechanisms allowing for evolutionary rescue. A notable exception can be seen in the copper-evolved lines where many profiles have a similar shape.

**Figure 2.**
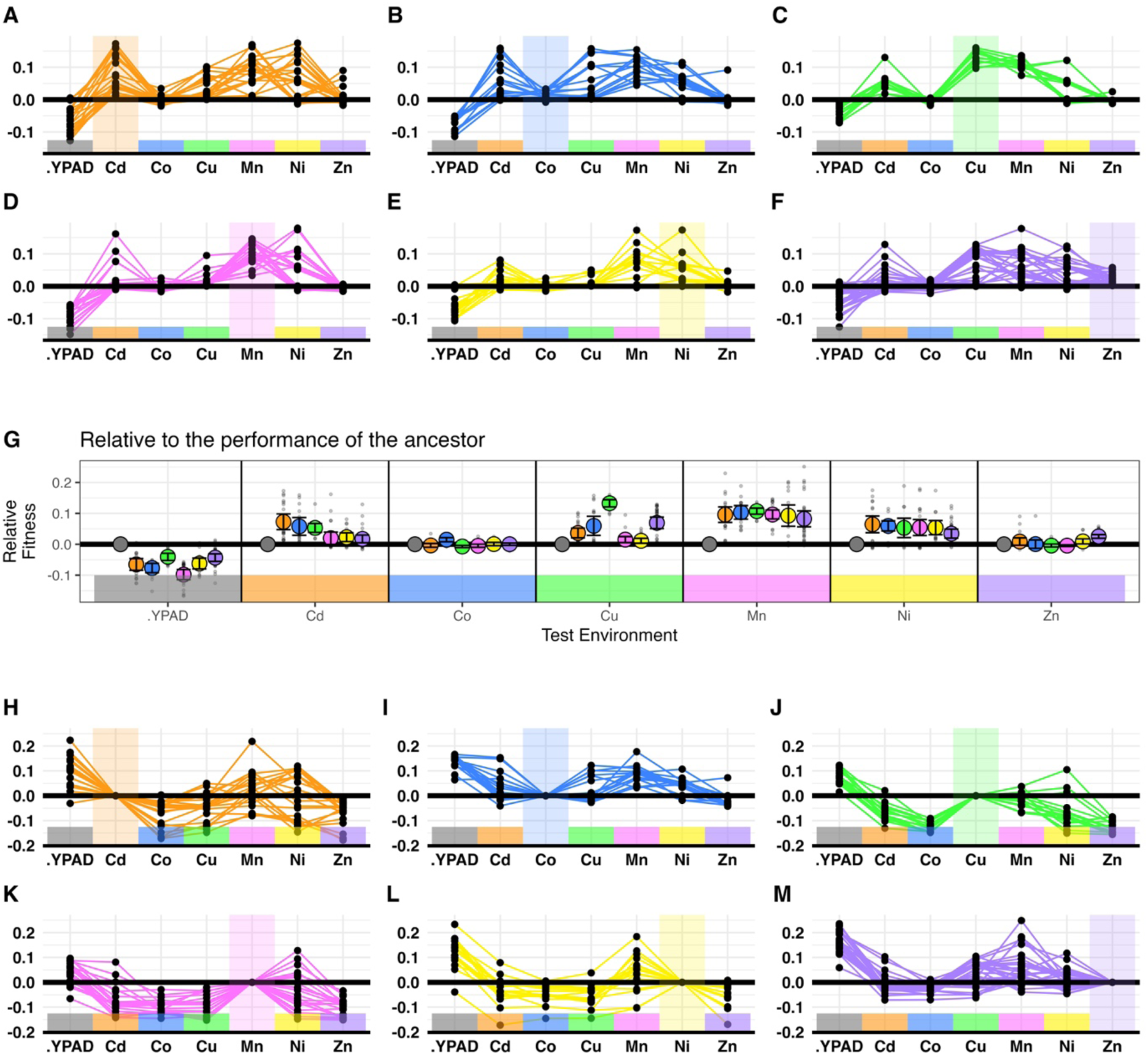
Relative growth rates of evolved lines tested in different metal environments. Panels show the growth rates of rescued lines relative to ancestral lines when evolved in (A) cadmium, (B) cobalt, (C) copper, (D) manganese, (E) nickel, (F) zinc. The colored lines connect dots representing a single evolved line assayed in different test environments, with shading highlighting the lines’ performance in their evolution environment. (G) The same data is rearranged to show the relative growth of evolved lines grouped by test environment (mean +/- SE). Panels (H-M) show the growth of rescued lines relative to their performance in the home environment; (H) cadmium, (I) cobalt, (J) copper, (K) manganese, (L) nickel, (M) zinc. The colored lines connect dots representing a single evolved line assayed in different test environments, with shading highlighting the lines’ performance in their evolution environment.

Figure 2G displays the average growth rate of lines evolved in each environment relative to the ancestor. In general, with the exception of manganese- and nickel-evolved lines, populations that had evolved in a particular metal performed better in that environment than those evolved in another metal. Regardless of evolution environment, metal-evolved lines commonly demonstrated cross-tolerance in manganese and nickel environments (Tukey’s HSD; p<0.05 for 6/6 mean comparisons of evolved-ancestor in Mn; p<0.05 for 5/6 mean comparisons of evolved-ancestor in Ni), whereas very few demonstrated cross-tolerance in cobalt and zinc (Tukey’s HSD; p<0.05 for 1/6 mean comparisons of evolved-ancestor in Co; p<0.05 for 1/6 mean comparisons of evolved-ancestor in Zn).

Figure 2 H-M display average growth rate of evolved lines assayed in a given metal, relative to the average growth rate of lines that had evolved in that metal. This alternative presentation highlights that some metal-adapted lines performed well across environments (generalists), such as cobalt and zinc adapted lines. By contrast, other metal-adapted lines were more specialized, performing well primarily in the environment to which they had evolved, such as copper and manganese adapted lines. When tested in permissive conditions (YPAD), generalist lines tended to grow better than specialist lines.

### Predicting cross-tolerance

We expected that cross-tolerance would be greater for metals that were more similar. We proposed and tested three measures of metal similarity: I. similarity in metal reactivity, II. overlap of known metal-resistance genes in SGD, III. correlation of abundance in environmental samples. In no case were these predictors significantly associated with narrow-scale CT_(X,Y)_, when tested with a regression analysis (Fig. 3).

**Figure 3.**
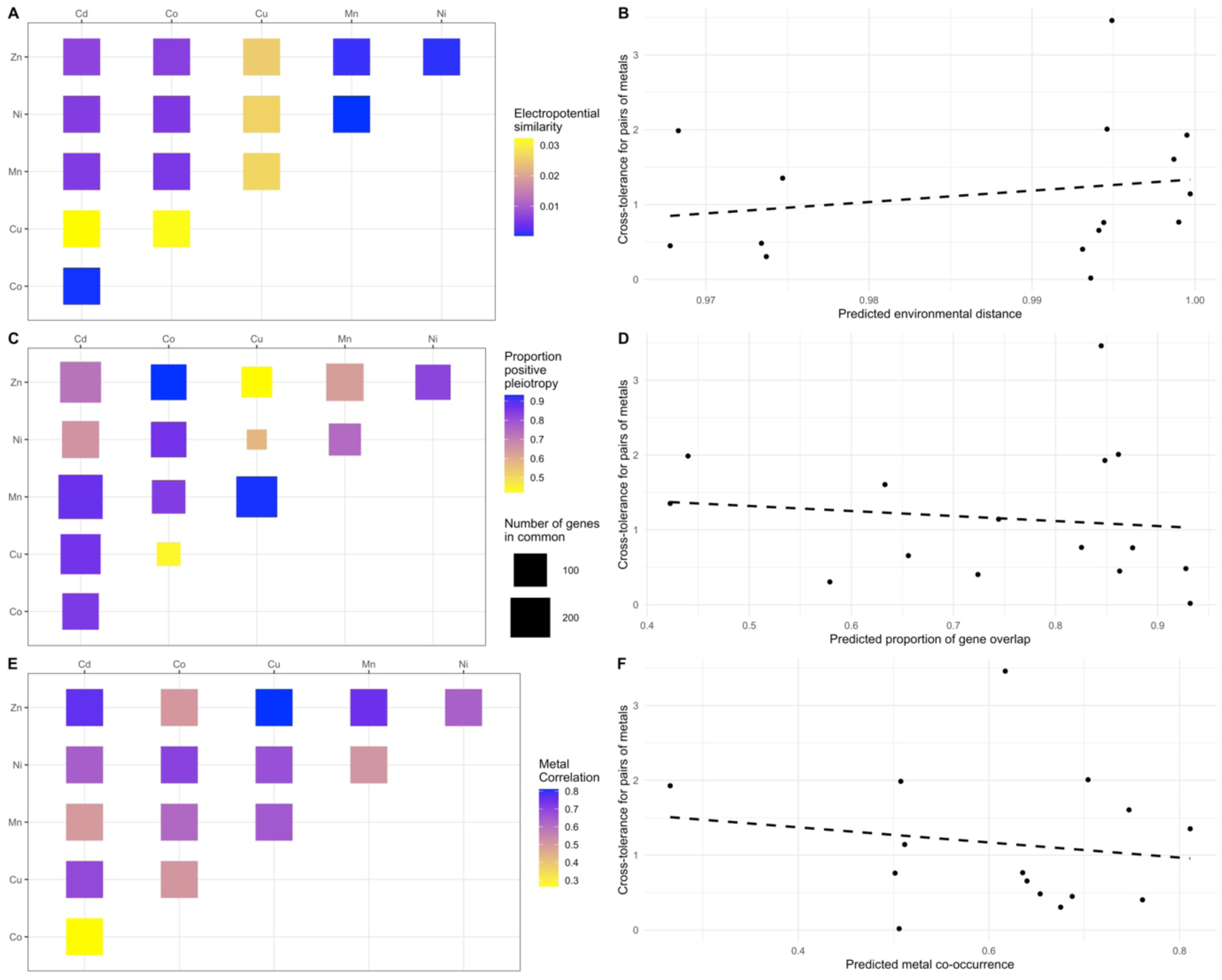
Predictions for cross-tolerance to metals. (A) Heat-map of the similarity in Oxidative-Reductive Potential (ORP) for different metal media. (B) Linear model of ORP similarity scores against narrow-scale cross-tolerance (CT(Y|X)). (C) Heat-map of gene overlap (<λ) from the SGD knockout and over-expression libraries. (D) Linear model of gene overlap (<λ) against narrow-scale cross-tolerance (CT_(Y|X)_). (E) Heat-map of the co-occurrence of metals in soils from the US Geological Survey. (F) Linear model of soil metal correlations against narrow-scale cross-tolerance (CT_(Y|X)_).

These similarity indices do not account for the order of exposure, such that lines evolved in metal A and tested in metal B are predicted to have the same amount of cross-tolerance as lines evolved in metal B and tested in metal A. According to our results, cross-tolerance is not generated reciprocally for metal pairs. For example, the copper-adapted lines performed well in manganese, however manganese-adapted lines performed poorly in copper (Fig. 2G). Similarly, cadmium-adapted lines performed well in nickel, but nickel-adapted lines did not show enhanced performance in cadmium.

Thus, even with similar metal stressors, the evolution of cross-tolerance is idiosyncratic and depends on the specific mutations that arose rescuing the yeast lines (evolutionary stochasticity).

### Genomic changes underlying cross-tolerance

We sequenced 110 genomes from this experiment (109 evolutionary rescue lines and 1 ancestral strain; 11 lines did not yield enough cells for WGS). After filtering out poor quality genotype calls (see methods), we recorded several types of genomic mutations (Table S2). Across all evolved lines, we found a total of 450 SNPs and indels that passed our quality filters, falling in 331 genic regions (Table S3).

Two lines had an unusually high number of SNPs (MnBM42 with 102 and MnBM14 with 36, see Fig. S4). Manganese is known to be mutagenic^32,33^, and both lines exhibit mutations in DNA repair genes (e.g., *FYV6*, *MKT1*, *RAD17*, and *IRC20*). These two lines are thus potentially interesting examples of mutator phenotypes evolving in the context of an evolutionary rescue experiment, and we leave their investigation to a future study. Excluding these two lines from further analyses, we found a total of 312 high-quality SNPs occurring in 210 genes (Table S3). On average, the remaining evolved lines had 3 mutations in genic regions, though this varies significantly among lines evolved in different environments (ANOVA, p=3×10^-9^; a mean number of genic mutations [SE] of 3.63 [0.59] in Cd, 5.38 [0.5] in Co, 2.33 [0.4] in Cu, 3.55 [0.62] in Mn, 1.41 [0.32] in Ni, and 1.65 [0.2] in Zn).

For most of the 109 sequenced lines, we found at least one mutation that could account for the increased metal tolerance based on information in SGD for each gene (as detailed below and in Table S2). There were seven lines for which we could not confidently detect any genetic change despite high average sequence coverage: CuBM8, CuBM16, NiBM22, NiBM25, CdBM36, ZnBM25, and ZnBM34. These lines may harbor mutations that we did not investigate, including regulatory changes or structural changes beyond those tested (aneuploidy and *CUP1* coverage). Alternatively, epigenetic changes may have been involved.

Analysis of the genes mutated in the rescued lines revealed significant overrepresentation of several GO terms (Fig. 4 and Table S4B) including: ‘fungal-type vacuole membrane’, ‘incipient cellular bud site’, and ‘vacuolar transporter chaperone complex’ among cellular components; ‘response to stimulus’, ‘biological regulation’, ‘cytokinetic process’, and ‘polyphosphate biosynthetic process’ among biological processes; and ‘phosphotransferase activity’, ‘ubiquitin-protein transferase activity, and ‘MAP-kinase scaffold activity’ among molecular functions. Comparison of significantly overrepresented terms in the most and least cross-tolerant lines show that specialist lines tend to have more mutations affecting the plasma membrane, while generalist lines benefit from changes to ubiquitin ligase and transferase activity (Table S5). Analysis of genes mutated in each evolution environment suggest that different metals target different cellular functions. For example, the vacuolar transporter chaperone complex is targeted in cobalt, incipient cellular bud site in manganese, symporters in nickel, and cell signalling in zinc (Table S6).

**Figure 4.**
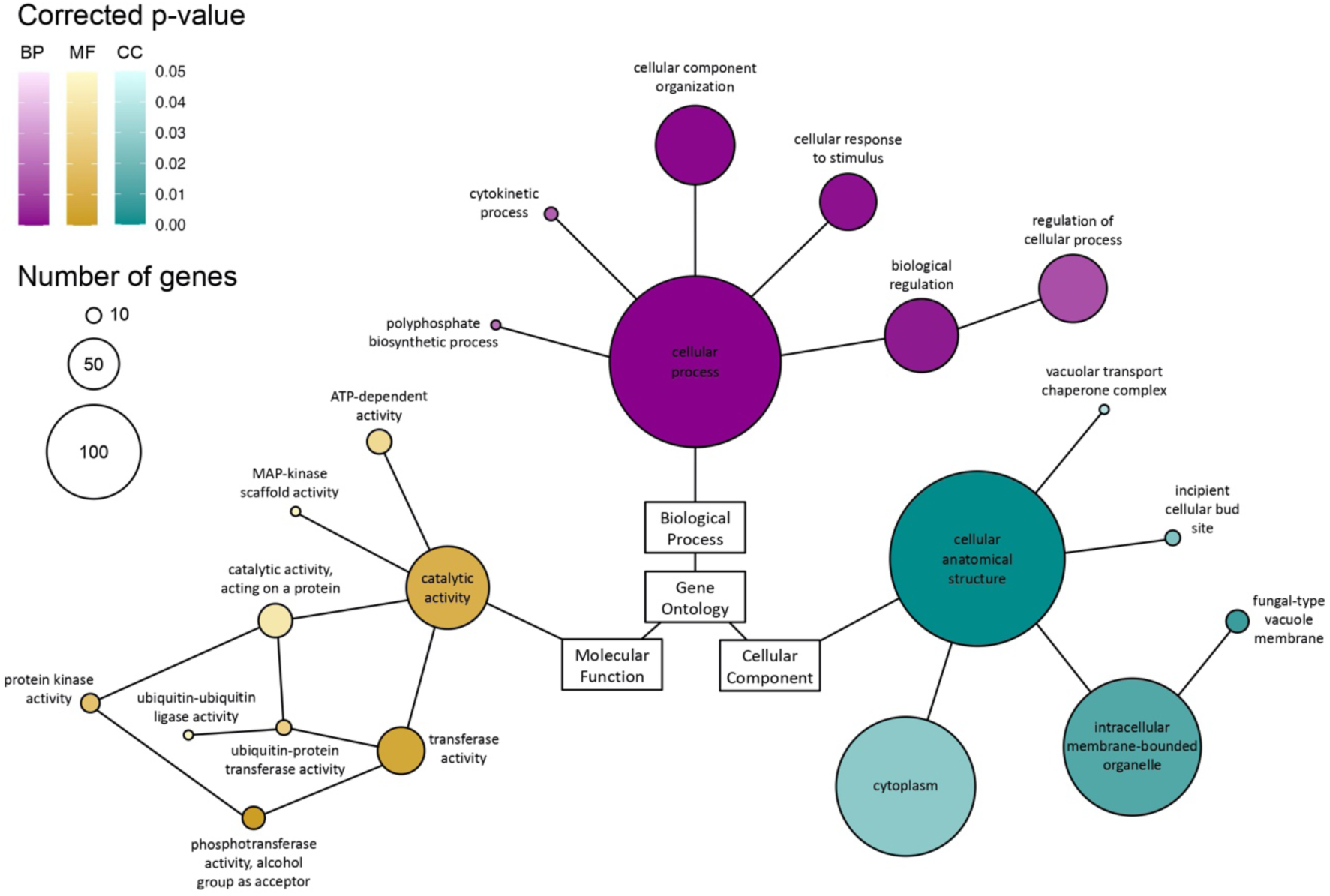
Hierarchical tree showing relation of significantly overrepresented GO terms found in GO Term Finder. All terms on this tree have at least 3 genes involved and a Bonferroni-corrected p<0.05. The size of the circle represents the number of genes in our mutation list that are associated with that term. See Table S4(B) for precise values.

Concordant with findings by Ruotolo et al.^34^, the vacuole was of significant importance in the adaptive response for metal detoxification. The genes associated with the vacuole and vacuolar transporter chaperone complex had significantly more loss of function mutations than expected, *X*^2^ (2, *N* = 37) = 18.033, p = .00012 and *X*^2^ (2, *N* = 10) = 25.899, p = 2.38E-06, respectively (Table S7).

We recorded 40 genes with mutations occurring in multiple lines, 24 of which involved different SNPs. Mutations within the same gene did not always exhibit the same cross-tolerance phenotype. For example, CuBM3, ZnBM29, and ZnBM37 all had a single genic SNP, occurring in *PMA1*, but these lines displayed moderate, high, and low maximal growth rates in manganese, respectively (Fig. 5).

**Figure 5.**
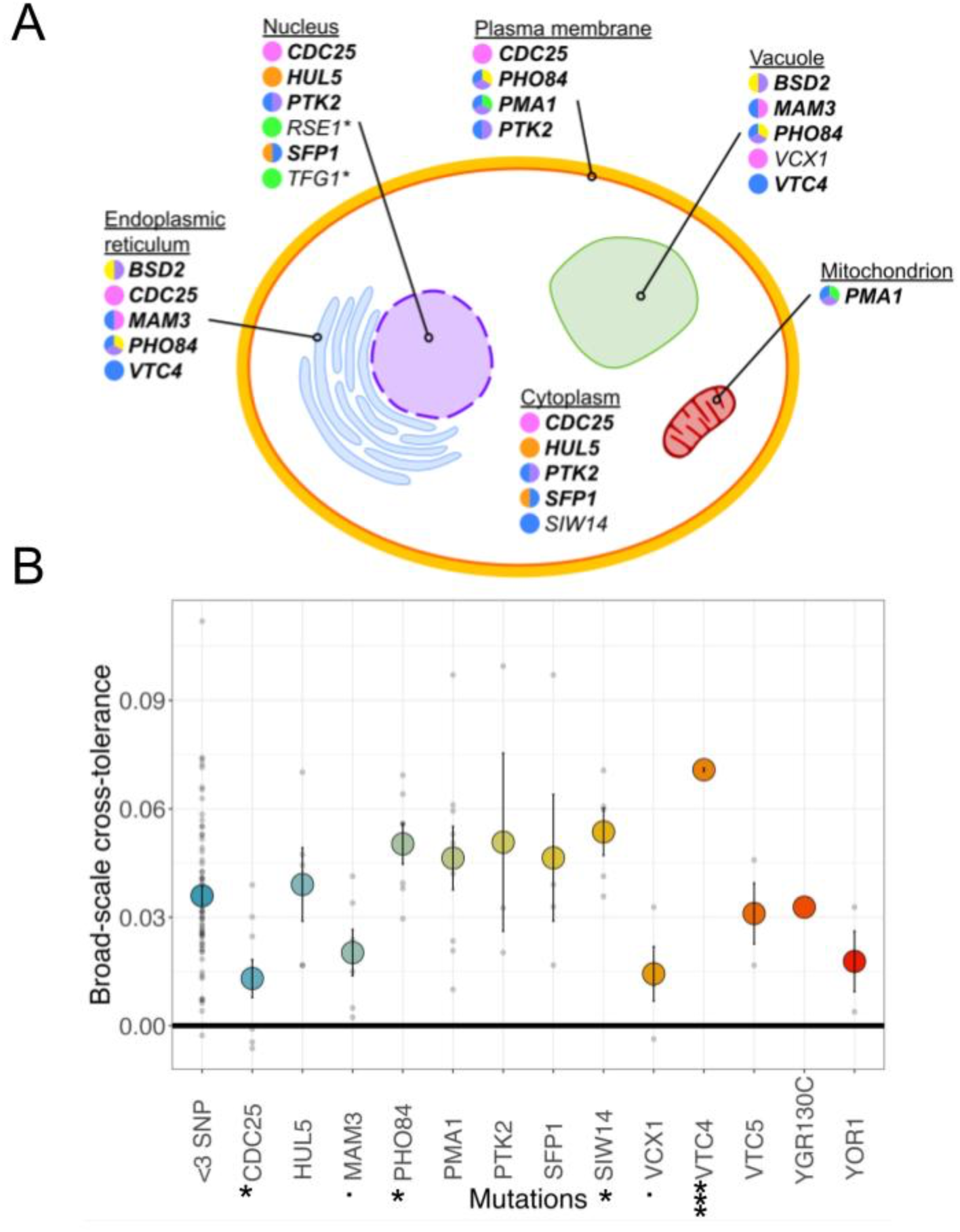
Localization and broad-scale cross-tolerance of genes involved in metal tolerance showing parallelism. (A) Cellular localization of genes associated with SNPs. The color (Cd = orange, Co = blue, Cu = green, Mn = pink, Ni = yellow, Zn = lilac) of the pie-charts indicate the evolution environment of the mutation. (B) Broad-scale cross-tolerance comparison of lines with mutations in genes showing a high degree of parallelism (≥3 independent mutations, Table S8) to the remaining lines (mutations only in genes hit by fewer than 3 SNPs). Error bars ±SE. Asterisk (*) indicates genes significantly different (t-test) from the remaining lines (bearing mutations only in genes hit <3 times) with a p-value <0.05, whereas a dot (.) indicates marginal significance (0.05 ≤ p ≤ 0.07). Three asterisks (***) indicate a p-value much smaller than 0.0001.

We next tested whether parallel mutations involving the same genes occurred more often than expected by chance in each metal, excluding identical base pair changes to be conservative. For copper, no gene carried SNP mutations in multiple lines, suggesting no parallelism, but the copper metallothionein gene, *CUP1*, showed parallel amplification in many of these lines (see below). For every other metal, significantly more genes bore SNP mutations in multiple lines than expected, with the mean number of mutations observed at mutated genes being higher than all 1000 randomizations (Fig. 5B and S2-3 Appendix). Similarly, considering all metals together, the mean number of mutations per gene bearing a mutation was higher than all 1000 randomizations. This was true even when we counted a maximum of one mutation from each metal for a given gene, indicating that there was significant parallelism among lines adapted to different metals as well (S2-3 Appendix).

Three or more independent mutations were observed within 13 genes (including MnBM14 and MnBM42; nine excluding these lines; Table S8). Lines bearing mutations in these genes do not show significantly higher broad cross-tolerance, on average, than remaining lines, although some specific genes do (Fig. 5B; t-test, p=0.71). Notably, *VTC4*, a gene only mutated in cobalt-evolved lines, provides significantly broader cross-tolerance than genes showing no parallelism (t-test, p<0.0001). Additionally, *SIW14* mutations evolved only in cobalt and *PHO84* evolved in cobalt, nickel, and zinc also show a significantly higher broad cross-tolerance (t-test, p<0.05). Conversely, some genes (e.g., *MAM3, p=0.06*) do not provide broad cross tolerance despite arising frequently. Further, *CDC25*, a gene only mutated in manganese-evolved lines, provide significantly lower broad-scale cross-tolerance than even the genes showing no evidence of parallelism (t-test, p=0.002).

Some of the genes that repeatedly acquired mutations were also observed in the evolutionary rescue study of Gerstein et al.^11^ with copper. They observed multiple (≥2) mutations in four genes: *VTC1* and *VTC4*, whose proteins are constituents of the vacuolar transporter chaperone complex, *PMA1*, essential for proton transport across the plasma-membrane (a P2-type H+-ATPase), and *MAM3*, which plays a role in mitochondrial organization and vacuolar sequestration and whose protein localizes to the vacuolar membrane^22^. We also observed mutations in these four genes, but involving other metals (not copper), including four *VTC4* mutations in cobalt (two of the SNPs occurring in the same CoBM3 line), two *VTC1* mutation in cobalt, and five independent *MAM3* mutations in cobalt and manganese.

We further investigated the degree of parallel evolution observed in different metals (Table S8). Mutations accumulated in more than one metal at ten genes (*DNF1, FYV10, KSP1, MAM3, PDR1*, *PHO84, PMA1, PTK2, SFP1, VTC5*), with eight additional genes if we include the mutator lines MnBM14 and MnBM42 (*HUL5*, *LAM1, PPQ1*, *SIW14*, *SNT2*, *TOM1*, *YDL199C*, *YOR1*). This is a highly significant degree of parallelism, even counting parallel mutations accumulated in the same metal only once (S2-3 Appendix). Many of these mutations play a role in transport across membranes (Table S8). Of note, *PTK2* is required for the activation of *PMA1*^35^, *PHO84* produces a plasma membrane protein involved in transport of inorganic phosphates with some activity as a manganese transporter (we did not observe *PHO84* mutations in manganese but in cobalt, nickel and zinc).

### Direct mutagenic effects of metals

We observed a disproportionate amount of G:C → A:T transitions. We found that the mutational spectra for three of the six metals (cadmium, cobalt, and manganese) were significantly different from the mutational spectrum for yeast reported by Lynch et al.^36^ (Fig. S6). The two mutator lines evolved in manganese also exhibited significantly more G:C → A:T transitions. As in Gerstein et al.^11^, we did not find the mutations accumulated in copper to deviate significantly from the expected spectrum, even though exposure to high concentrations of copper has been shown to have mutagenic effects on DNA replication in phage, leading to a preponderance of C → T transitions^37^.

Deletion mutations from this study were predominantly found in cadmium. Of the total deletions we identified (61), half were in cadmium (31), 13 in manganese (one of which was in MnBM42), eight in cobalt, six in zinc, three in nickel, and none in copper. Consistent with this result, *in vitro* studies of *S. cerevisiae*^16,38^ have shown cadmium to induce deletions via increased recombinagenic activity.

### Chromosome and *CUP1* amplification

We estimated whole-chromosome coverage and found chromosome duplications evolved in all metals except copper (Fig. S5A and S7 Fig). Whole-chromosome duplications convergently evolved in nickel, where four lines had duplications in chromosome XIII and XIV, and in cadmium, where three evolved lines had duplications in chromosome II. We were limited in our ability to test if any instance of chromosome duplication would enhance cross-tolerance. However, we tested copper-evolved lines with chromosome duplications from Gerstein et al.^11^ in the metals we used in our experiment. Gerstein et al.^11^, unlike us, found many whole-chromosome duplications in their copper evolved lines and tested for the statistical association of chromosome II duplication with copper tolerance but did not find a significant one. We also could not find a significant difference in growth rate between aneuploid and euploid lines in our experiment lines and the copper-evolved lines from Gerstein et al.^11^ when tested in any of the metals (see Fig. S8).

Copper lines showed the lowest number of SNP-based mutations, on average, and we found no support for significant parallelism among these SNP mutations. When we estimated the scaled coverage of the *CUP1* gene, a known chelator of copper ions, we found increased copy-number in 7 out of 15 copper lines (Fig. S5B, Table S9). Six of the seven lines with an increased number of copies of *CUP1* also had identical SNP mutations in gene *TFG1* (Fig. S8), the largest subunit of *TFIIF* (Transcription Factor II) facilitating both transcription initiation and elongation of RNA polymerase II^22^. This suggests that *CUP1* amplification in these lines might have occurred through contamination; *CUP1* amplification was nevertheless observed in one other well (CuBM15) of the same plate without the *TFG1* mutation (Fig. S9). We report estimates of *CUP1* compared to the estimated copy-number of our sequenced ancestral line W303 (18.1 copies relative to the three reference loci, which is similar to the 18.13 average copy number reported by Gerstein et al. in their BY4741 strains). Previous studies have found that *CUP1* amplification can increase tolerance to cadmium as well as copper^39,40^. We observed a slight increase in *CUP1* copy numbers compared to the ancestor in two cadmium-evolved lines (1.18 in CdBM45 and 1.16 in CdBM46), one zinc-evolved line (1.18 in ZnBM17), and two manganese-evolved lines (1.13 in MnBM17 and 1.15 in MnBM27). Additionally, copper-adapted lines were not particularly cross-tolerant to cadmium except for CuBM13, which did not exhibit *CUP1* amplification but mutations in *NGG1* and *RSE1*, a component of the pre-spliceosome, also involved in transport from the endoplasmic reticulum to the golgi (Fig. S2C).

### Mitochondrial function

Of the 120 metal-evolved lines in our experiment, 38 were unable to grow on glycerol-based media, indicating loss of mitochondrial function during evolutionary rescue. Oxidative phosphorylation in the mitochondria produces ROS intrinsically^41^, potentially contributing to oxidative load when the cell encounters exogenous oxidizers, such as heavy metals. The repeated loss of respiration observed in this experiment may not, however, be adaptive. Cobalt and manganese stress are known to induce the loss of mitochondrial function in yeast^33^. All the lines evolved in cobalt (16) evolved a petite phenotype, 17 out of the 22 in manganese, three in copper, and two in nickel (Fig. S6A). Using ten petite and ten grande lines from a different experiment^23^, we found no significant effect of the petite phenotype on metal tolerance (Fig. S6B).

We mapped the mitochondrial reads and estimated coverage across the mitochondrial genome. Cobalt evolved lines showed no trace of mitochondrial DNA. Manganese, copper, and nickel petite lines, however, showed extensive loss of mitochondrial genes but a remarkable increase in coverage of some breakpoint regions between respiratory genes (Fig. S6C). In the cases of partial loss with intermittent amplification, respiration genes were always eliminated (i.e., *COX1*, *COX2*, *COB*) ^42–44^, with high coverage in various non-coding regions. These fragmented mitochondrial genomes are consistent with previous studies that have found the petite phenotype can arise from recombination events between repeats of GC clusters and AT stretches, especially at highly homologous origins of replication^45,46^.

## Discussion

We assayed the cross-tolerance of lines evolved in one stressful metal to other metal environments by comparing growth rates across environments and whole-genome sequencing of 109 evolved yeast lines, adopting the methods of previous evolutionary rescue studies to detect first-step mutations^11,47^. We developed three *a priori* predictions for cross-tolerance based on the similarity between pairs of metal environments in their physiochemical properties, their genetic overlap, and their co-occurrence in the environment. None of these three measures predicted cross-tolerance values. Instead, the exact order of metals encountered (Fig. 2) and the specific mutations acquired played the largest role in the development of cross-tolerance, highlighting the importance of evolutionary stochasticity, as noted by Jerison et al.^2^ In making these predictions, we assumed that mutations evolved in each metal would have similar cross-tolerance for the other metal within a pair. Instead, our observations show that cross-tolerance does not evolve reciprocally, again highlighting the importance of stochasticity. Even yeast populations evolving in the same stressful environment can accumulate different mutations that confer more or less cross-tolerance^2,48^ (see criss-crossing lines in Fig. 2, e.g., for manganese-adapted lines in panel D).

We expected that the genes involved in metal tolerance would span the many strategies organisms evolve to cope with metal stress such as transporters or increased production of antioxidants^10,12,14^. Indeed, several observed mutations occurred in transporters previously reported to be important in metal adaptation: *PMA1*^49–52^, *PHO84*^53–55^, *PTK2*^56^, *VCX1*^57^. We also found mutations in genes that help with oxidative stress. For example, *SNT2* regulates gene expression in response to oxidative stress from peroxide (mutated in a zinc line and MnBM42, Table S3), as does the cytosolic quality control protein encoded by *HUL5*^58^ (observed in four cadmium lines and MnBM42, Table S3).

Many of the overrepresented GO terms (e.g., phosphotransferase activity, kinase activity, MAP-kinase scaffold activity, response to stimulus, biological regulation) include genes whose products contribute to the stress response system through highly conserved signalling networks like the PKA, MAPK, and TOR pathways^59^. These pathways regulate cell growth in response to environmental and cellular cues (e.g., oxidative/genotoxic/heat stress), allowing resources to be diverted from growth and reproduction toward damage repair and maintenance^60^. Loss of function in central components of these pathways (e.g. *SFP1*^61^, *CDC25*^62^, *PBS2*^63^, etc.) may prevent transduction of signals that halt the cell cycle in toxic metal environments, allowing for continued reproduction.

The stress response system also regulates autophagy and the ubiquitin-proteasome system (UPS), both of which target damaged cellular components for degradation. While both of these functions are repeatedly targeted in our study (significantly for UPS, but not autophagy), the mutations do not imply a favoured direction for degradation of cellular components in metal environments. For example, the TOR and PKA pathways negatively regulate autophagy, and their inhibition induces robust autophagy activity^64–66^. Conversely, most of our mutations in the ubiquitin-proteasome system are predicted to decrease degradation of the targeted cellular components (i.e., due to their high impact, such as frameshift mutations, in *HUL5*, *BSD2*, *YLR108C* ^67^).

Changes to pathways controlling phosphate metabolism have the potential to provide cross-tolerance among multiple metals. A subset of mutations arose in multiple metals (Table S8), three of which (*PHO84, VTC4, SIW14*) conferred broader cross-tolerance. These genes are all involved in phosphate metabolism and homeostasis via the PHO pathway. The PHO pathway is repressed under high inorganic phosphate (Pi) levels, leading to degradation of the Pi transporter Pho84p^68^. Pho84p also transports metal cations (e.g., Mn²⁺, Cu²⁺, Co²⁺, Zn²⁺), with loss of function linked to metal resistance and overexpression to metal accumulation^53,55^. Loss of *VTC4* and *SIW14* function may suppress *PHO84* expression or signal Pho84p degradation. The *VTC* complex synthesizes polyphosphate (polyP) while translocating it into vacuoles^69^, and reduced polyP synthesis increases cytosolic Pi, lowering Pho84p activity^70–72^. Siw14p is an inositol phosphatase which modifies inositol pyrophosphates (i.e. transforming IP_8_ into 1-IP_7_ and 5-IP_7_ into IP_6_), key signalling molecules of the PHO pathway^73^. Recent work suggests that the principal signal for activation of the PHO pathway is loss of IP_8 74_, so loss of function in *SIW14* may maintain IP_8_ levels such that the PHO pathway remains inactive.

Despite the idiosyncratic nature of the cross-tolerance we observed, two broad patterns are readily apparent. First, some metal environments tend to generate lines that are broadly cross-tolerant (generalists), while others generate lines that can only grow in that environment (specialists). Second, certain metal environments are broadly permissive for growth of lines evolved in other metals, while others are not.

The breadth of cross-tolerance demonstrated by an evolved line can be largely explained by the metal in which the line evolved. Cobalt-evolved lines exhibit the broadest cross-tolerance, likely due to their higher mutation count, increasing the likelihood of beneficial mutations across environments. In contrast, manganese-evolved lines show significantly less cross-tolerance. This may stem from manganese-specific adaptations, as Mn²⁺ complexed with orthophosphate (Pi) can scavenge reactive oxygen species (ROS)^75,76^, reducing ROS-driven selection pressure. Consequently, manganese adaptation may prioritize other metal toxicity mechanisms, such as disrupted signaling or inhibited protein biosynthesis^77^.When introduced into novel metal environments that induce a higher direct ROS burden, manganese-adapted yeast may be less protected. We also note that manganese-evolved yeast have lower growth rates in the ancestral environment (YPAD) compared to other metal-evolved lines suggesting that the mechanisms allowing adaptation to manganese may be more costly.

The cobalt test environment posed significant challenges for most metal-evolved lines, resulting in low average growth rates, even for cobalt-adapted lines. Lines that could grow typically had many mutations or mutations in genes linked to numerous GO terms (Table S10), suggesting cobalt’s widespread cellular impact requires extensive adaptation. In contrast, the manganese environment supported growth for most metal-evolved lines (Fig. 2H-M). Reduced ROS levels, potentially due to manganese-phosphate complexes^53^, may have benefited lines from other environments by alleviating the need to counter ROS, allowing them to focus resources on growth. However, it remains unclear whether these cross-tolerance differences stem from inherent metal properties or the specific concentrations used. While toxicity levels were calibrated to inhibit ancestral strain growth, some variability persisted.

Exposure to a stressful environment can increase genetic instability and mutation rates^78^. We found that cadmium, cobalt, and manganese of our six metals had mutational effects on the yeast nuclear genome, specifically with an increase in G:C → A:T transitions in all three and an increase in deletions in cadmium. Among these, two manganese-adapted lines (MnBM42 and MnBM14) evolved mutator phenotypes and accumulated a disproportionate number of G:C → A:T transitions. Thus, while evolutionary rescue typically involved the accumulation of few mutations, multi-step mutations also arose, likely due to the combined effects of manganese exposure and mutations in key error correction genes.

In addition to point mutations, additional strategies for metal adaptation include increased copy number of resistance genes, either through aneuploidies or single gene duplications. Whole chromosome duplications evolved in several metals (e.g. chromosome XIII in nickel and zinc, chromosome II in cadmium and zinc). While it is possible that having extra copies of certain genes may be beneficial in some circumstances, our evidence does not suggest that these chromosome duplications were adaptive in metal tolerance and may instead be a consequence of altered cell cycle mechanisms. Gerstein et al.^11^ found extensive duplications of whole chromosomes in their copper-adapted lines (particularly, chromosomes II and VIII) but also found no evidence of a beneficial effect of those mutations. Gorter et al.^52^ evolved yeast in cadmium and found whole genome duplications and increased copy number of chromosomes XIV, III and VI but a decreased copy-number of chromosome I. Given the oxidative nature of metal ions, and following previous studies on copper^11^, we expected to find amplification of chelator proteins that bind metals and render them bio-unavailable. We did indeed find increased copy-number of the *CUP1* gene, encoding for the copper binding protein, but this phenomenon only occurred in the copper environment. Despite previous work suggesting that cadmium might also be neutralized in this way, we found that copper lines with higher copy-number of *CUP1* did not exhibit higher performance in cadmium^11,43,79^.

In conclusion, we developed a series of evolutionary rescue lines with yeast exposed to different metals. These lines were then assayed for cross-tolerance across the six metals examined. We found no relationship between the three predictors of cross-tolerance: I. similarity in metal reactivity, II. overlap of known metal-resistance genes in SGD, III. correlation of abundance in environmental samples. Although not predicted, cross-tolerance was high for lines that had adapted to cobalt or zinc (Fig. 2I and 2M) and for lines bearing changes in *VTC4* (Fig. 5B), presumably due to their effect on Pho84p activity. Thus, some evolutionary paths were more likely to lead to general cross-tolerance, while others to narrow tolerance, depending on the series of environments encountered and the particular mutations that accumulated.

It remains an important challenge for evolutionary biologists to predict the combinations of stressors that will most often lead to extinction and those that allow evolutionary rescue. While the three predictions that we tested failed to account for cross-tolerance in different metals, these tests were limited by the information used to measure each predictor. For similarity in metal reactivity, incorporating other chemical properties of metal ions beyond redox potential using a multivariate approach or accounting for intracellular ROS production may have led to more accurate predictions. For overlap of known metal-resistance genes in SGD, focusing on genes with the strongest effect on metal tolerance might also have been more predictive, such as the subset of genes that we identified *post hoc* as involved in adaptations to multiple metals (Appendix S3). Finally, the correlation of metal abundance in environmental samples used all samples available from the USGS, but targeted sampling of habitats more relevant in the evolutionary history of *S. cerevisiae* might paint a clearer picture of the metal combinations that yeast have evolved to tolerate through common genetic pathways. Further work is needed to understand the evolutionary challenges imposed by combinations of stressors, as they occur in natural and polluted habitats. Yeast evolutionary experiments provide a promising avenue forward by revealing the correlated effects of adaptive mutations to different stressors.

## Acknowledgements

Thanks to Linnea Sandell, Matt Stasiuk, Lauren McBurnie, Sayantan Datta, Kian Mousakhan Bakhtiari for help in the lab. This research was supported by the NSERC Discovery Grant to SPO (RGPIN-2022-03726).

## Supplementary Tables

TS1 Metal concentrations for evolutionary rescue experiment and for reciprocal testing. Number of lines and number of petite phenotypes.

TS2 Genome statistics and types of mutations per metal adapted line.

TS3 SNP mutation annotation, position and and SnpEff output.

TS4 GO Terms found (A) and GO term over-representation analysis statistics by component, function and process (B).

TS5 GO term over-representation against broad-scale cross-tolerance statistics.

TS6 Overrepresentation of GO terms evolved in each metal environment, identified by GO Term Finder.

TS7 Chi-squared goodness of fit test on distribtuion of mutational effects

TS8 Genes with three or more mutations (SNPs and indels)

TS9 *In silico* estimates of *CUP1* copy-numbers

TS10 Top 10 growers in each environment - number of terms per gene.

## Supplementary Figure legends

S1 Experimental blocking design from 240 isogenic yeast colonies derived from the same ancestor.

S2 Range of ten concentrations of initial pilot study used to determine experimental treatment.

S3 Each evolved line performance profile.

S4 Natural log of mutations evolved per line (excluding zeroes).

S5 GO terms analysed by their SnpEff results showing how types of mutations are distributed across the GO terms.

S6 Mutational spectra by evolution metal against Lynch et al. and Gerstein et al (top row). Asterisk indicates mutational spectrum of single mutator lines (colored in pink because both were evolved in manganese).

S7 Graphs of coverage by chromosome used to estimate aneuploid evolved lines.

S8 Relative fitness of lines comparing euploid and aneuploid lines from the current study experiment (A) and lines tested in metals from Gerstein et al. (2015).

S9 Whole-chromosome duplications and *CUP1* amplification. (A) Whole-chromosome coverage estimates. Estimates are scaled to the ancestral (W303) to be able to compare across lines. (B) *CUP1* coverage, scaled to ancestral line (W303), see Table S2 for estimates compared to three control genes. Ancestral W303 line in grey had 18.1 estimated copies compared to three control genes, this estimate is similar to the 18.13 copies estimated in the ancestral line used by Gerstein et al. (2015) compared to the same three control genes.

S10 Distribution of copper mutations on deep-well experimental plate. Non-green circles indicate wells with no detected growth (extinct populations) and green circles indicate lines isolated for this study. Legend indicates types of mutations evolved in each well.

S11 Petite performance in metals and mitochondrial genome coverage. (A) Grande and petite performance in metals and YPD. (B) Grande and petite performance in manganese, cobalt and YPD from an outcrossing experiment using the same ancestral strain (W303). (C) Mitochondrial genome coverage for petite lines with some remnant mitochondrial genome calculated in 100bp windows. All cobalt evolved lines and two copper evolved lines had complete loss of mitochondrial genomic material and are not shown in this graph.

