## Supplemental Figures for "Evolution of cross-tolerance to metals in yeast"

S1

The ancestral strain W303 background is grown up in liquid media and plated on petri dishes to sample 240 single colonies.

240 single-colony ancestral populations in 3 (A, B, C) 96-well plates (180 per plate).
Yeast was grown up and then diluted to the same OD to have equivalent starting population sizes.

A, B, and C plates with 240 normalized populations are used to inoculate metal media. Yeast was left to grow for two weeks. When growth was observed in a well, the well was then sampled after 24 hours.

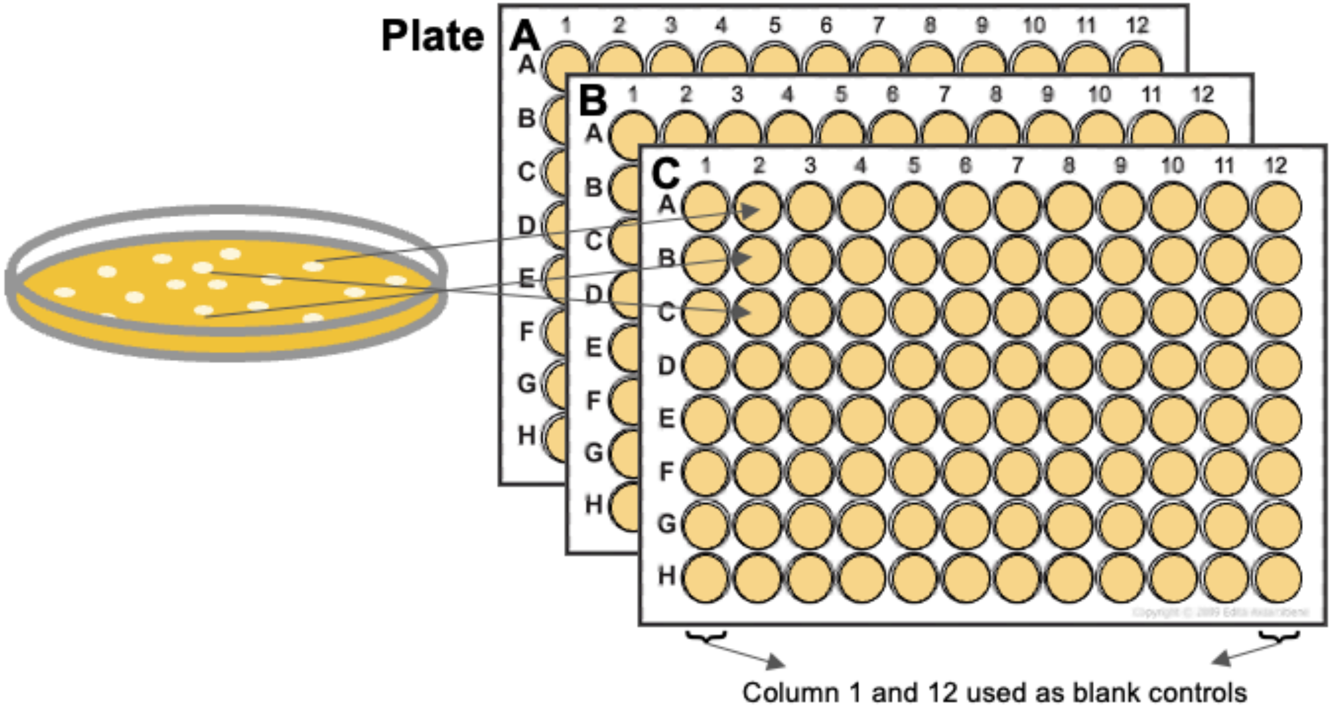


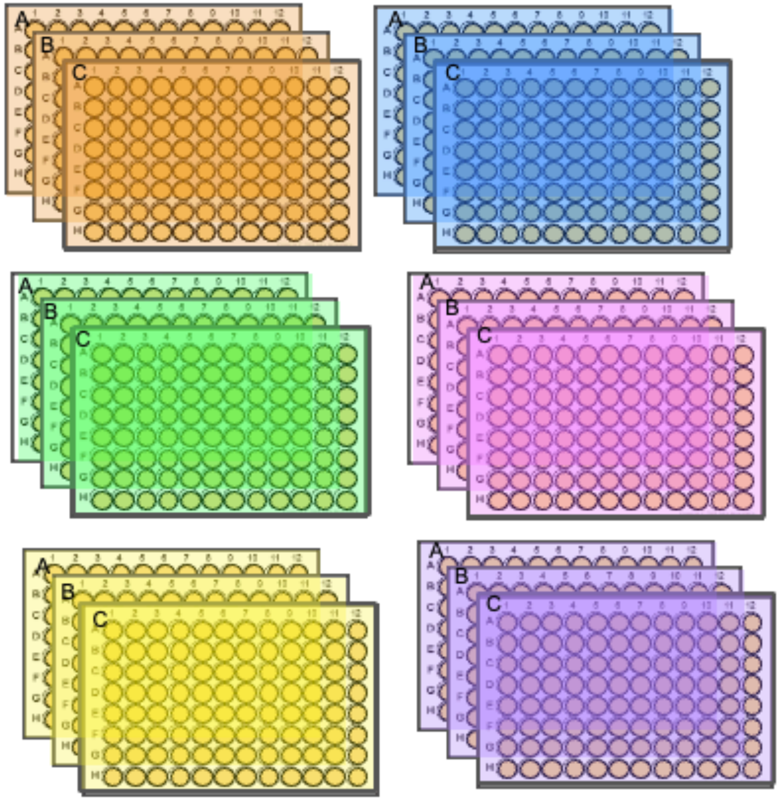

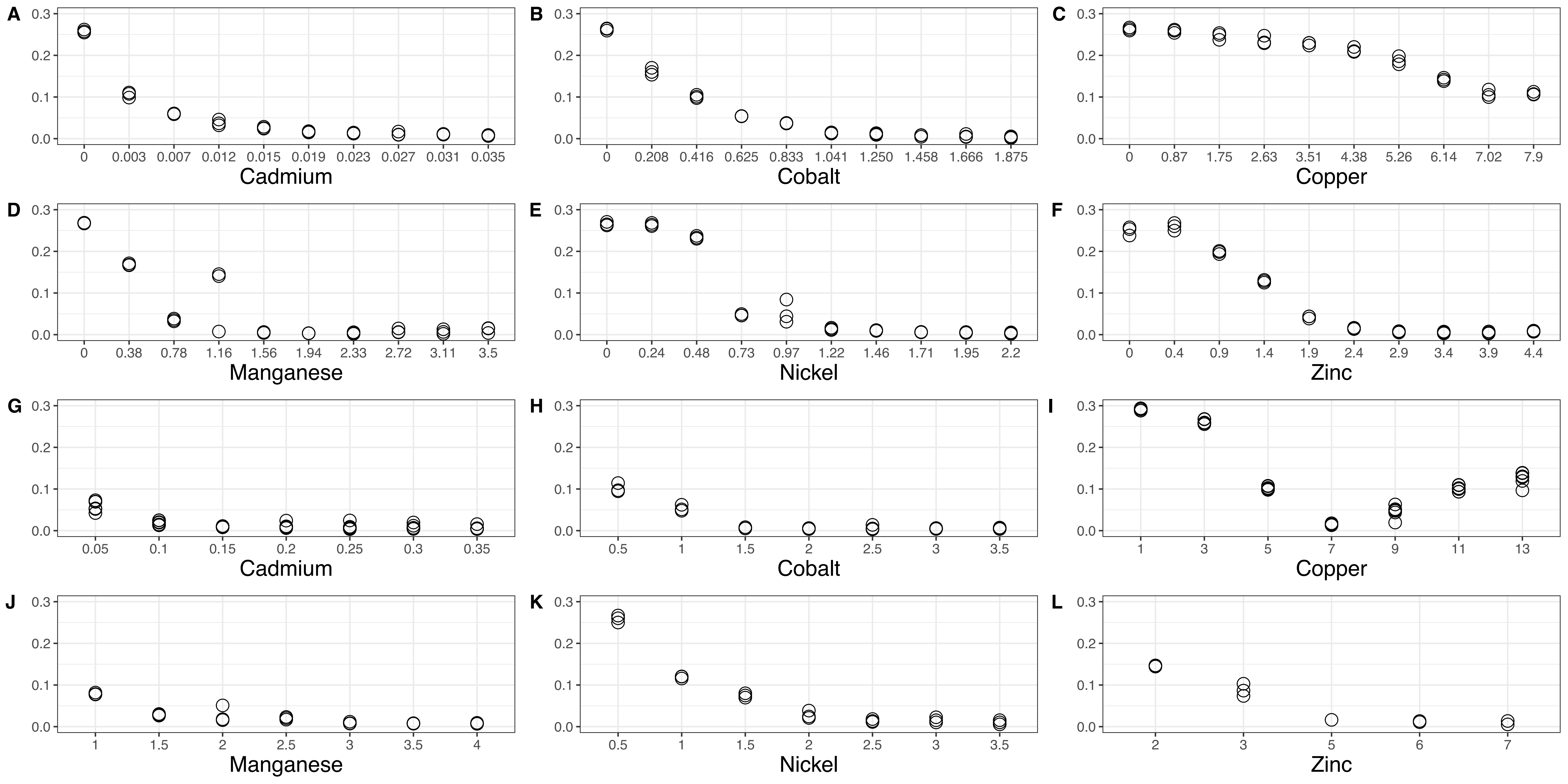
A
B
C
D
E
F
G
H
I
J
K
L
Cadmium
0 0.003 0.007 0.012 0.015 0.019 0.023 0.027 0.031 0.035
Cobalt
0 0.208 0.416 0.625 0.833 1.041 1.250 1.458 1.666 1.875
Copper
0 0.87 1.75 2.63 3.51 4.38 5.26 6.14 7.02 7.9
Manganese
0 0.38 0.78 1.16 1.56 1.94 2.33 2.72 3.11 3.5
Nickel
0 0.24 0.48 0.73 0.97 1.22 1.46 1.71 1.95 2.2
Zinc
0 0.4 0.9 1.4 1.9 2.4 2.9 3.4 3.9 4.4
Cadmium
0.05 0.1 0.15 0.2 0.25 0.3 0.35
Cobalt
0.5 1 1.5 2 2.5 3 3.5
Copper
1 3 5 7 9 11 13
Manganese
1 1.5 2 2.5 3 3.5 4
Nickel
0.5 1 1.5 2 2.5 3 3.5
Zinc
2 3 5 6 7
0.0 0.1 0.2 0.3

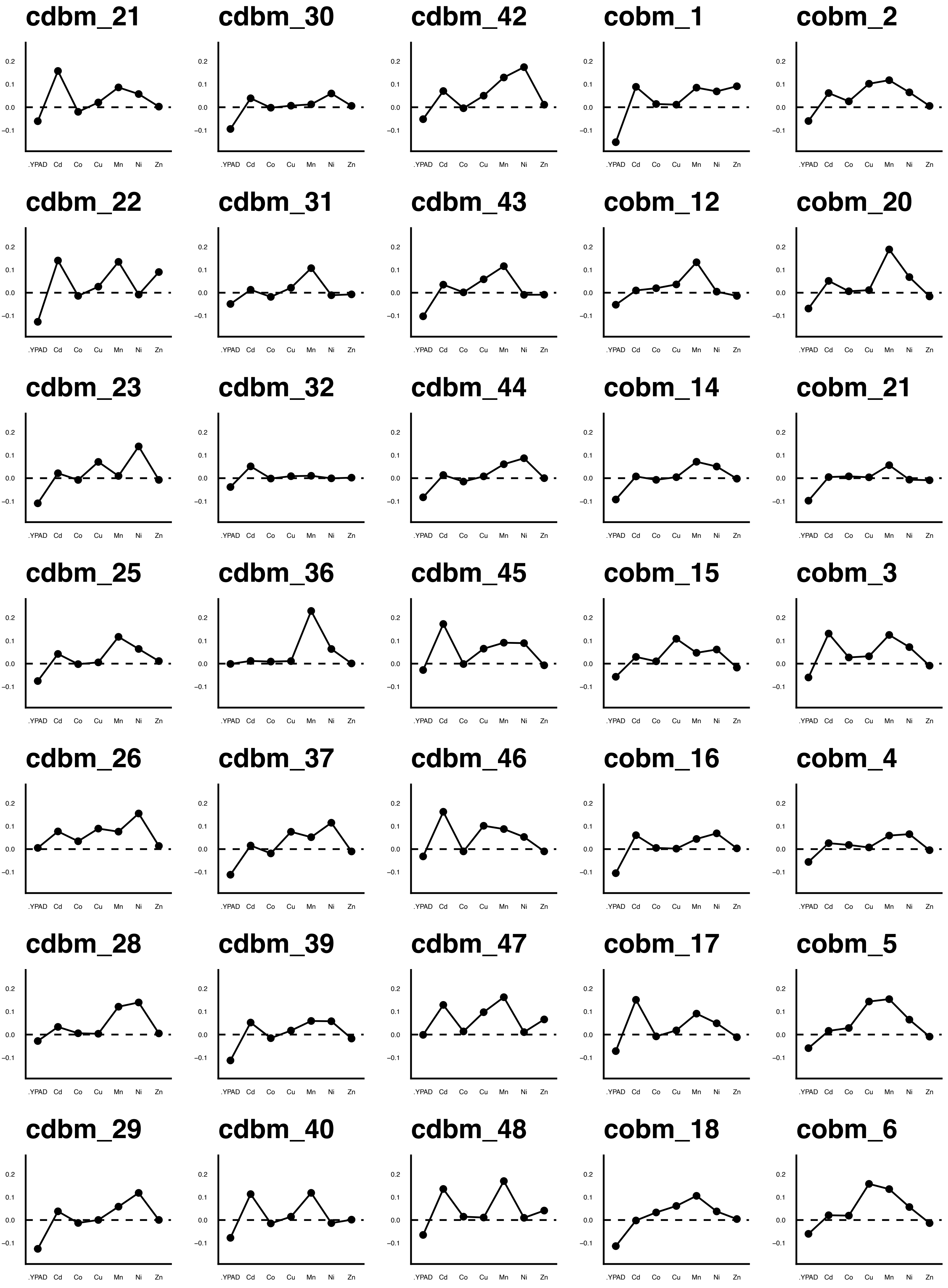
cdbm_21
cdbm_30
cdbm_42
cobm_1
cobm_2
cdbm_22
cdbm_31
cdbm_43
cobm_12
cobm_20
cdbm_23
cdbm_32
cdbm_44
cobm_14
cobm_21
cdbm_25
cdbm_36
cdbm_45
cobm_15
cobm_3
cdbm_26
cdbm_37
cdbm_46
cobm_16
cobm_4
cdbm_28
cdbm_39
cdbm_47
cobm_17
cobm_5
cdbm_29
cdbm_40
cdbm_48
cobm_18
cobm_6
0.2
0.1
0.0
−0.1
.YPAD
Cd
Co
Cu
Mn
Ni
Zn

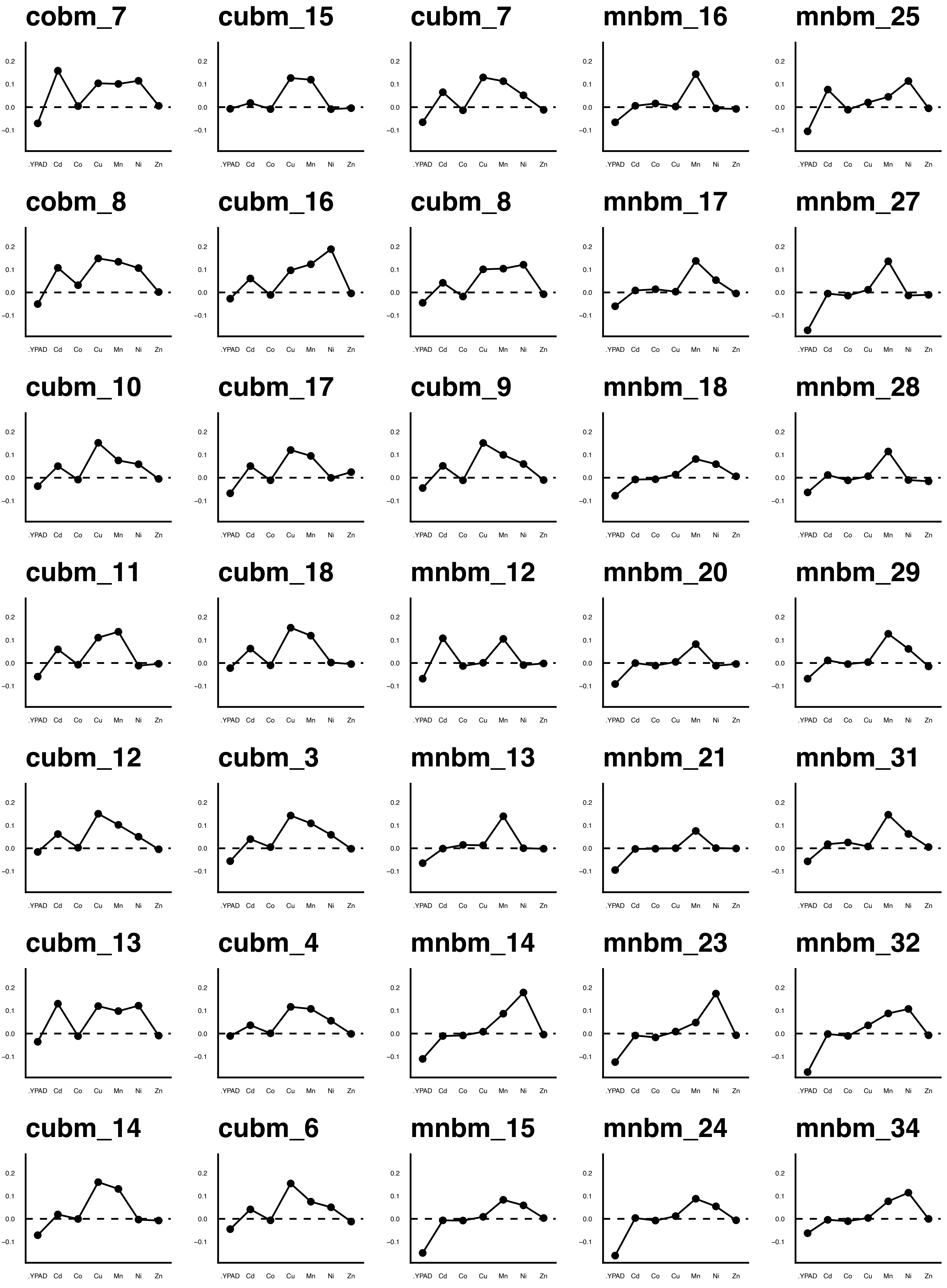
cobm_7
cubm_15
cubm_7
mnbm_16
mnbm_25
cobm_8
cubm_16
cubm_8
mnbm_17
mnbm_27
cubm_10
cubm_17
cubm_9
mnbm_18
mnbm_28
cubm_11
cubm_18
mnbm_12
mnbm_20
mnbm_29
cubm_12
cubm_3
mnbm_13
mnbm_21
mnbm_31
cubm_13
cubm_4
mnbm_14
mnbm_23
mnbm_32
cubm_14
cubm_6
mnbm_15
mnbm_24
mnbm_34
0.2
0.1
0.0
-0.1
.YPAD
Cd
Co
Cu
Mn
Ni
Zn

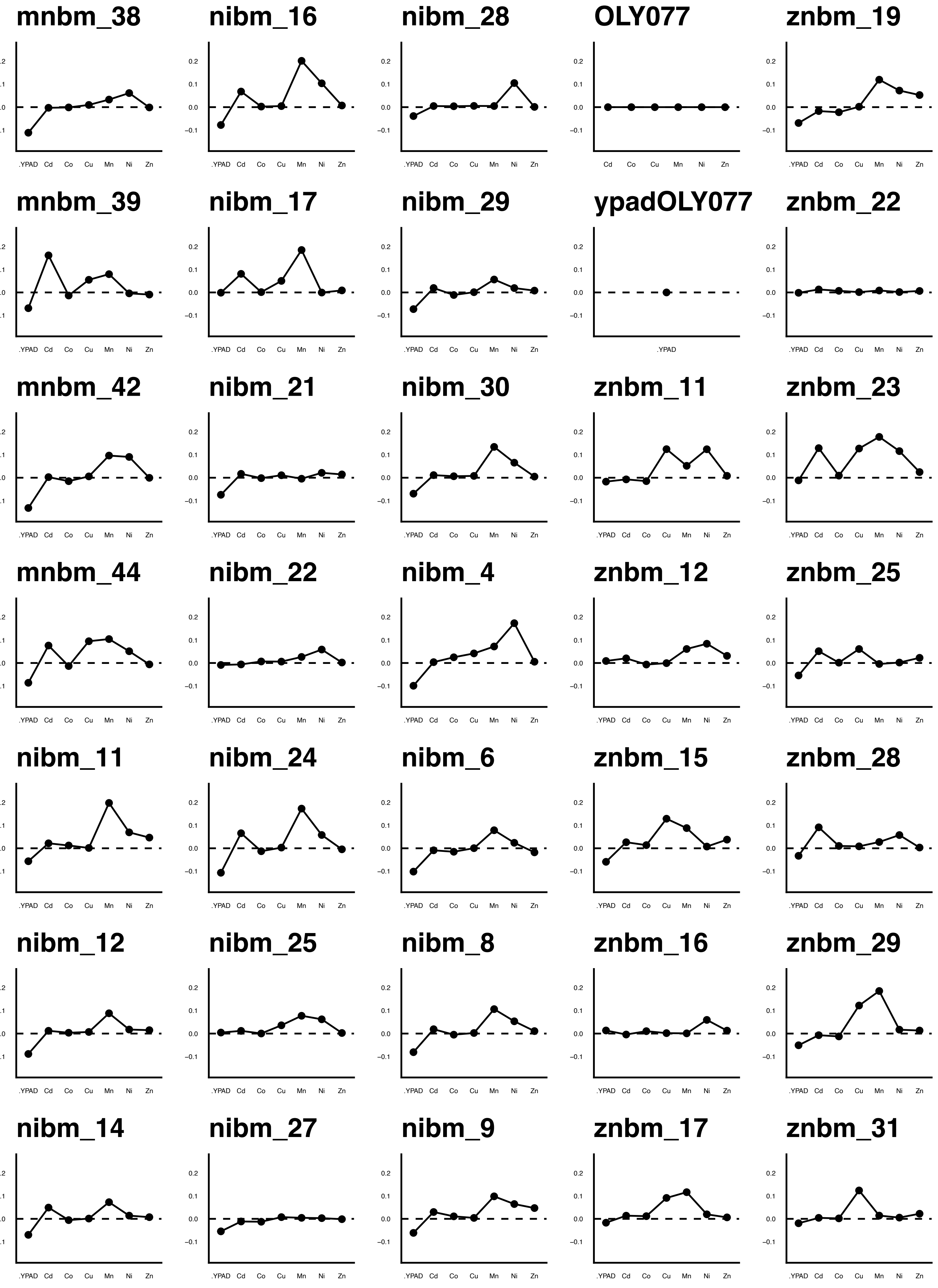
mnbm_38
nibm_16
nibm_28
OLY077
znbm_19
mnbm_39
nibm_17
nibm_29
ypadOLY077
znbm_22
mnbm_42
nibm_21
nibm_30
znbm_11
znbm_23
mnbm_44
nibm_22
nibm_4
znbm_12
znbm_25
nibm_11
nibm_24
nibm_6
znbm_15
znbm_28
nibm_12
nibm_25
nibm_8
znbm_16
znbm_29
nibm_14
nibm_27
nibm_9
znbm_17
znbm_31
0.2
0.1
0.0
-0.1
.YPAD
Cd
Co
Cu
Mn
Ni
Zn

#### znbm_32

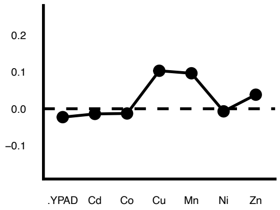


#### znbm_34

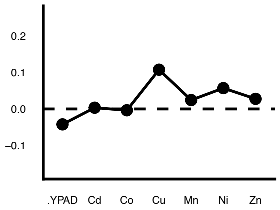


#### znbm_37

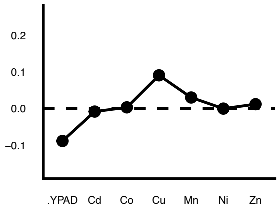


#### znbm_38

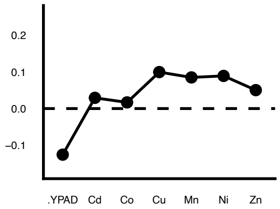


#### znbm_39

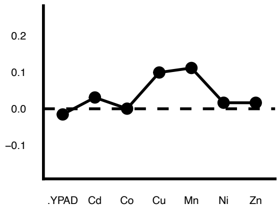


#### znbm_41

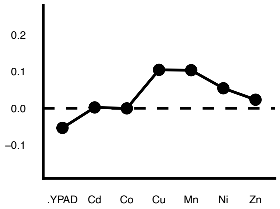


#### znbm_42

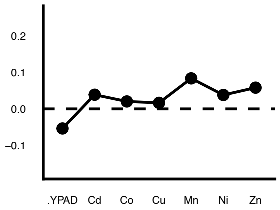


#### znbm_43

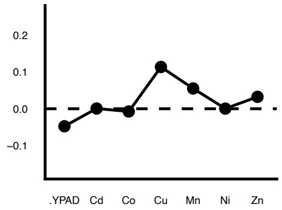


#### znbm_44

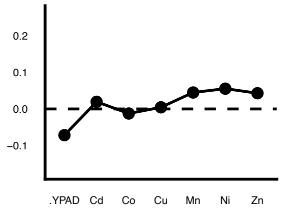


#### znbm_45

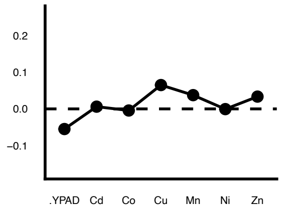


#### znbm_46

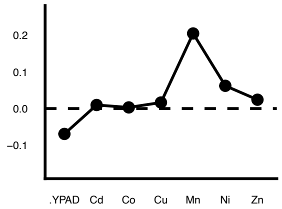


#### znbm_47

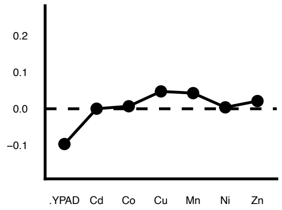


#### znbm_5

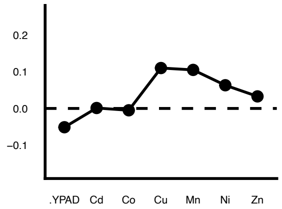


#### znbm_6

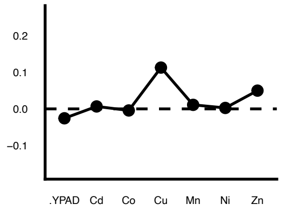


#### znbm_7

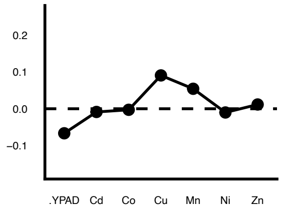


#### znbm_8

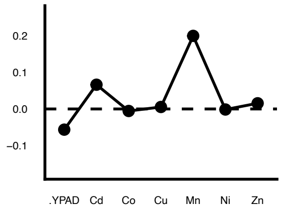


#### znbm_9

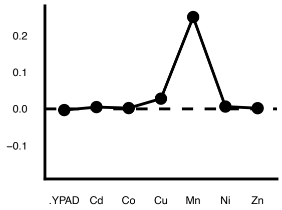

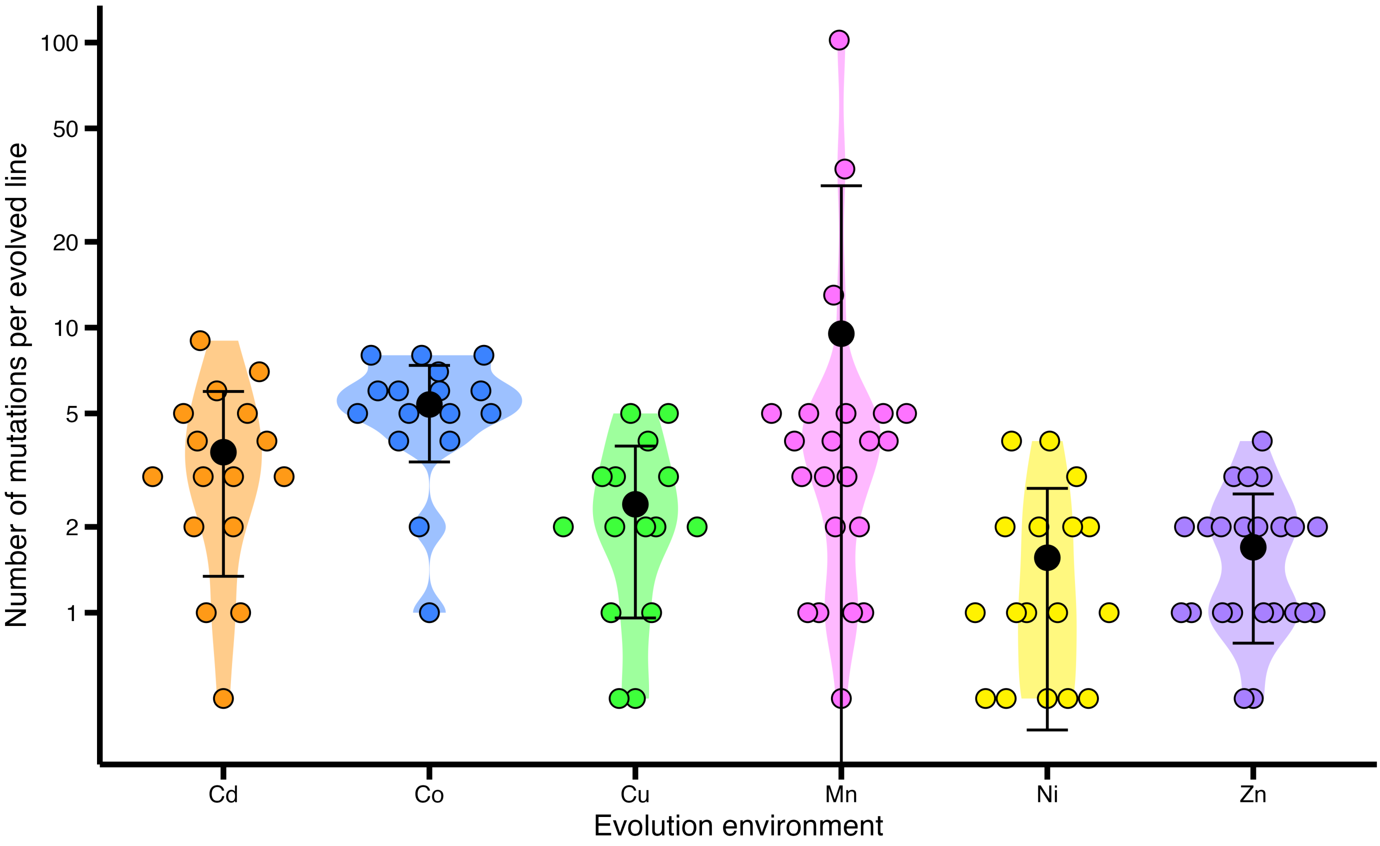
Number of mutations per evolved line
100
50
20
10
5
2
1
Cd
Co
Cu
Mn
Ni
Zn
Evolution environment

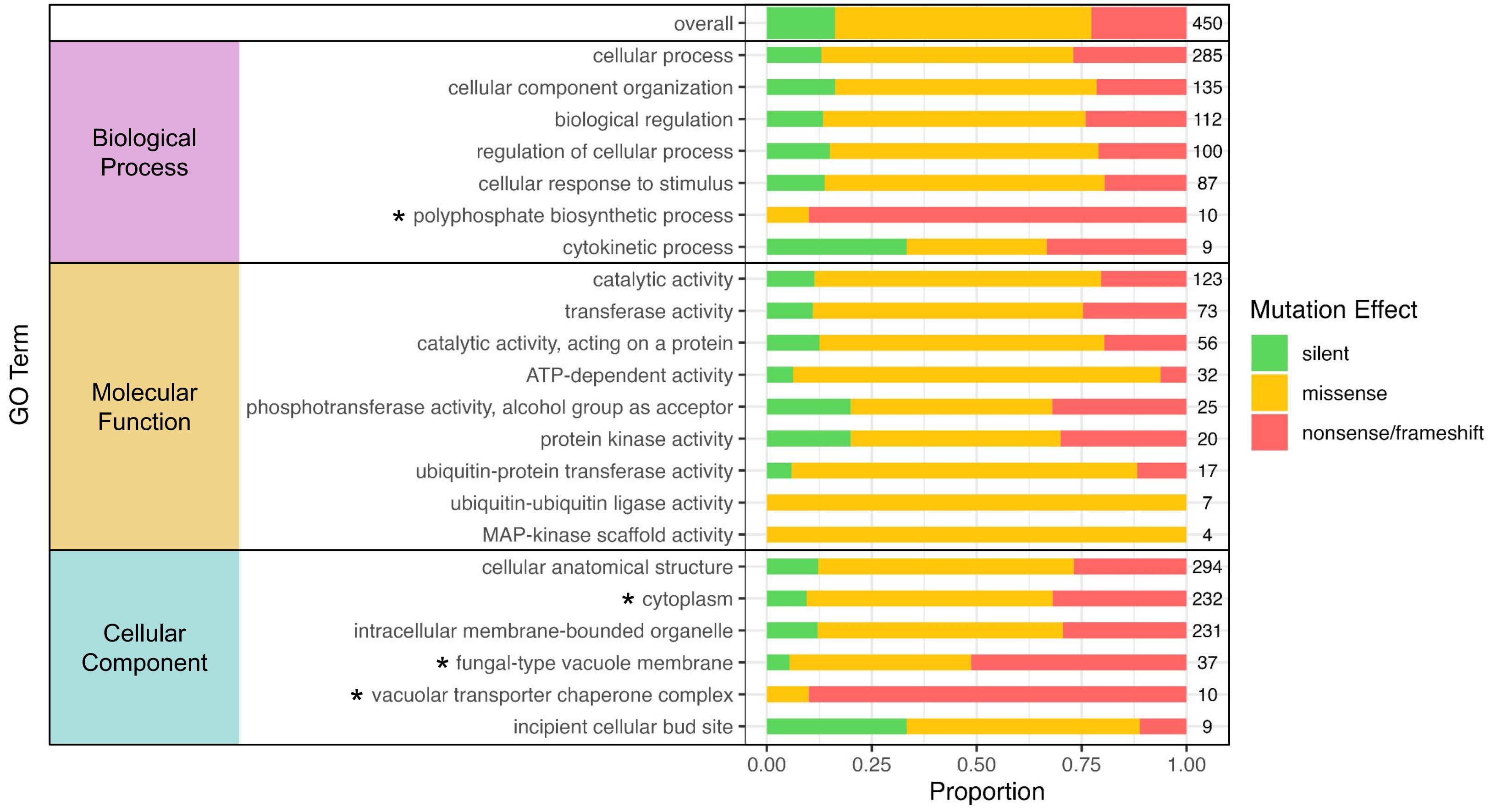

GO Term
overall
Biological Process
cellular process
cellular component organization
biological regulation
regulation of cellular process
cellular response to stimulus
* polyphosphate biosynthetic process
cytokinetic process
Molecular Function
catalytic activity
transferase activity
catalytic activity, acting on a protein
ATP-dependent activity
phosphotransferase activity, alcohol group as acceptor
protein kinase activity
ubiquitin-protein transferase activity
ubiquitin-ubiquitin ligase activity
MAP-kinase scaffold activity
Cellular Component
cellular anatomical structure
* cytoplasm
intracellular membrane-bounded organelle
* fungal-type vacuole membrane
* vacuolar transporter chaperone complex
incipient cellular bud site
450
285
135
112
100
87
10
9
123
73
56
32
25
20
17
7
4
294
232
231
37
10
9
0.00
0.25
0.50
0.75
1.00
Proportion
Mutation Effect
silent
missense
nonsense/frameshift

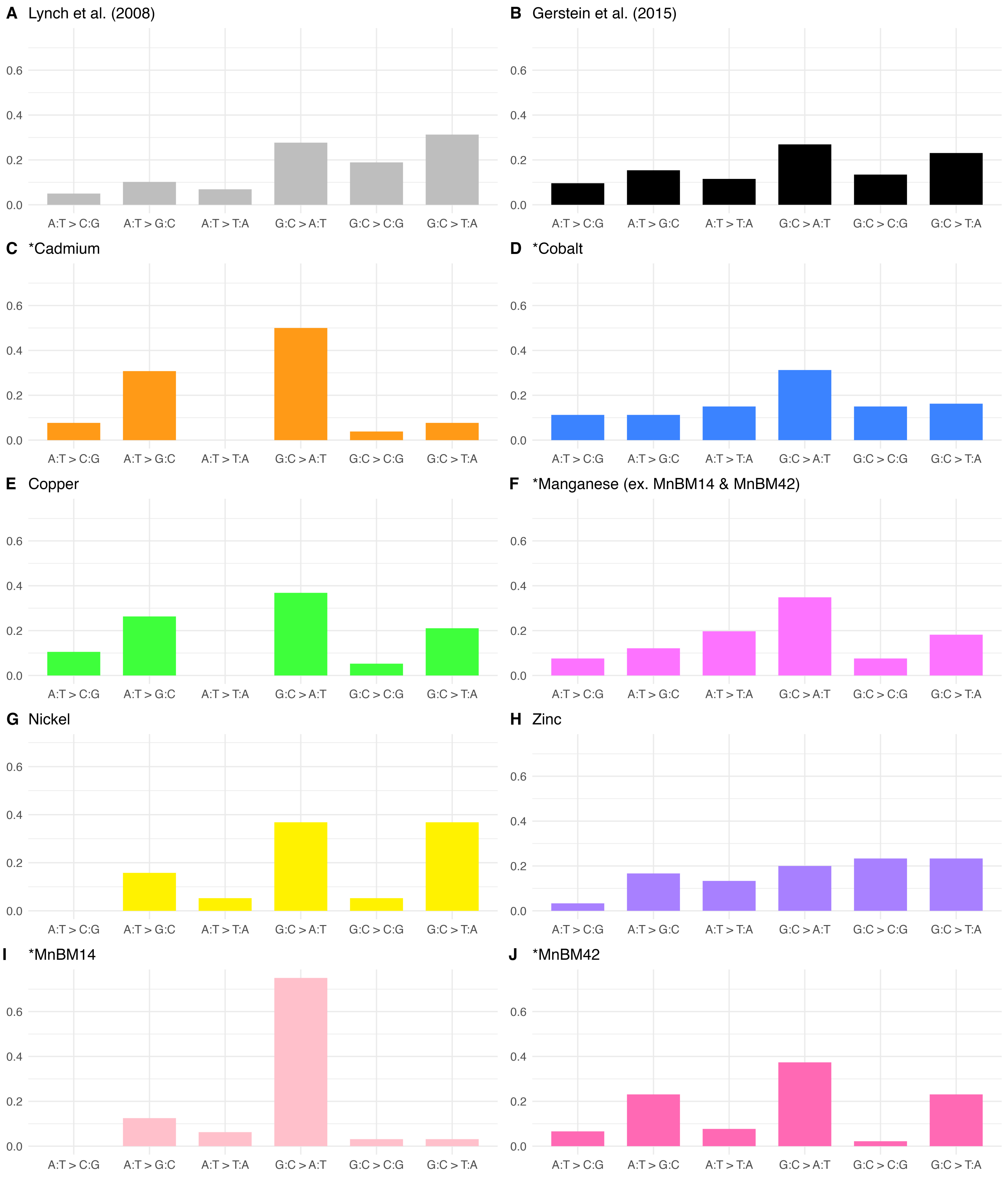

A Lynch et al. (2008)
B Gerstein et al. (2015)
C *Cadmium
D *Cobalt
E Copper
F *Manganese (ex. MnBM14 & MnBM42)
G Nickel
H Zinc
I *MnBM14
J *MnBM42
0.0
0.2
0.4
0.6
A:T > C:G
A:T > G:C
A:T > T:A
G:C > A:T
G:C > C:G
G:C > T:A

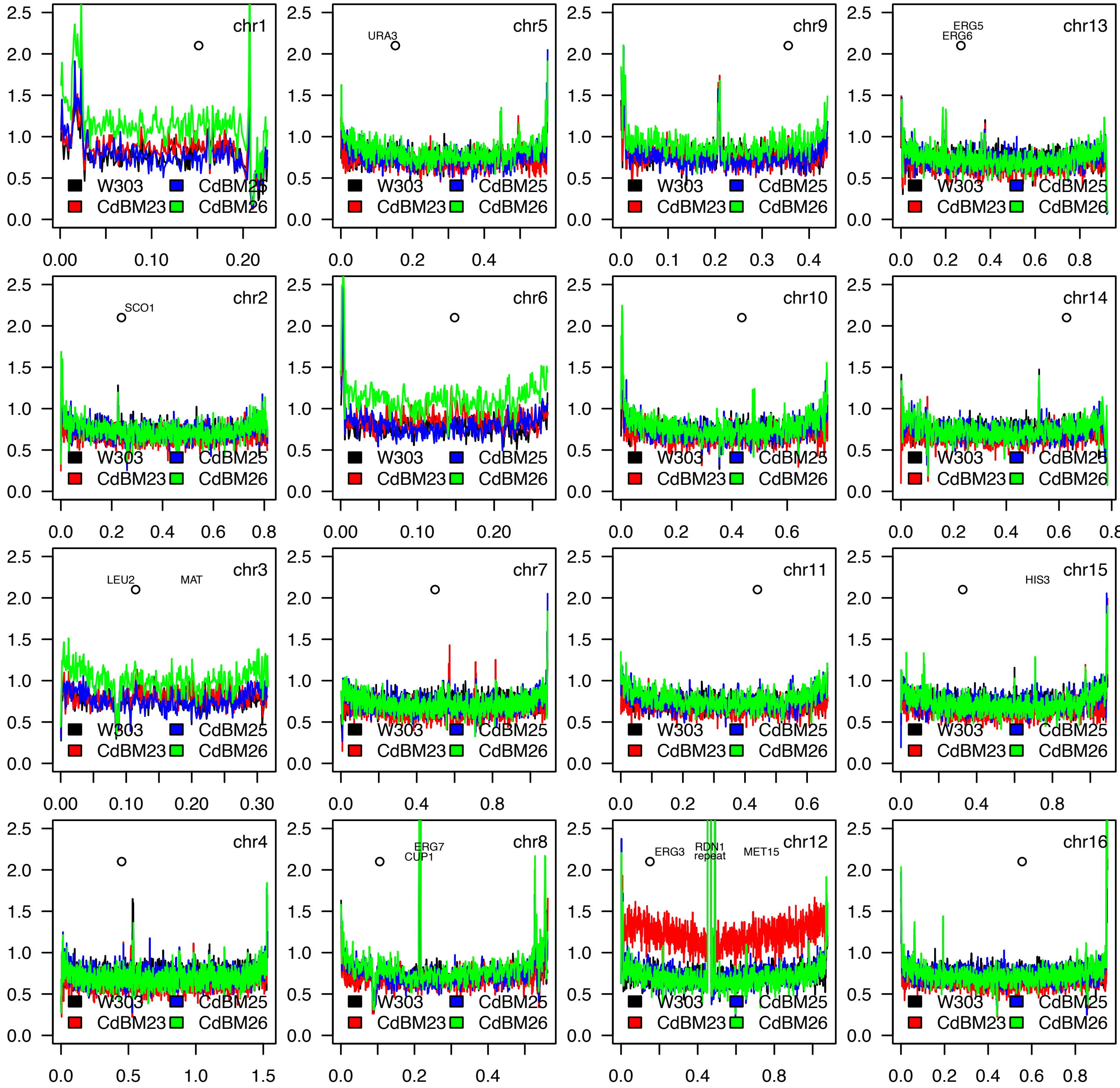
chr1
chr5
URA3
chr9
chr13
ERG5
ERG6
chr2
SCO1
chr6
chr10
chr14
chr3
LEU2
MAT
chr7
chr11
chr15
HIS3
chr4
chr8
ERG7
CUP1
chr12
ERG3
RDN1 repeat
MET15
chr16
W303
CdBM25
CdBM23
CdBM26

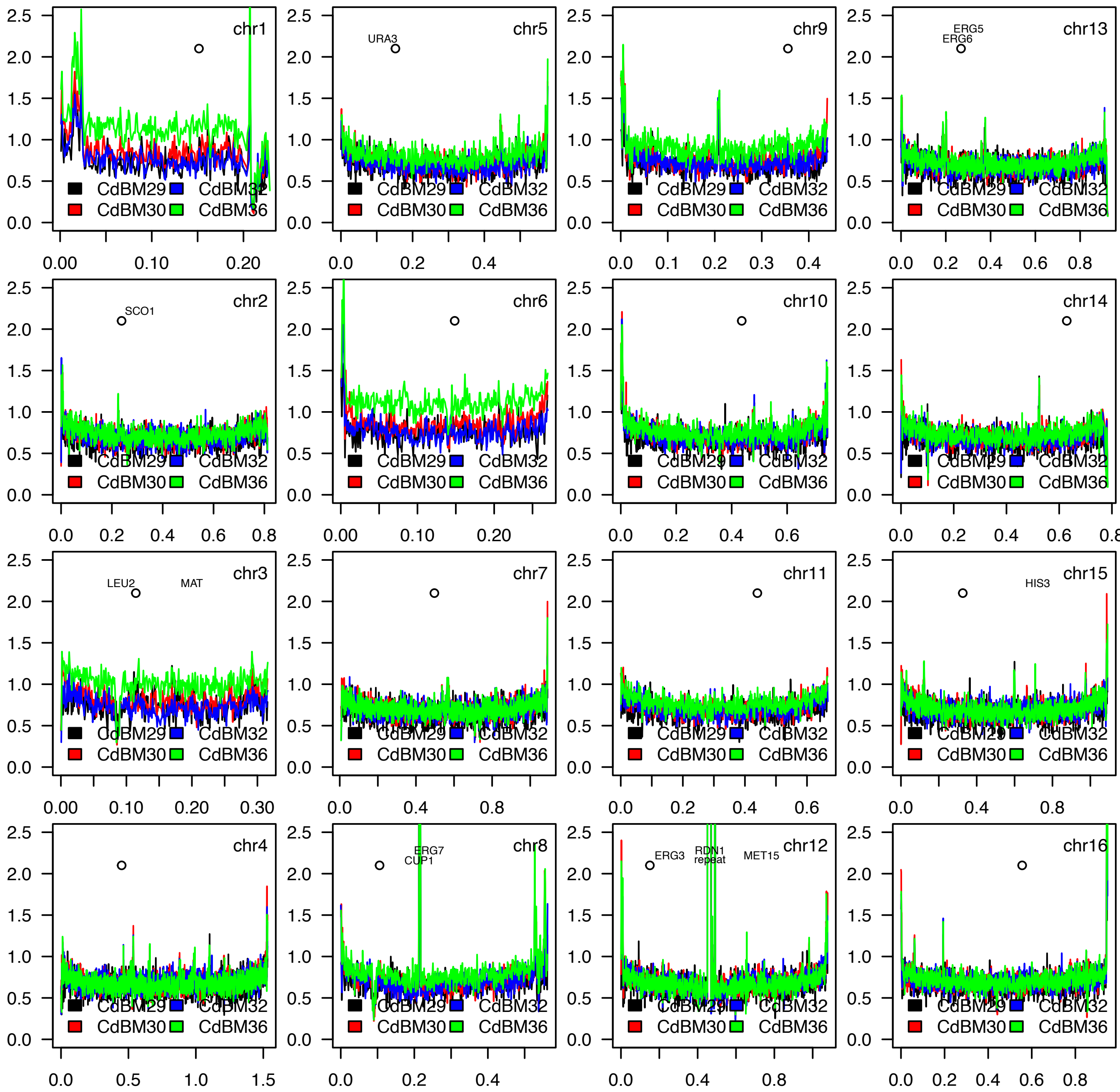
chr1
chr5
URA3
chr9
chr13
ERG5
ERG6
chr2
SCO1
chr6
chr10
chr14
chr3
LEU2
MAT
chr7
chr11
chr15
HIS3
chr4
chr8
ERG7
CUP1
chr12
ERG3
RDN1
repeat
MET15
chr16
CdBM29
CdBM30
CdBM32
CdBM36

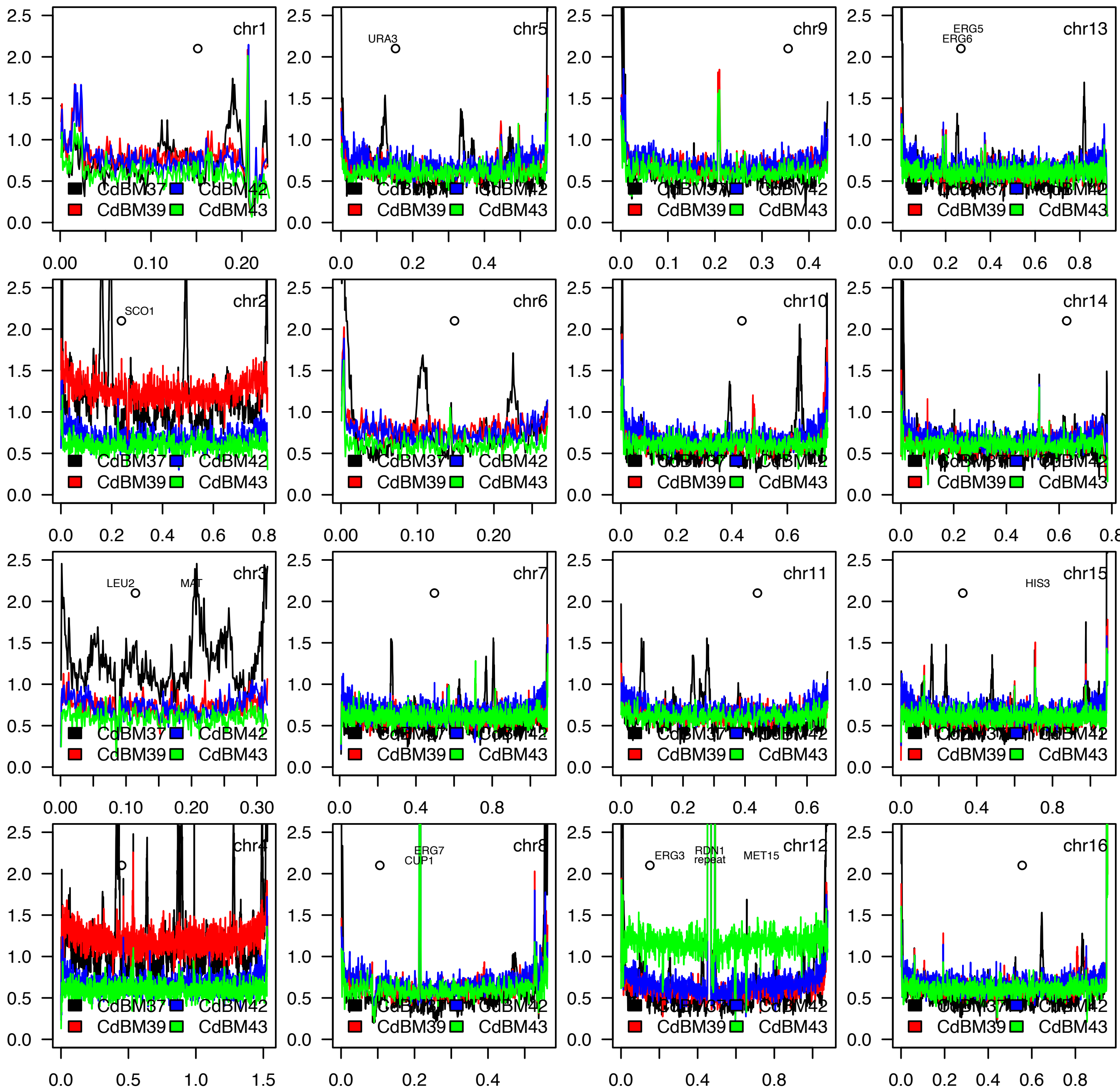
chr1
chr5
URA3
chr9
chr13
ERG5
ERG6
chr2
SCO1
chr6
chr10
chr14
chr3
LEU2
MAT
chr7
chr11
chr15
HIS3
chr4
chr8
ERG7
CUP1
chr12
ERG3
RDN1
repeat
MET15
chr16
CdBM37
CdBM42
CdBM39
CdBM43

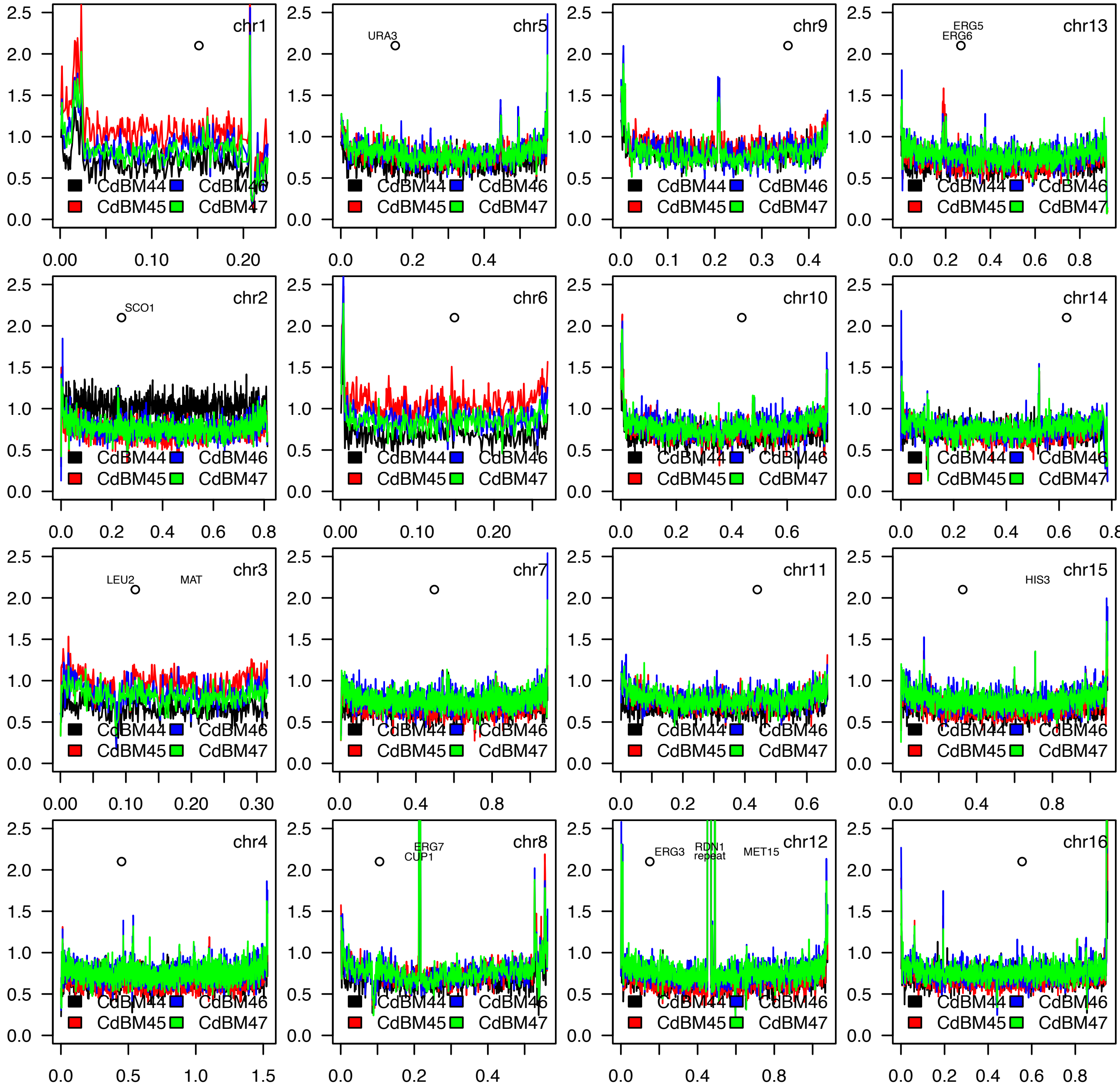
chr1
chr5
URA3
chr9
chr13
ERG5
ERG6
chr2
SCO1
chr6
chr10
chr14
chr3
LEU2
MAT
chr7
chr11
chr15
HIS3
chr4
chr8
ERG7
CUP1
chr12
ERG3
RDN1
repeat
MET15
chr16
CdBM44
CdBM45
CdBM46
CdBM47

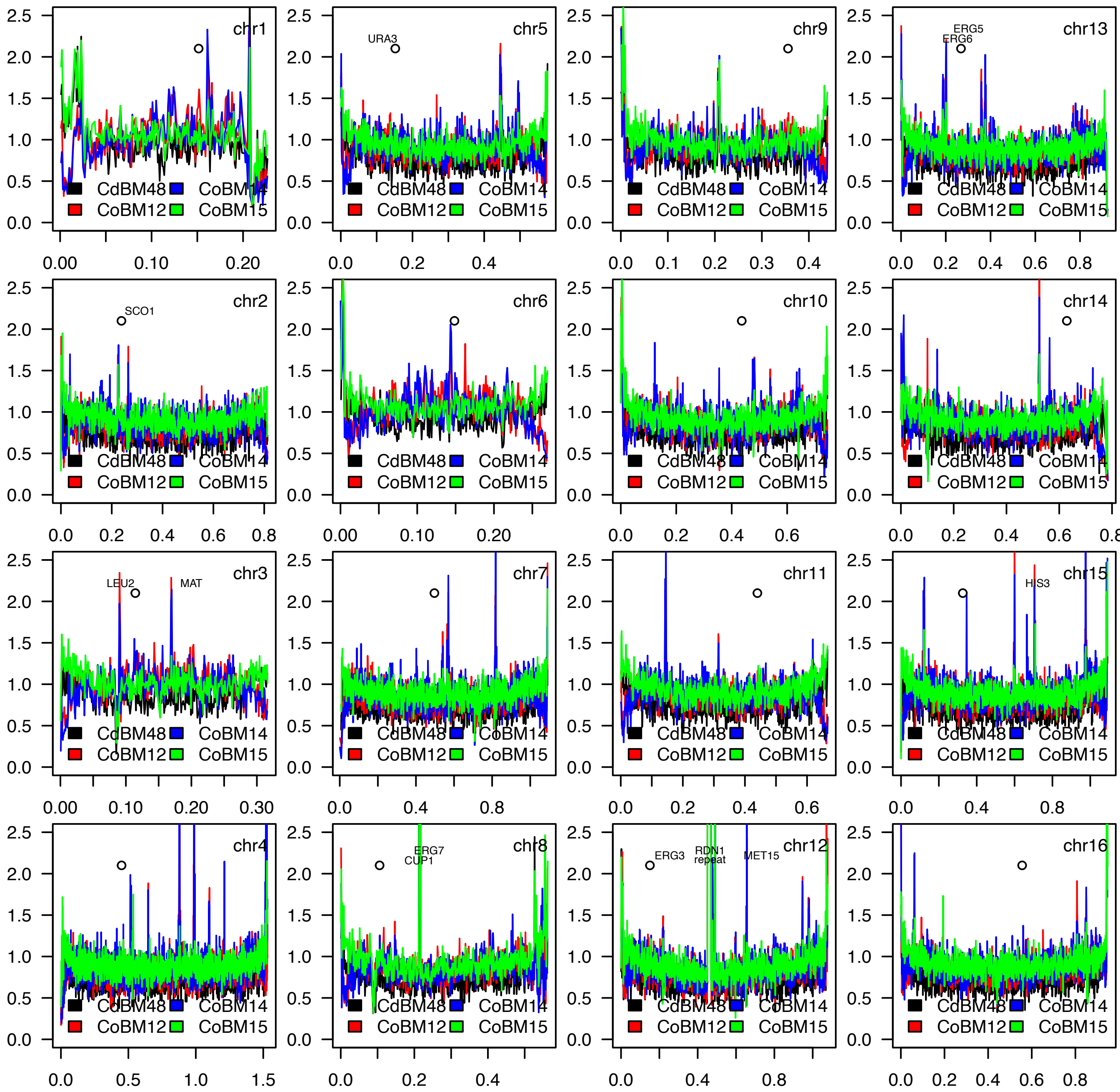
chr1
CdBM48 CoBM14 CoBM12 CoBM15
chr5
URA3
CdBM48 CoBM14 CoBM12 CoBM15
chr9
CdBM48 CoBM14 CoBM12 CoBM15
chr13
ERG5
ERG6
CdBM48 CoBM14 CoBM12 CoBM15
chr2
SCO1
CdBM48 CoBM14 CoBM12 CoBM15
chr6
CdBM48 CoBM14 CoBM12 CoBM15
chr10
CdBM48 CoBM14 CoBM12 CoBM15
chr14
CdBM48 CoBM14 CoBM12 CoBM15
chr3
LEU2
MAT
CdBM48 CoBM14 CoBM12 CoBM15
chr7
CdBM48 CoBM14 CoBM12 CoBM15
chr11
CdBM48 CoBM14 CoBM12 CoBM15
chr15
HIS3
CdBM48 CoBM14 CoBM12 CoBM15
chr4
CdBM48 CoBM14 CoBM12 CoBM15
chr8
ERG7
CUP1
CdBM48 CoBM14 CoBM12 CoBM15
chr12
ERG3
RDN1 repeat
MET15
CdBM48 CoBM14 CoBM12 CoBM15
chr16
CdBM48 CoBM14 CoBM12 CoBM15

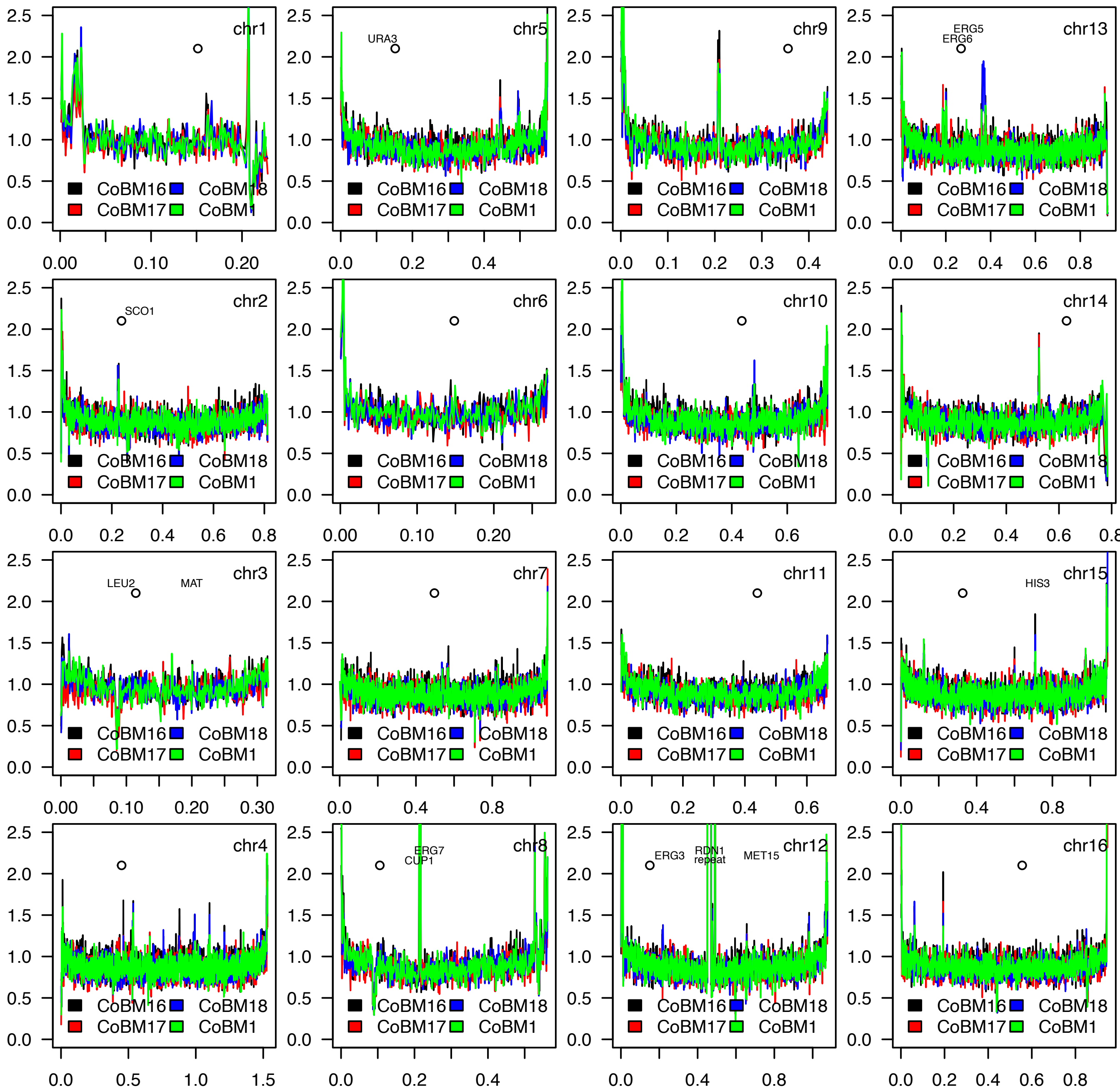
chr1
chr5
URA3
chr9
chr13
ERG5
ERG6
chr2
SCO1
chr6
chr10
chr14
chr3
LEU2
MAT
chr7
chr11
chr15
HIS3
chr4
chr8
ERG7
CUP1
chr12
ERG3
RDN1
repeat
MET15
chr16
CoBM16
CoBM17
CoBM18
CoBM1

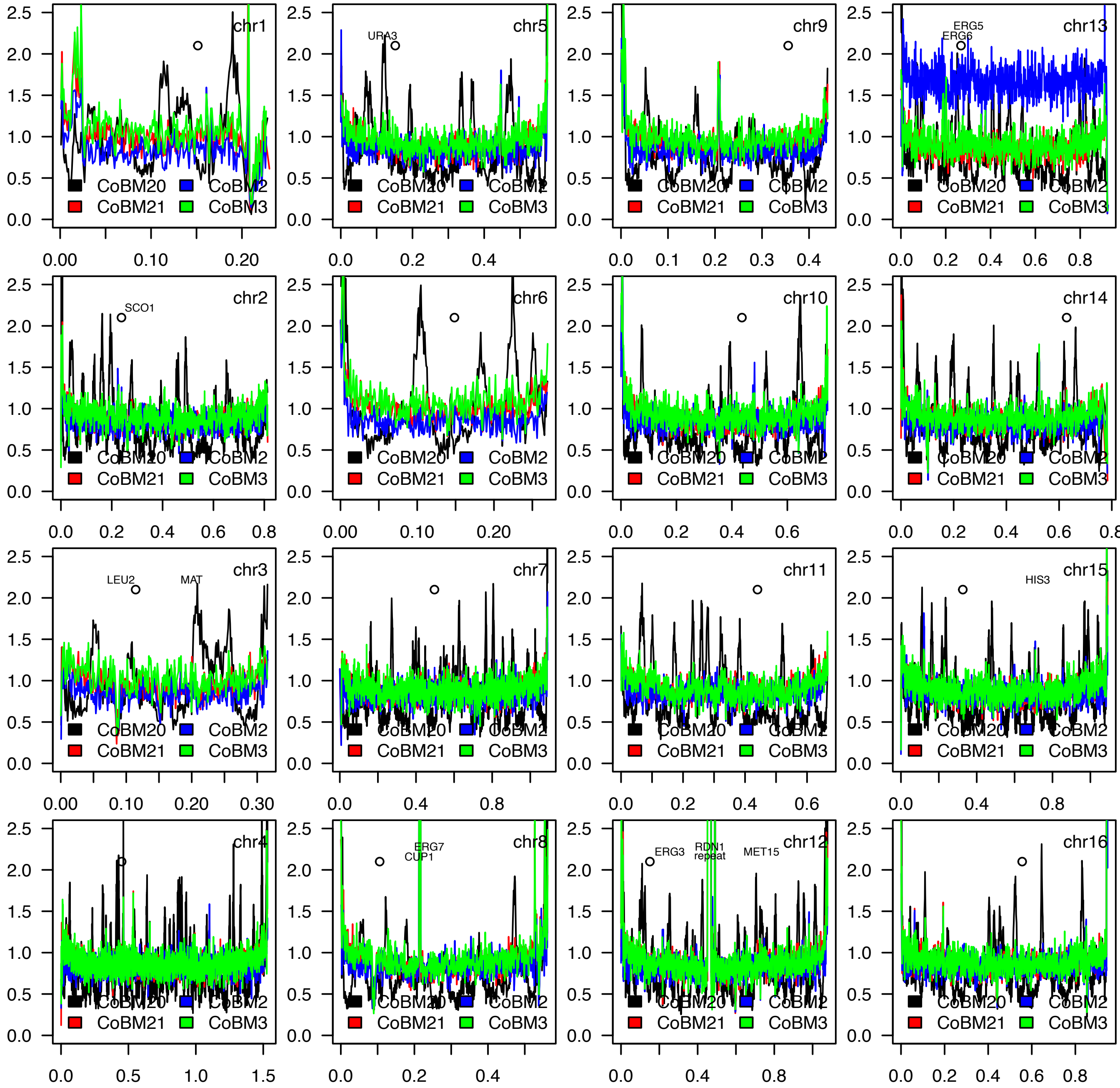

chr1
chr5
URA3
chr9
chr13
ERG5
ERG6
chr2
SCO1
chr6
chr10
chr14
chr3
LEU2
MAT
chr7
chr11
chr15
HIS3
chr4
chr8
ERG7
CUP1
chr12
ERG3
RDN1
repeat
MET15
chr16
CoBM20
CoBM2
CoBM21
CoBM3

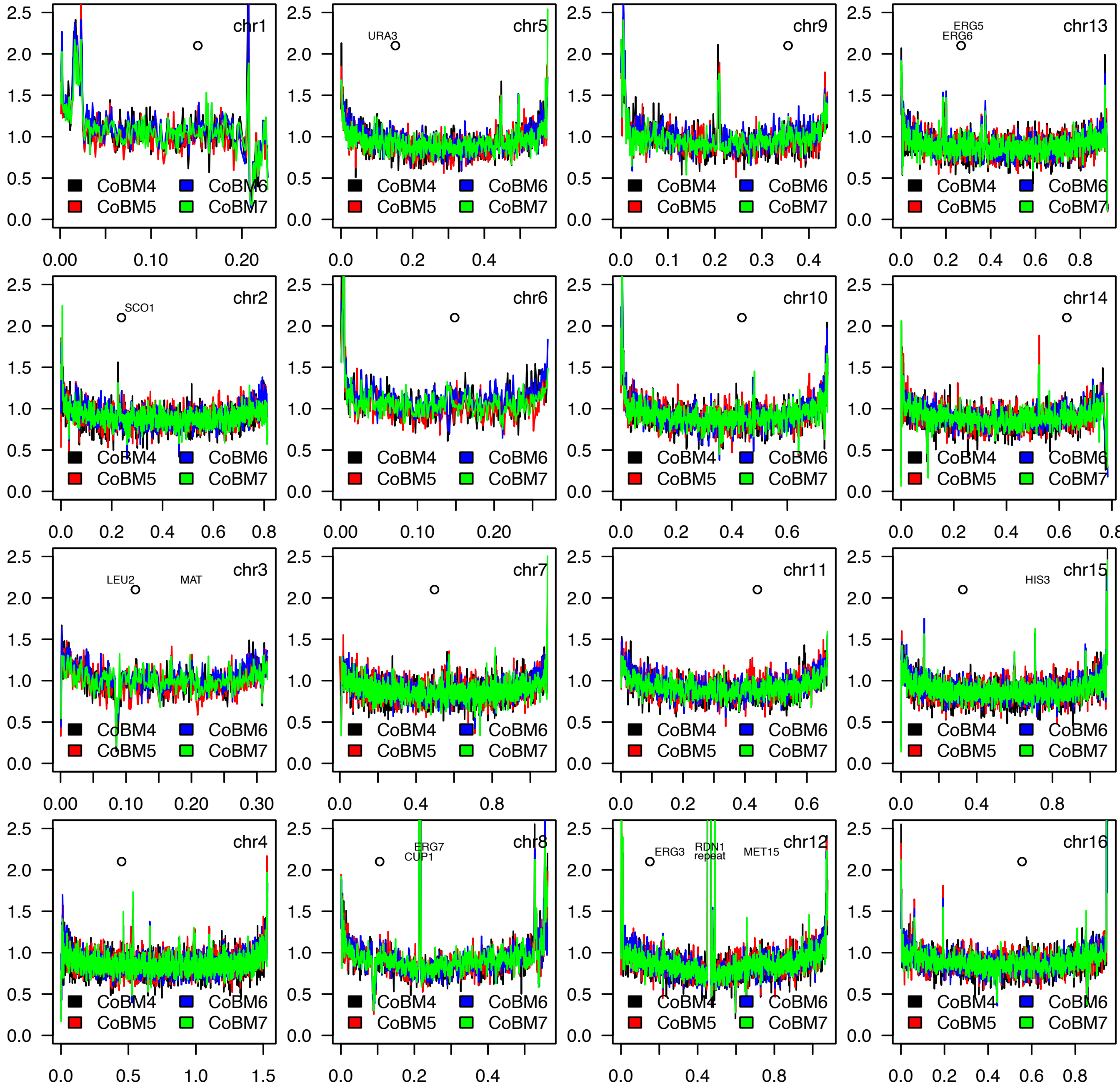

chr1
chr5
URA3
chr9
chr13
ERG5
ERG6
chr2
SCO1
chr6
chr10
chr14
chr3
LEU2
MAT
chr7
chr11
chr15
HIS3
chr4
chr8
ERG7
CUP1
chr12
ERG3
RDN1
repeat
MET15
chr16
CoBM4
CoBM5
CoBM6
CoBM7
0.0
0.5
1.0
1.5
2.0
2.5

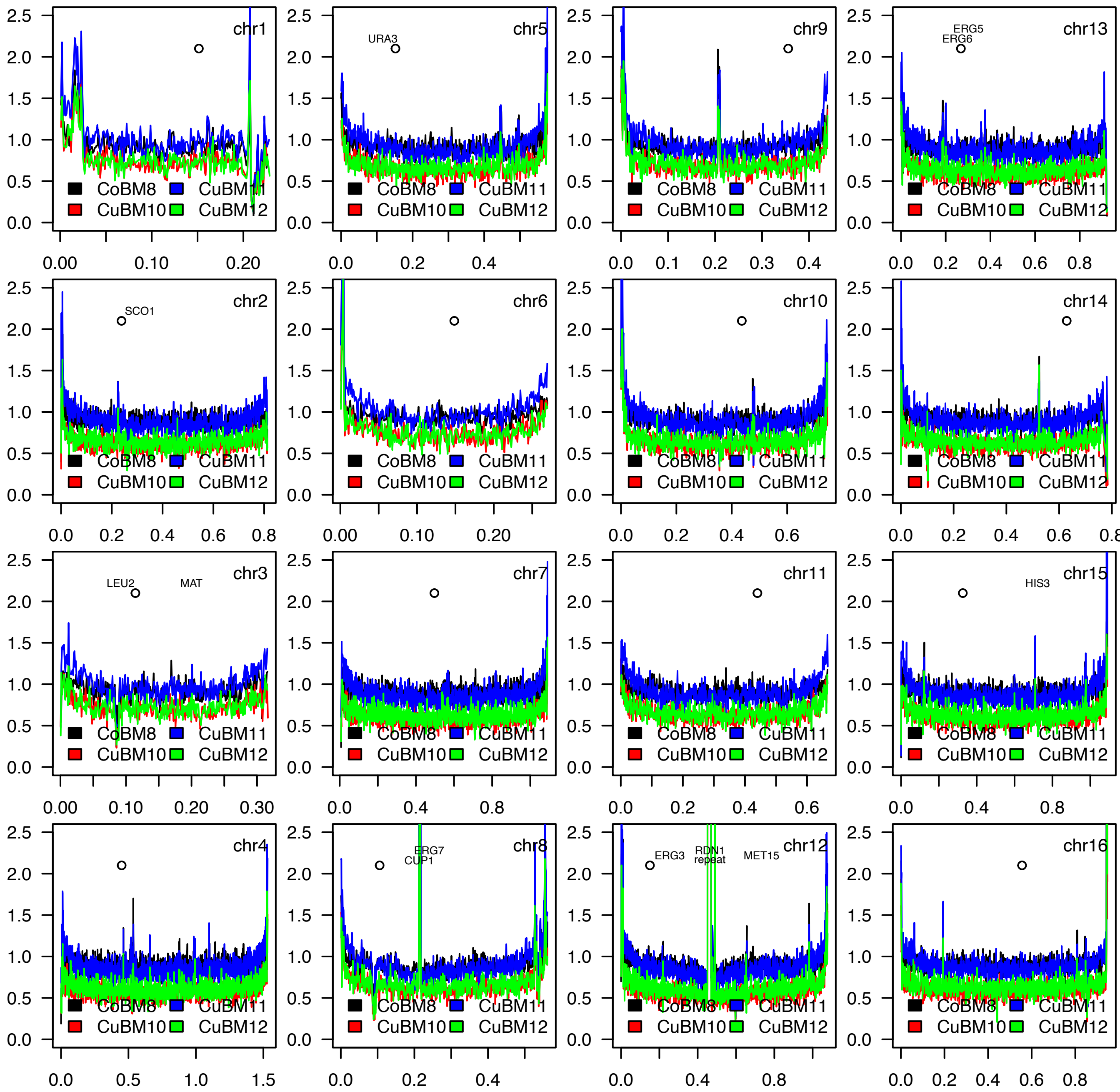
chr1
CoBM8
CuBM11
CuBM10
CuBM12
chr5
URA3
chr9
chr13
ERG5
ERG6
chr2
SCO1
chr6
chr10
chr14
chr3
LEU2
MAT
chr7
chr11
chr15
HIS3
chr4
chr8
ERG7
CUP1
chr12
ERG3
RDN1
repeat
MET15
chr16

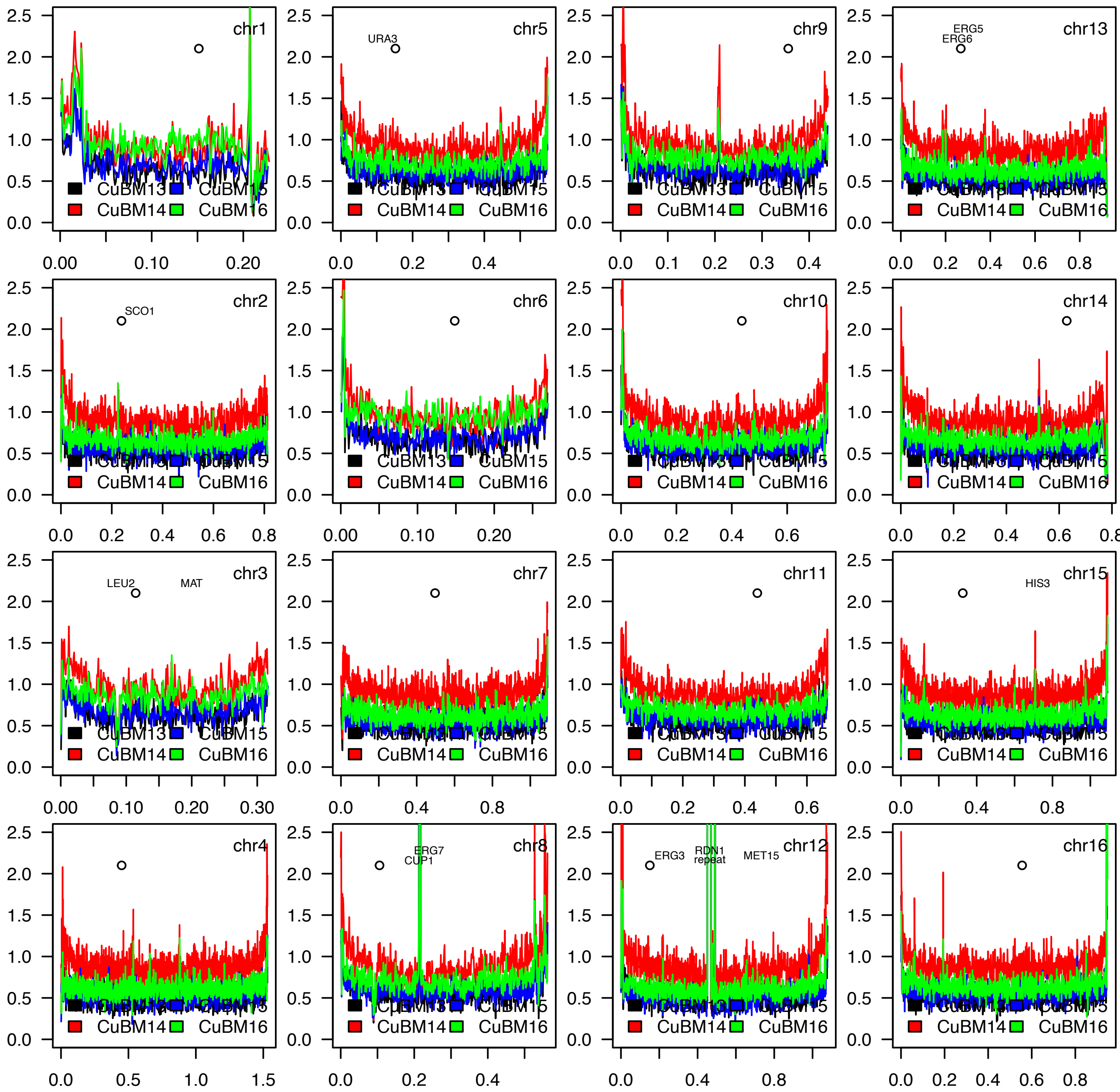
chr1
chr5
URA3
chr9
chr13
ERG5
ERG6
chr2
SCO1
chr6
chr10
chr14
chr3
LEU2
MAT
chr7
chr11
chr15
HIS3
chr4
chr8
ERG7
CUP1
chr12
ERG3
RDN1
repeat
MET15
chr16
CuBM13
CuBM15
CuBM14
CuBM16

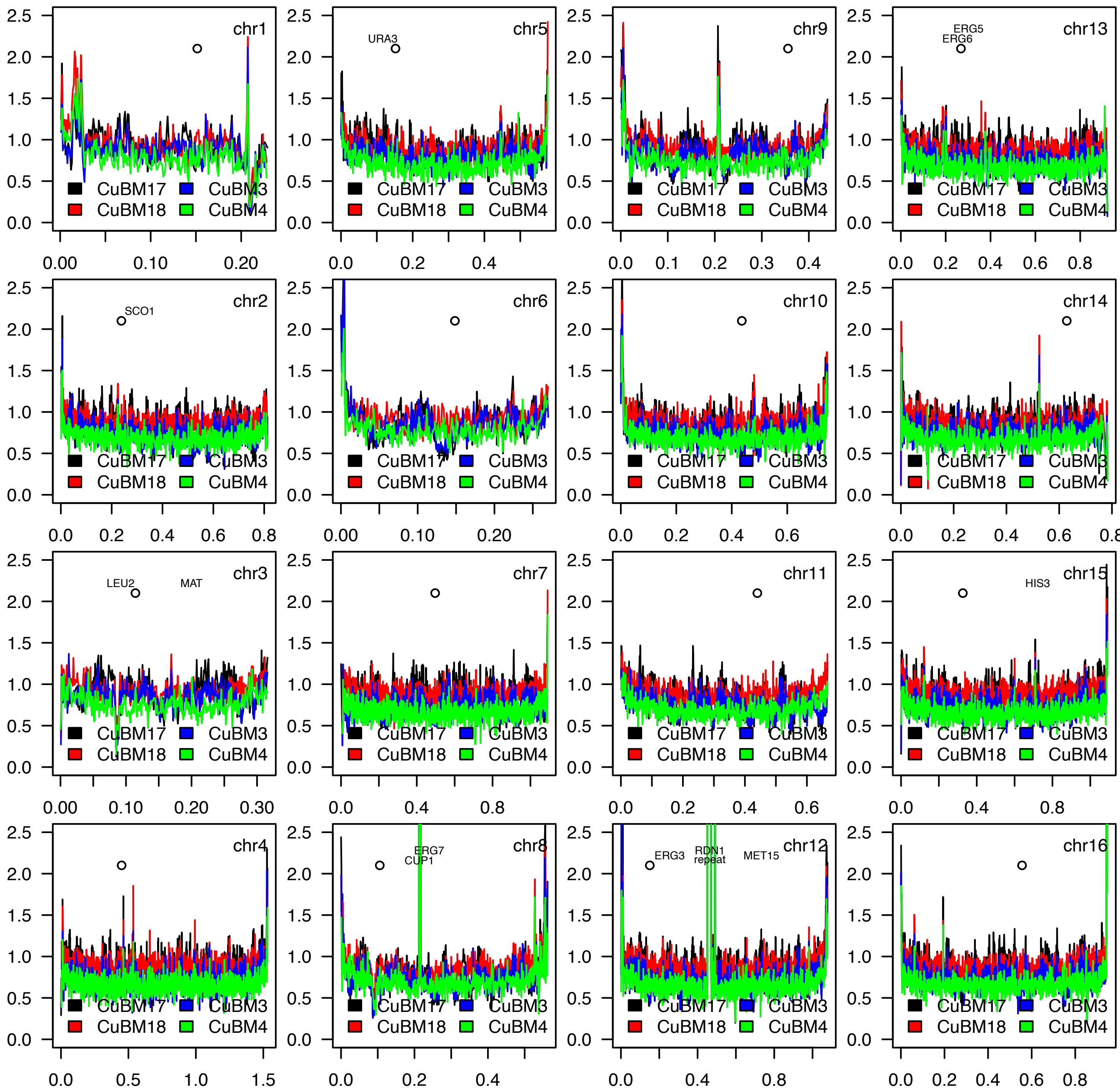
chr1
CuBM17
CuBM3
CuBM18
CuBM4
chr5
URA3
chr9
chr13
ERG5
ERG6
chr2
SCO1
chr6
chr10
chr14
chr3
LEU2
MAT
chr7
chr11
chr15
HIS3
chr4
chr8
ERG7
CUP1
chr12
ERG3
RDN1
repeat
MET15
chr16

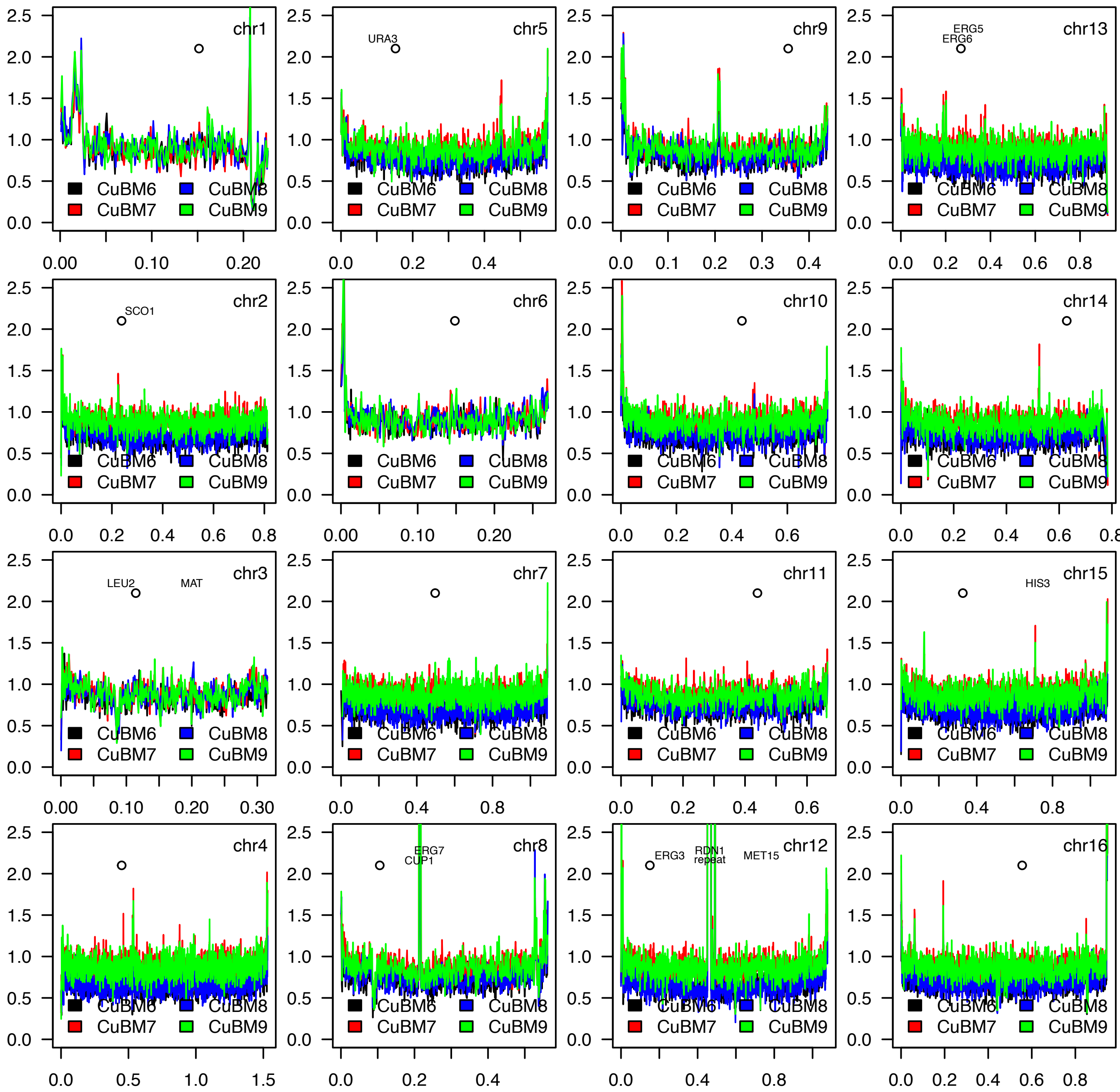
chr1
chr5
URA3
chr9
chr13
ERG5
ERG6
chr2
SCO1
chr6
chr10
chr14
chr3
LEU2
MAT
chr7
chr11
chr15
HIS3
chr4
chr8
ERG7
CUP1
chr12
ERG3
RDN1
repeat
MET15
chr16
CuBM6
CuBM7
CuBM8
CuBM9

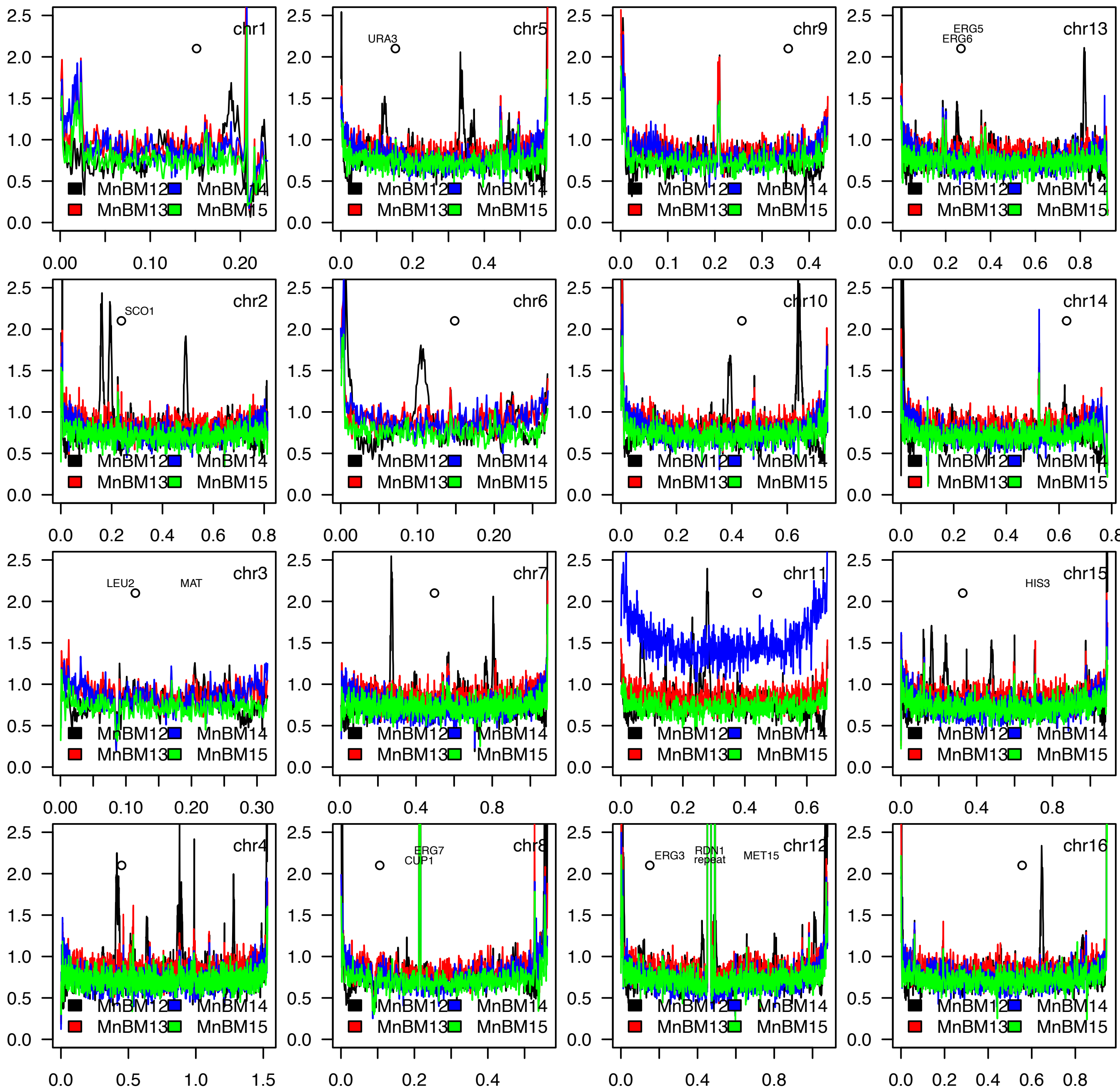

chr1
chr5
URA3
chr9
chr13
ERG5
ERG6
chr2
SCO1
chr6
chr10
chr14
chr3
LEU2
MAT
chr7
chr11
chr15
HIS3
chr4
chr8
ERG7
CUP1
chr12
ERG3
RDN1
repeat
MET15
chr16
MnBM12
MnBM14
MnBM13
MnBM15

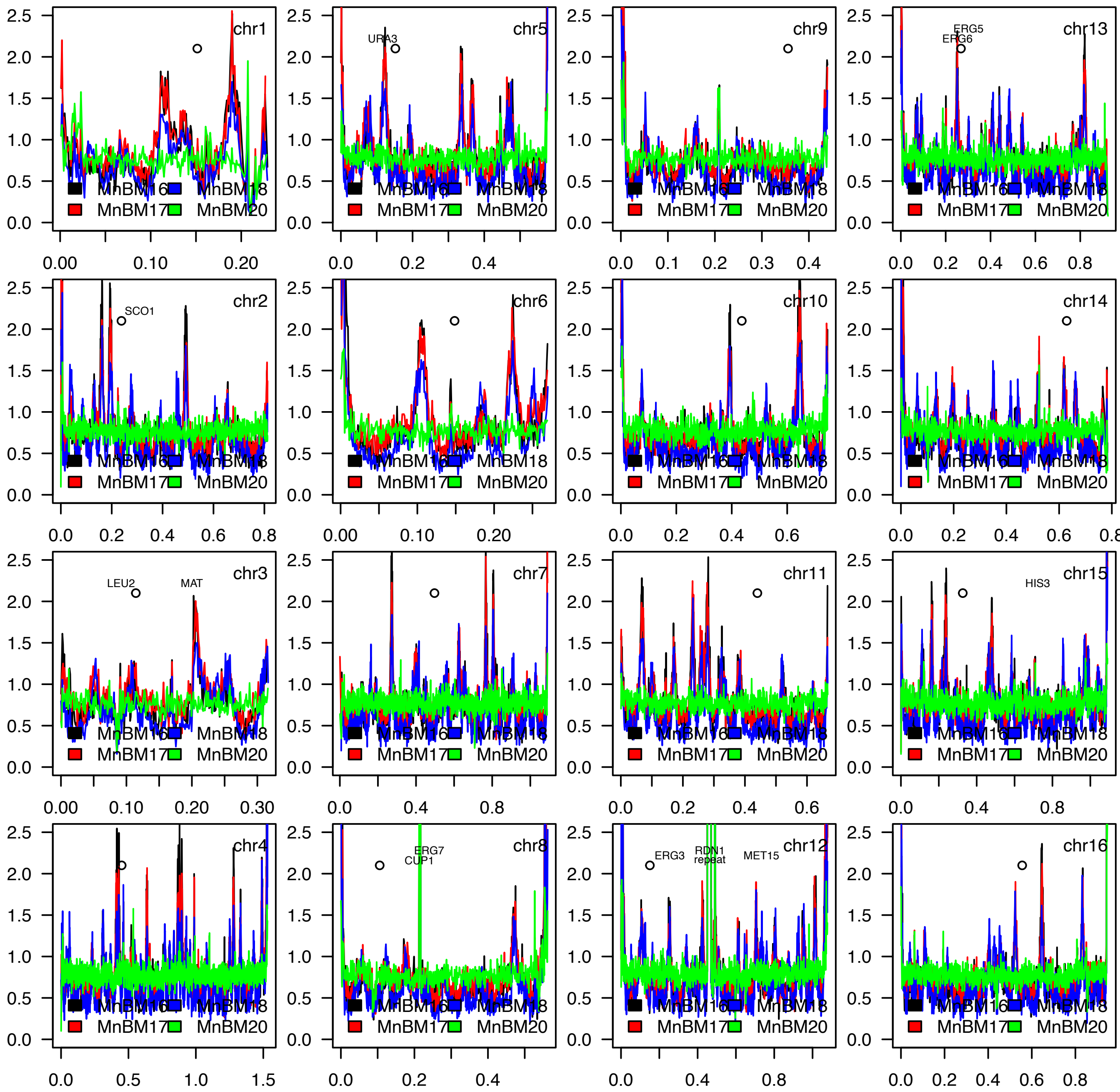
chr1
chr5
URA3
chr9
chr13
ERG5
ERG6
chr2
SCO1
chr6
chr10
chr14
chr3
LEU2
MAT
chr7
chr11
chr15
HIS3
chr4
chr8
ERG7
CUP1
chr12
ERG3
RDN1
repeat
MET15
chr16
MnBM16
MnBM18
MnBM17
MnBM20
0.0
0.5
1.0
1.5
2.0
2.5

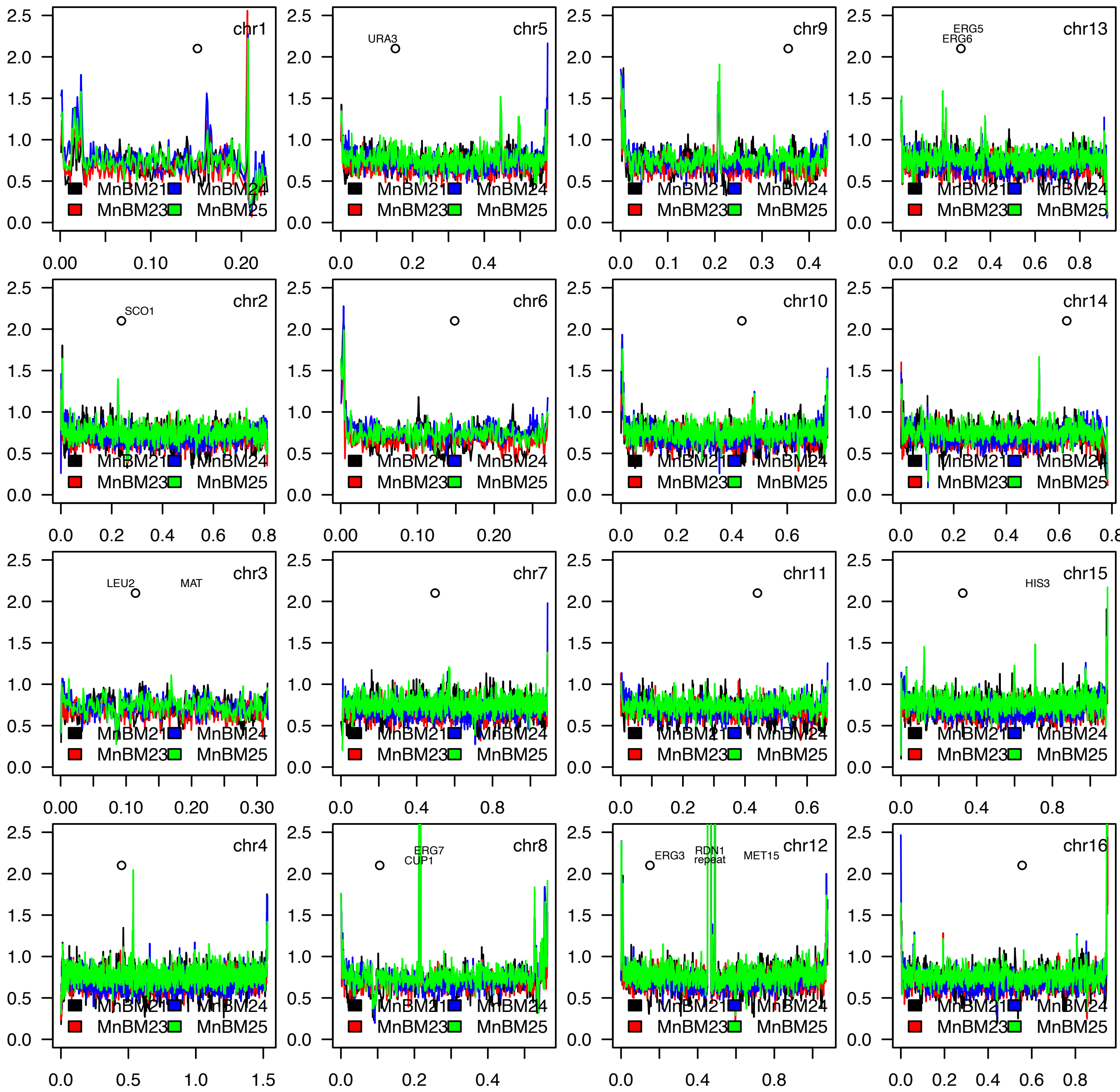
chr1
chr5
URA3
chr9
chr13
ERG5
ERG6
chr2
SCO1
chr6
chr10
chr14
chr3
LEU2
MAT
chr7
chr11
chr15
HIS3
chr4
chr8
ERG7
CUP1
chr12
ERG3
RDN1
repeat
MET15
chr16
MnBM21
MnBM24
MnBM23
MnBM25

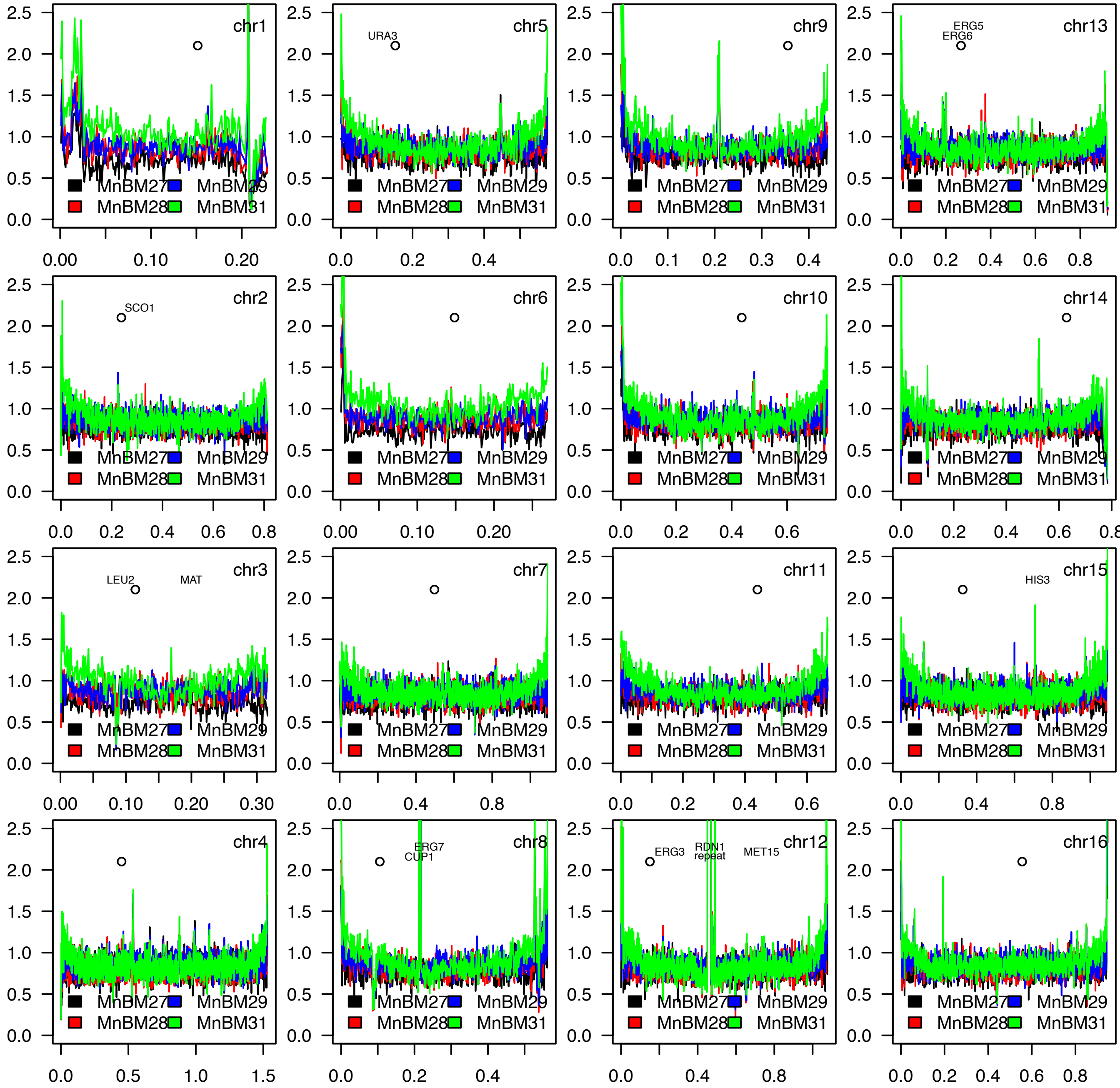
chr1
chr5
URA3
chr9
chr13
ERG5
ERG6
chr2
SCO1
chr6
chr10
chr14
chr3
LEU2
MAT
chr7
chr11
chr15
HIS3
chr4
chr8
ERG7
CUP1
chr12
ERG3
RDN1
repeat
MET15
chr16
MnBM27
MnBM29
MnBM28
MnBM31

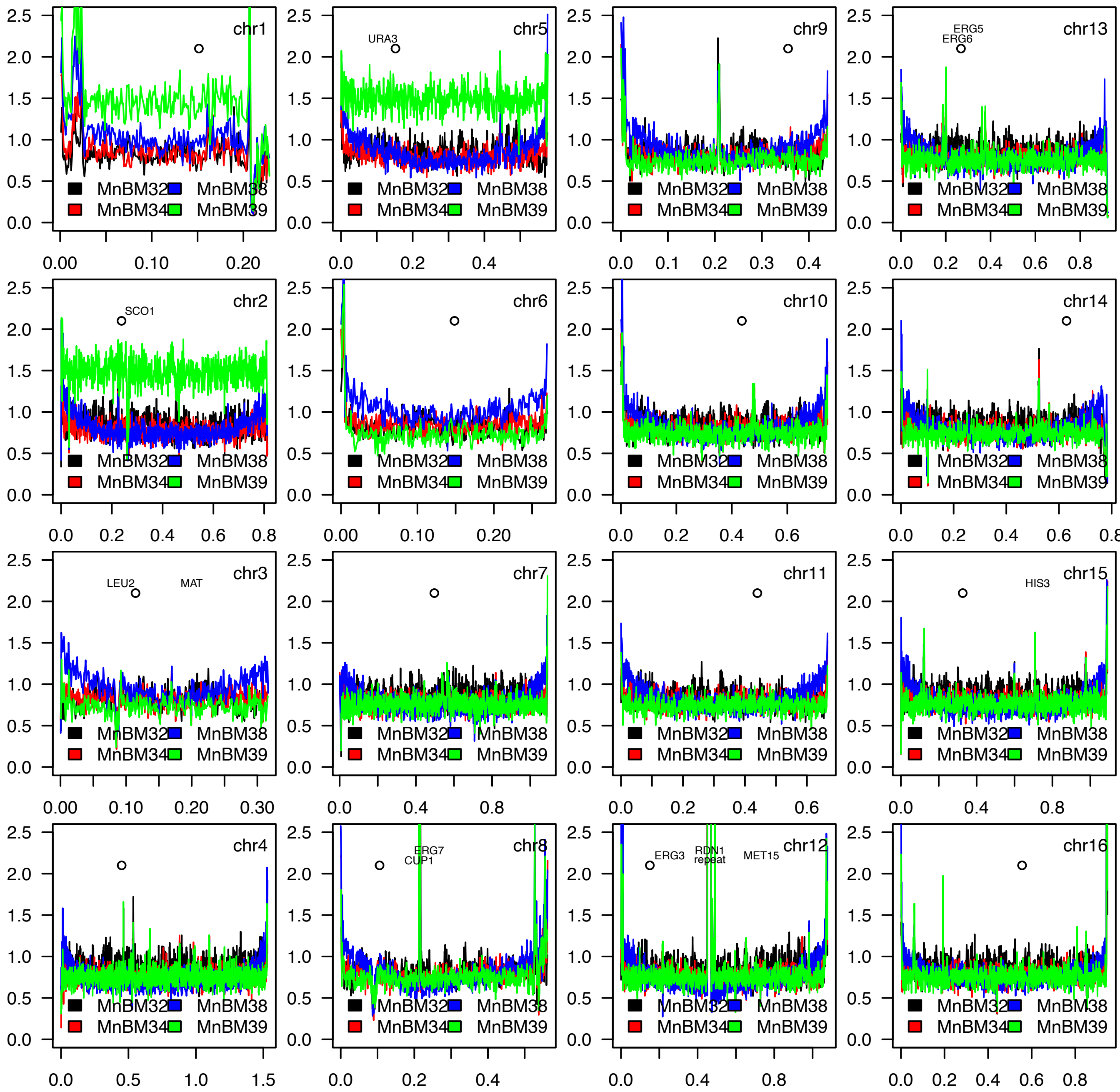
chr1
chr5
URA3
chr9
chr13
ERG5
ERG6
chr2
SCO1
chr6
chr10
chr14
chr3
LEU2
MAT
chr7
chr11
chr15
HIS3
chr4
chr8
ERG7
CUP1
chr12
ERG3
RDN1
repeat
MET15
chr16
MnBM32
MnBM38
MnBM34
MnBM39

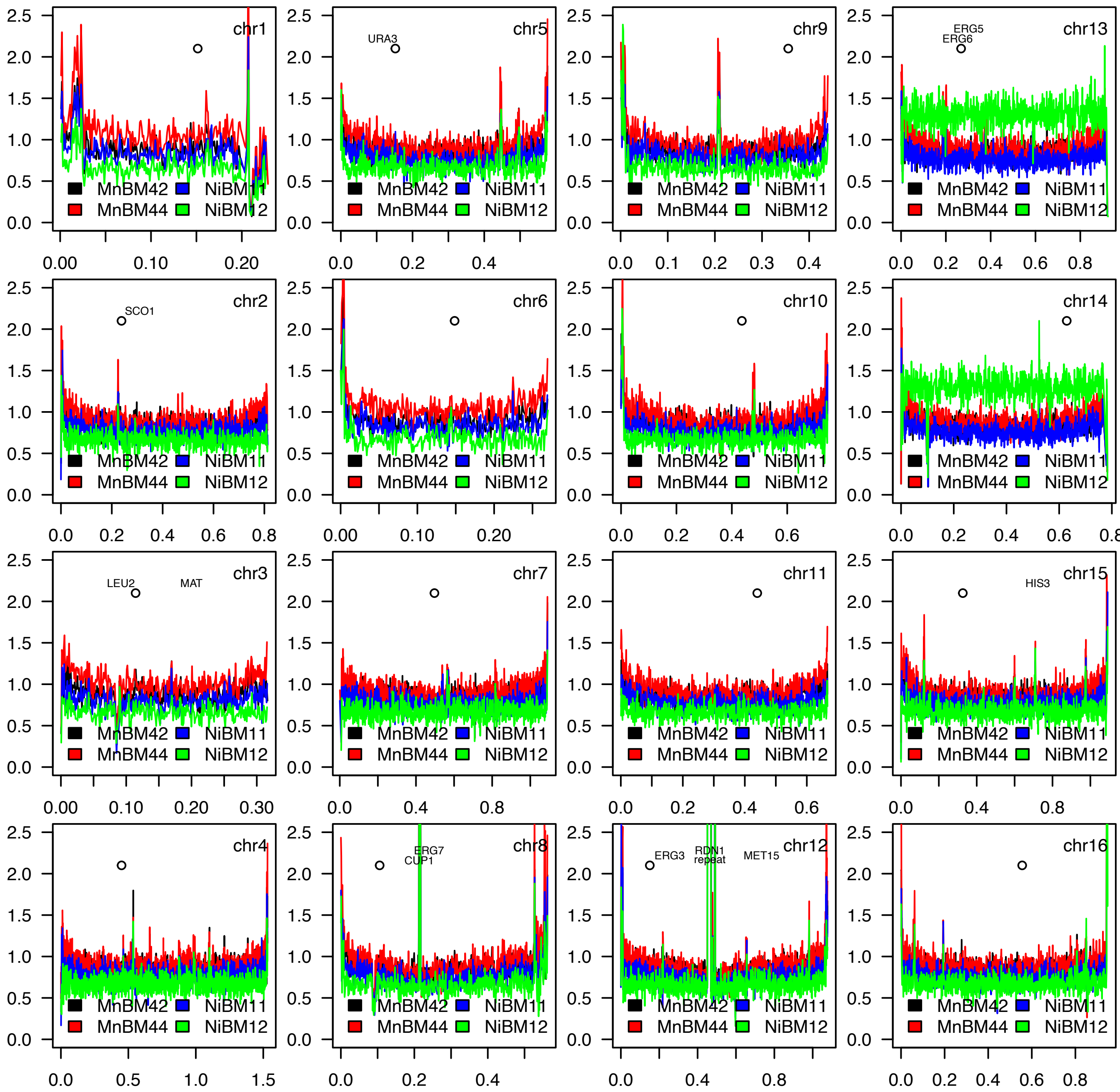
chr1
chr5
URA3
chr9
chr13
ERG5
ERG6
chr2
SCO1
chr6
chr10
chr14
chr3
LEU2
MAT
chr7
chr11
chr15
HIS3
chr4
chr8
ERG7
CUP1
chr12
ERG3
RDN1
repeat
MET15
chr16
MnBM42
NiBM11
MnBM44
NiBM12

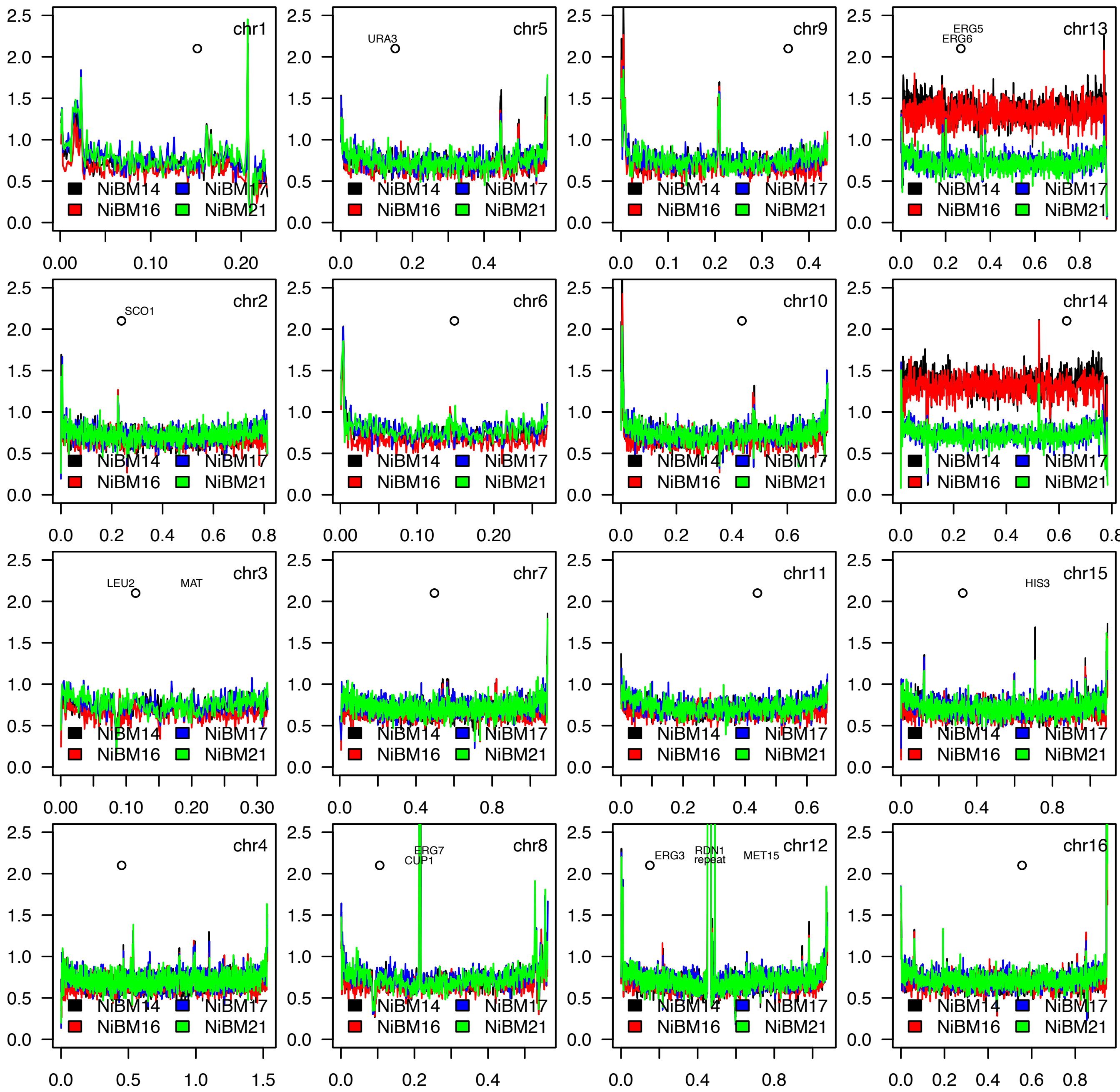
chr1
chr5
URA3
chr9
chr13
ERG5
ERG6
chr2
SCO1
chr6
chr10
chr14
chr3
LEU2
MAT
chr7
chr11
chr15
HIS3
chr4
chr8
ERG7
CUP1
chr12
ERG3
RDN1
repeat
MET15
chr16
NiBM14
NiBM17
NiBM16
NiBM21

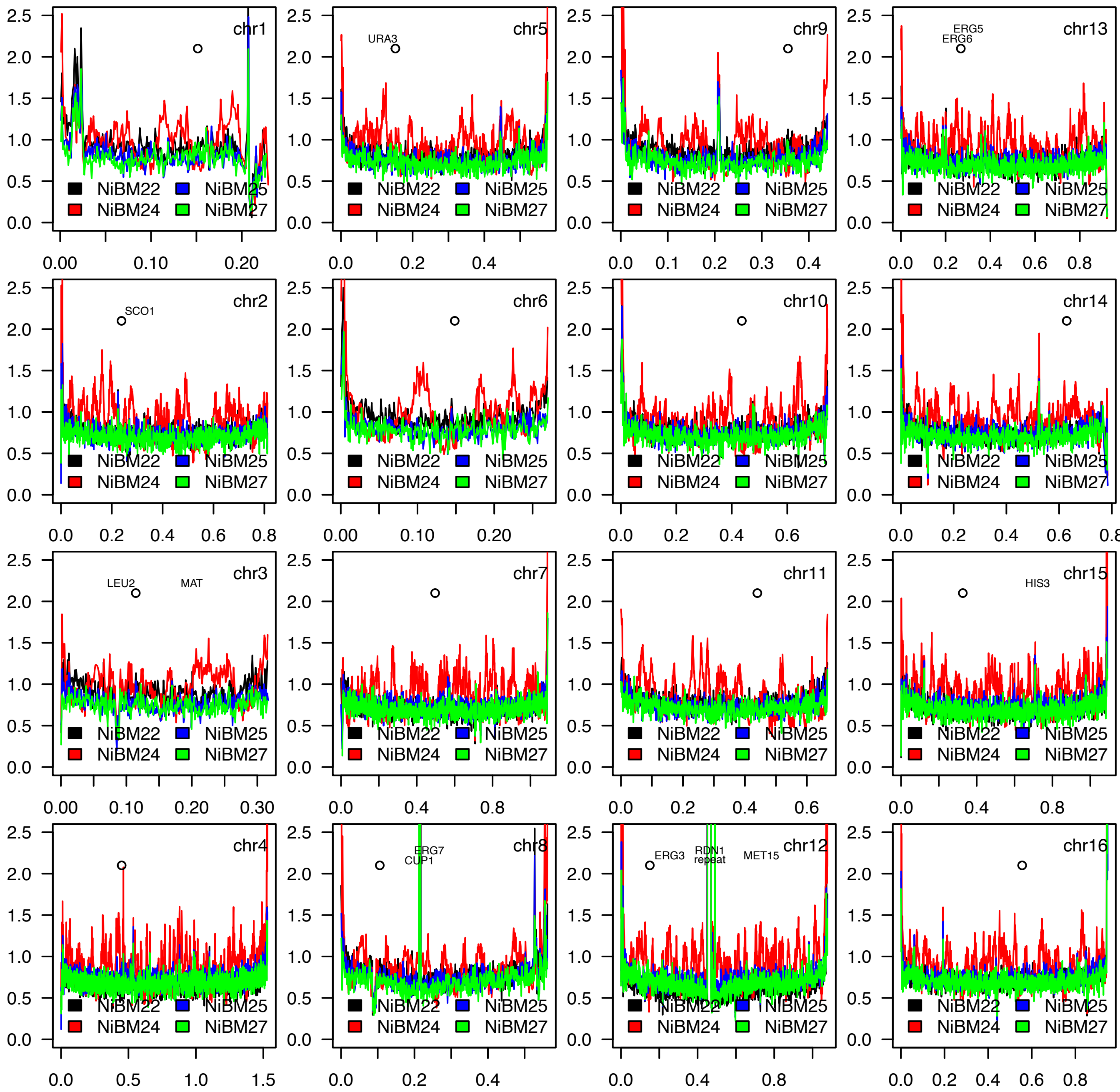
chr1
chr5
URA3
chr9
chr13
ERG5
ERG6
chr2
SCO1
chr6
chr10
chr14
chr3
LEU2
MAT
chr7
chr11
chr15
HIS3
chr4
chr8
ERG7
CUP1
chr12
ERG3
RDN1
repeat
MET15
chr16
NiBM22
NiBM24
NiBM25
NiBM27

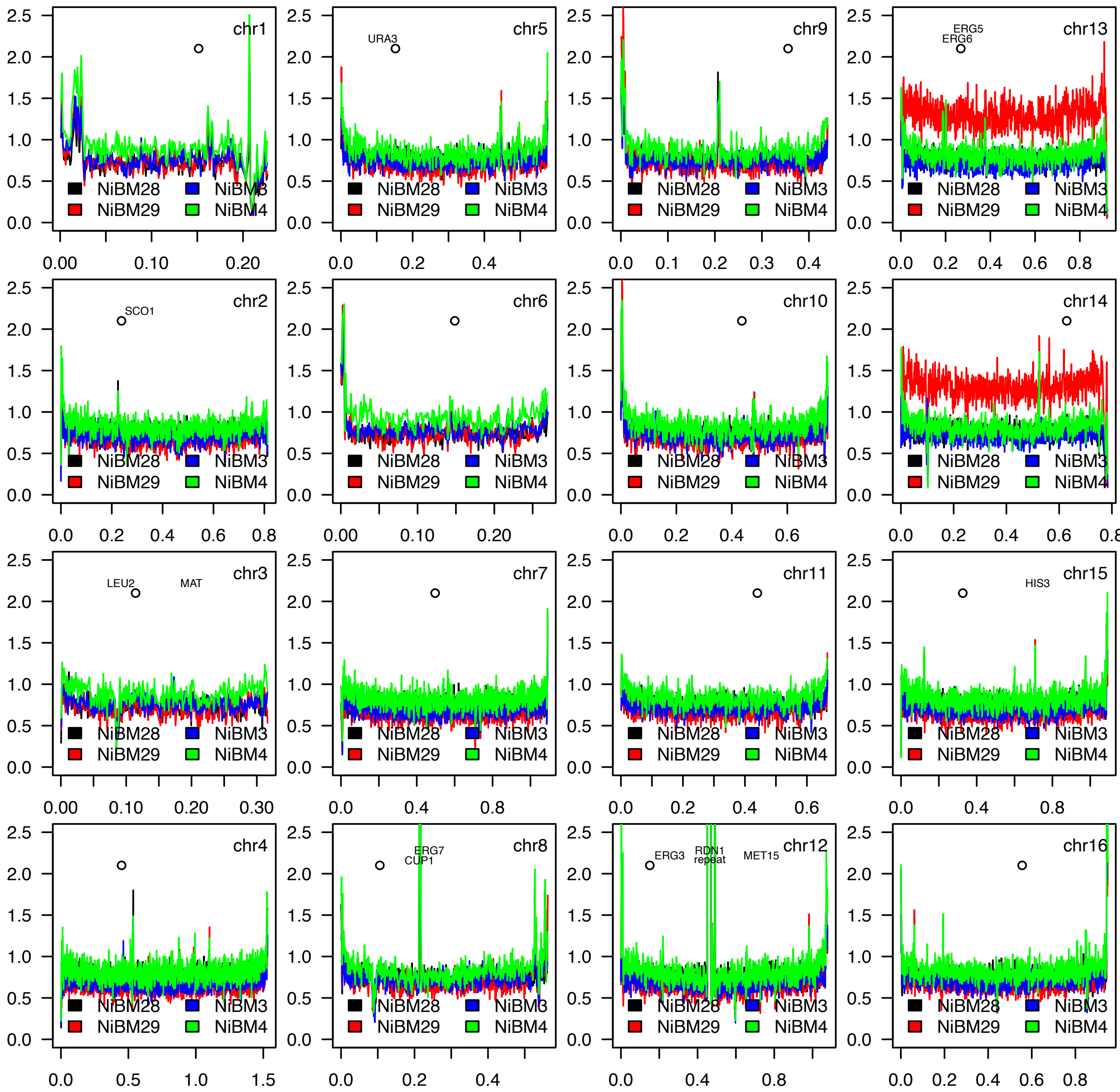
chr1
chr5
URA3
chr9
chr13
ERG5
ERG6
chr2
SCO1
chr6
chr10
chr14
chr3
LEU2
MAT
chr7
chr11
chr15
HIS3
chr4
chr8
ERG7
CUP1
chr12
ERG3
RDN1
repeat
MET15
chr16
NiBM28
NiBM3
NiBM29
NiBM4

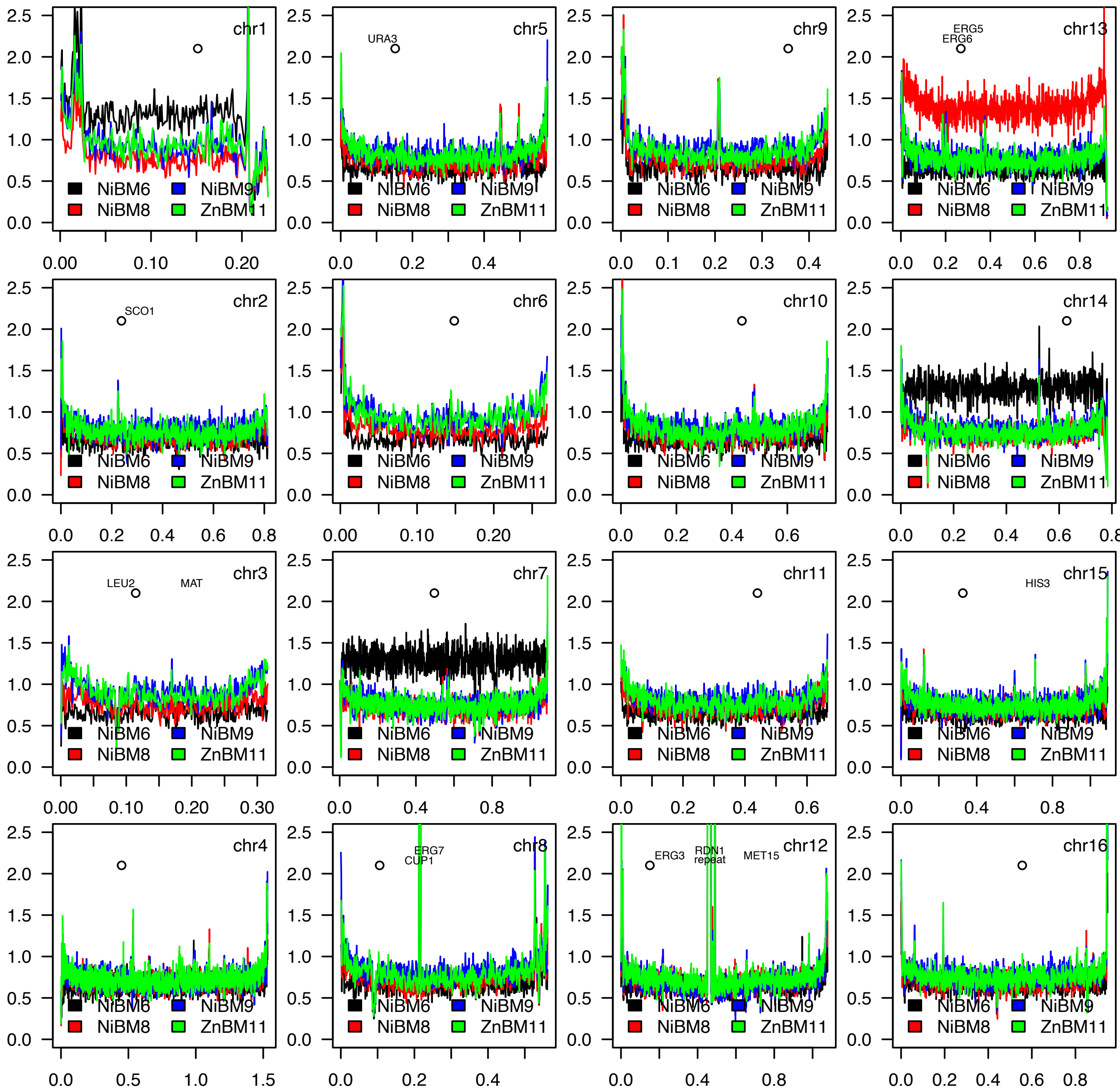
chr1
chr5
URA3
chr9
chr13
ERG5
ERG6
chr2
SCO1
chr6
chr10
chr14
chr3
LEU2
MAT
chr7
chr11
chr15
HIS3
chr4
chr8
ERG7
CUP1
chr12
ERG3
RDN1
repeat
MET15
chr16
NiBM6
NiBM9
NiBM8
ZnBM11

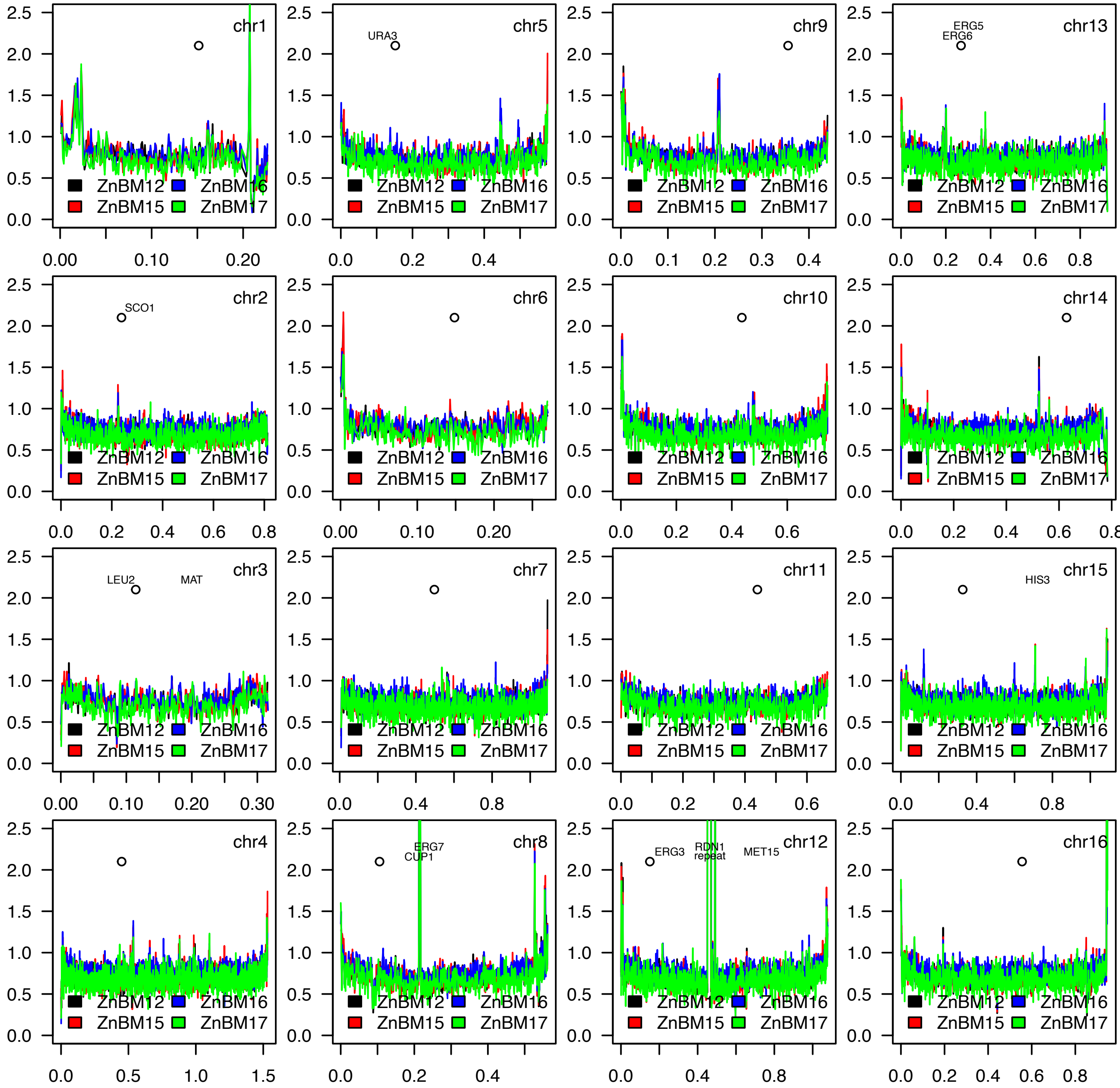
chr1
ZnBM12 ZnBM16
ZnBM15 ZnBM17
chr5
URA3
ZnBM12 ZnBM16
ZnBM15 ZnBM17
chr9
ZnBM12 ZnBM16
ZnBM15 ZnBM17
chr13
ERG5
ERG6
ZnBM12 ZnBM16
ZnBM15 ZnBM17
chr2
SCO1
ZnBM12 ZnBM16
ZnBM15 ZnBM17
chr6
ZnBM12 ZnBM16
ZnBM15 ZnBM17
chr10
ZnBM12 ZnBM16
ZnBM15 ZnBM17
chr14
ZnBM12 ZnBM16
ZnBM15 ZnBM17
chr3
LEU2
MAT
ZnBM12 ZnBM16
ZnBM15 ZnBM17
chr7
ZnBM12 ZnBM16
ZnBM15 ZnBM17
chr11
ZnBM12 ZnBM16
ZnBM15 ZnBM17
chr15
HIS3
ZnBM12 ZnBM16
ZnBM15 ZnBM17
chr4
ZnBM12 ZnBM16
ZnBM15 ZnBM17
chr8
ERG7
CUP1
ZnBM12 ZnBM16
ZnBM15 ZnBM17
chr12
ERG3
RDN1
repeat
MET15
ZnBM12 ZnBM16
ZnBM15 ZnBM17
chr16
ZnBM12 ZnBM16
ZnBM15 ZnBM17
0.0 0.5 1.0 1.5 2.0 2.5

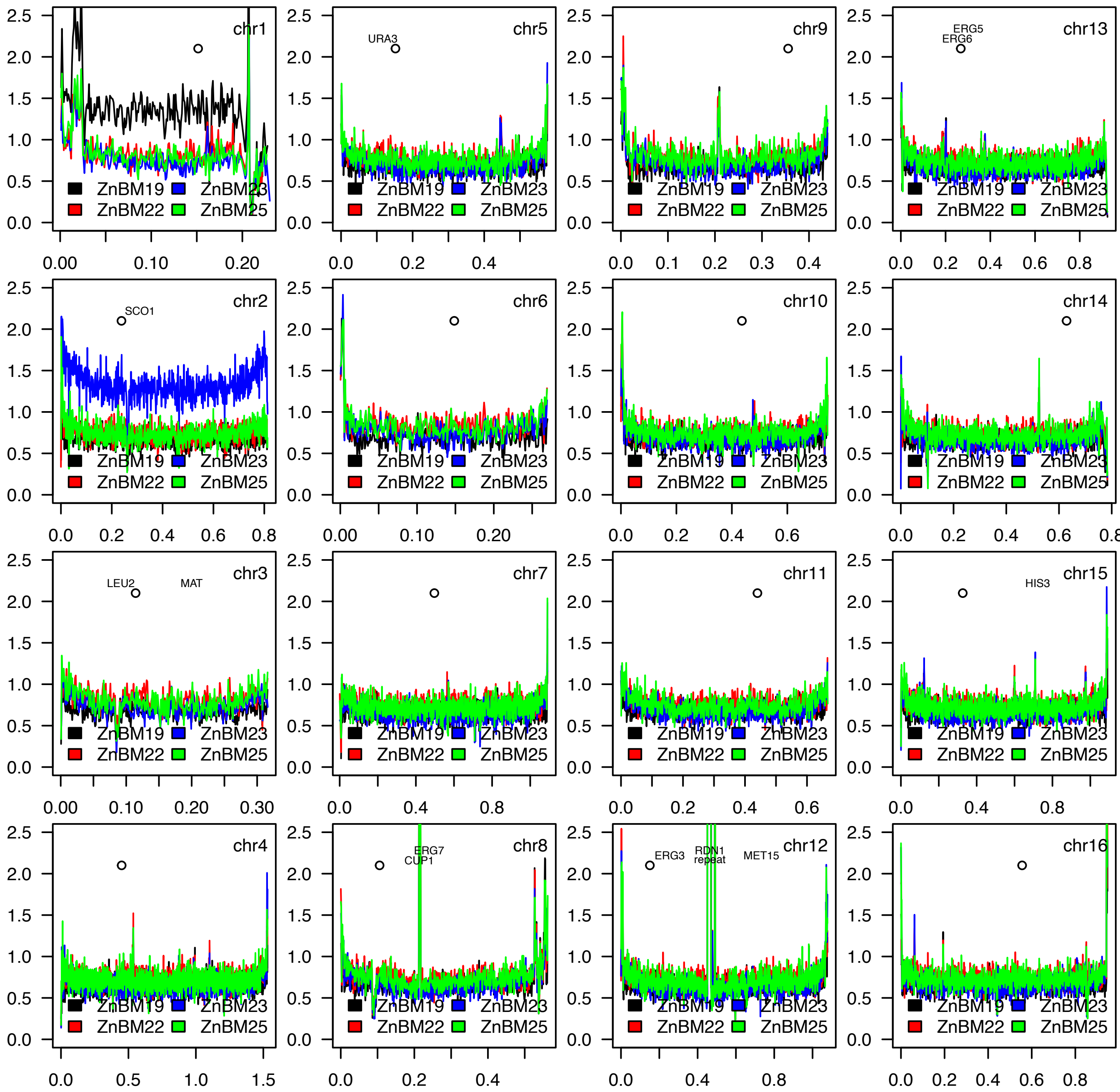
chr1
ZnBM19 ZnBM23
ZnBM22 ZnBM25
chr5
URA3
chr9
chr13
ERG5
ERG6
chr2
SCO1
chr6
chr10
chr14
chr3
LEU2
MAT
chr7
chr11
chr15
HIS3
chr4
chr8
ERG7
CUP1
chr12
ERG3
RDN1
repeat
MET15
chr16

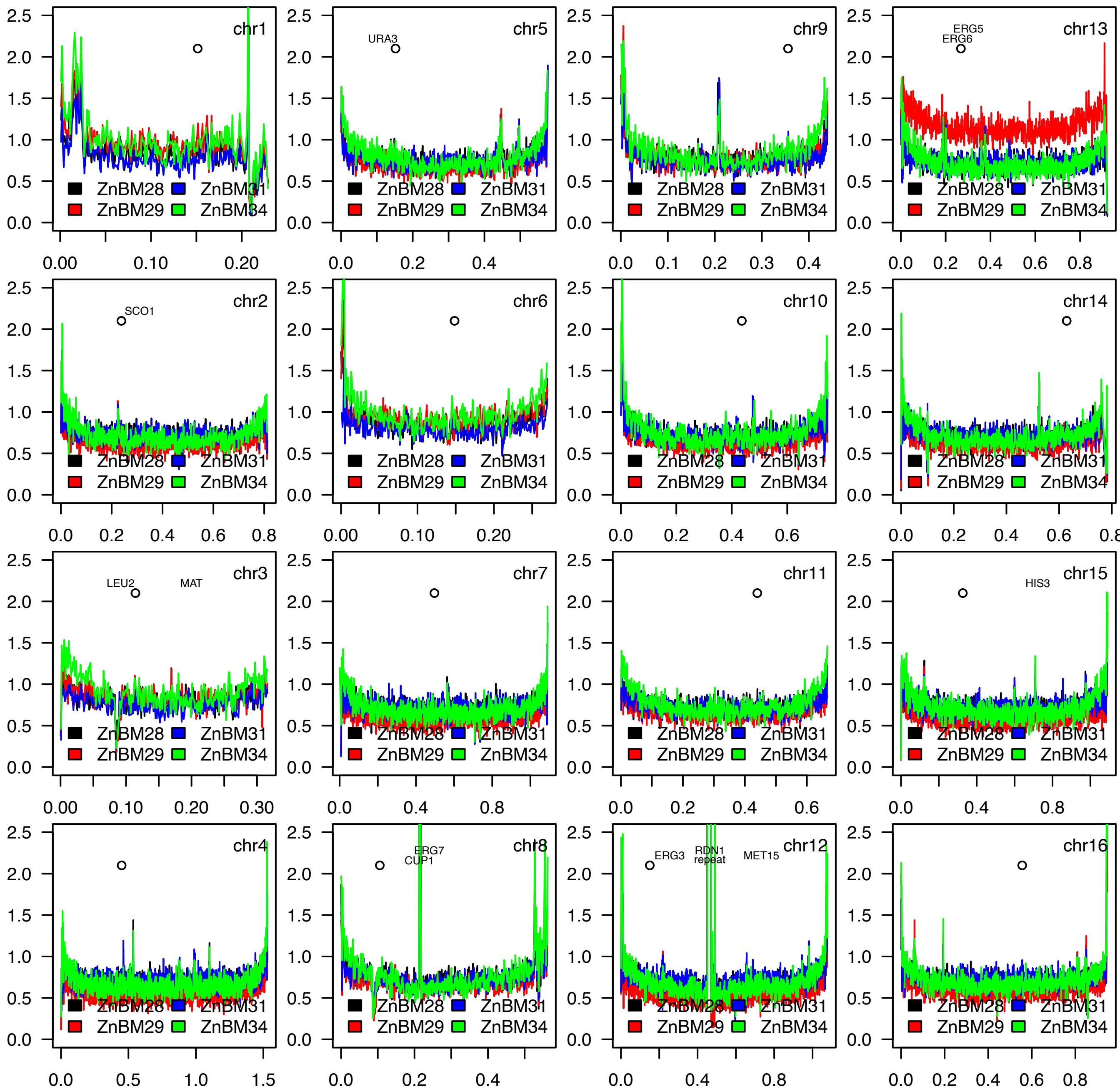
chr1
chr5
URA3
chr9
chr13
ERG5
ERG6
chr2
SCO1
chr6
chr10
chr14
chr3
LEU2
MAT
chr7
chr11
chr15
HIS3
chr4
chr8
ERG7
CUP1
chr12
ERG3
RDN1
repeat
MET15
chr16
ZnBM28
ZnBM31
ZnBM29
ZnBM34

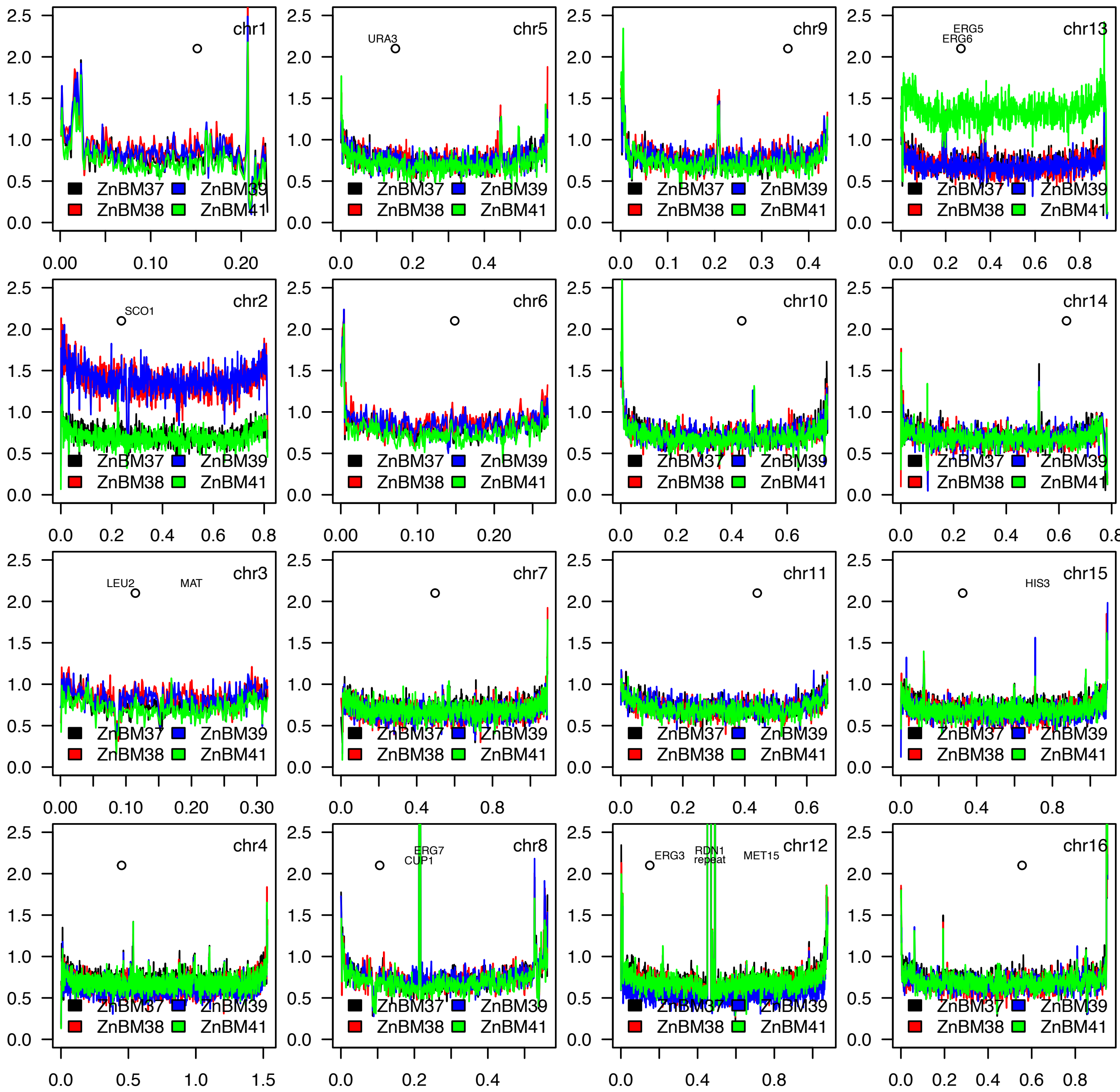
chr1
chr5
URA3
chr9
chr13
ERG5
ERG6
chr2
SCO1
chr6
chr10
chr14
chr3
LEU2
MAT
chr7
chr11
chr15
HIS3
chr4
chr8
ERG7
CUP1
chr12
ERG3
RDN1
repeat
MET15
chr16
ZnBM37
ZnBM38
ZnBM39
ZnBM41

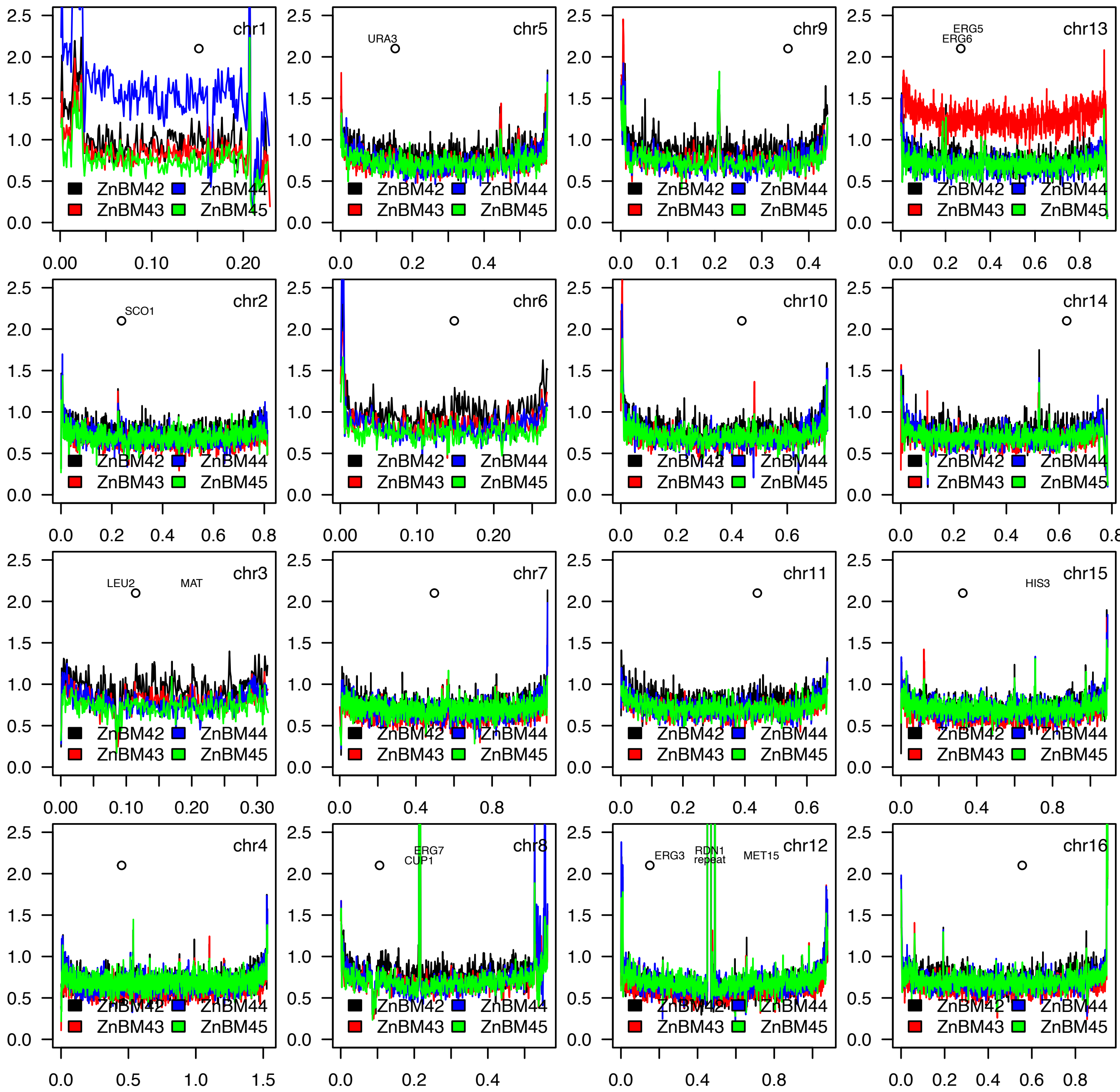

chr1
chr5
URA3
chr9
chr13
ERG5
ERG6
chr2
SCO1
chr6
chr10
chr14
chr3
LEU2
MAT
chr7
chr11
chr15
HIS3
chr4
chr8
ERG7
CUP1
chr12
ERG3
RDN1
repeat
MET15
chr16
ZnBM42
ZnBM43
ZnBM44
ZnBM45

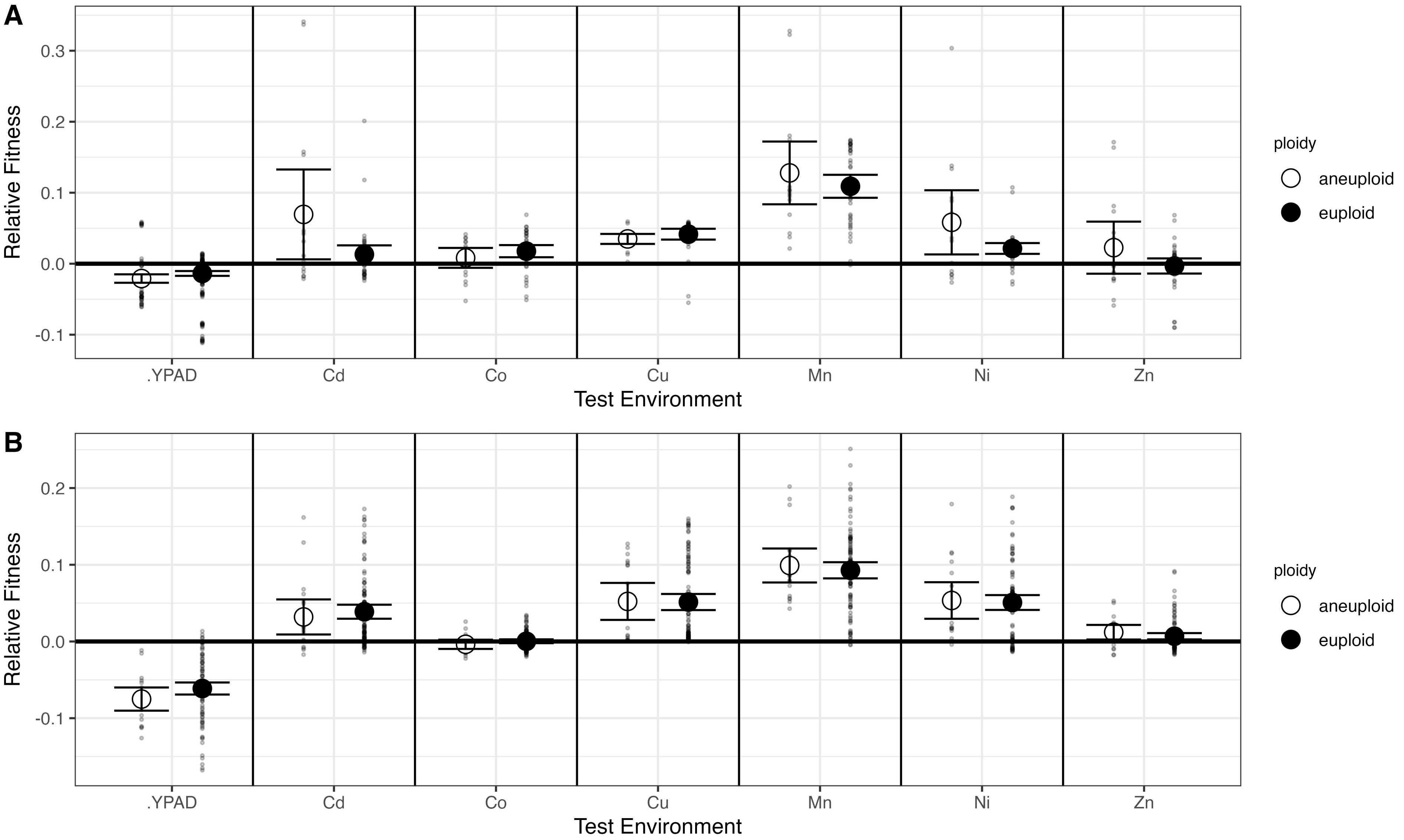
A
B
Relative Fitness
0.3
0.2
0.1
0.0
-0.1
.YPAD
Cd
Co
Cu
Mn
Ni
Zn
Test Environment
ploidy
aneuploid
euploid

A

| Chromosome | W303 | Cd.37 | Cd.39 | Cd.44 | Cd.45 | Co.2 | Mn.14 | Mn.39 | Ni.6 | Ni.8 | Ni.12 | Ni.14 | Ni.16 | Ni.29 | Zn.19 | Zn.23 | Zn.29 | Zn.38 | Zn.39 | Zn.41 | Zn.43 | Zn.44 | Zn.47 |
|---|---|---|---|---|---|---|---|---|---|---|---|---|---|---|---|---|---|---|---|---|---|---|---|
| I | 1 | 1.0 | 1.0 | 1.0 | 1.4 | 0.9 | 1.1 | 1.6 | 1.7 | 1.0 | 0.9 | 0.9 | 0.9 | 1.0 | 1.8 | 1.1 | 1.4 | 1.2 | 1.2 | 1.0 | 1.2 | 2.2 | 0.6 |
| II | 1 | 1.7 | 1.7 | 1.6 | 1.0 | 1.0 | 1.0 | 1.8 | 0.9 | 1.0 | 0.9 | 0.9 | 0.9 | 0.9 | 1.0 | 2.0 | 1.0 | 1.9 | 2.0 | 1.0 | 1.0 | 1.0 | 0.4 |
| III | 1 | 1.7 | 1.0 | 1.0 | 1.3 | 0.9 | 1.1 | 0.8 | 0.9 | 1.0 | 0.9 | 0.9 | 0.9 | 0.9 | 0.9 | 1.0 | 1.3 | 1.1 | 1.1 | 0.9 | 1.1 | 1.1 | 0.4 |
| IV | 1 | 1.6 | 1.6 | 1.0 | 0.9 | 1.0 | 0.9 | 0.9 | 0.9 | 1.0 | 1.0 | 0.9 | 0.9 | 0.9 | 1.0 | 1.0 | 0.9 | 0.9 | 0.9 | 0.9 | 0.9 | 1.0 | 0.4 |
| V | 1 | 0.9 | 0.9 | 1.0 | 1.1 | 1.0 | 1.0 | 1.7 | 0.9 | 1.0 | 0.9 | 0.9 | 0.9 | 1.0 | 0.9 | 1.0 | 1.0 | 1.0 | 1.0 | 0.9 | 1.0 | 1.0 | 0.8 |
| VI | 1 | 1.1 | 1.0 | 1.0 | 1.4 | 0.9 | 1.1 | 0.8 | 0.9 | 1.0 | 0.9 | 0.9 | 0.9 | 1.0 | 0.9 | 1.1 | 1.4 | 1.2 | 1.1 | 1.0 | 1.2 | 1.1 | 0.6 |
| VII | 1 | 0.8 | 0.9 | 1.0 | 1.0 | 1.0 | 1.0 | 0.9 | 1.8 | 1.0 | 0.9 | 0.9 | 0.9 | 0.9 | 1.0 | 1.0 | 0.9 | 0.9 | 0.9 | 1.0 | 0.9 | 1.0 | 0.8 |
| VIII | 1 | 0.8 | 0.9 | 1.0 | 1.1 | 1.0 | 1.0 | 0.9 | 0.9 | 1.0 | 0.9 | 0.9 | 0.9 | 0.9 | 1.0 | 1.0 | 1.0 | 1.0 | 1.0 | 0.9 | 1.0 | 1.0 | 0.8 |
| IX | 1 | 0.8 | 0.9 | 1.2 | 1.2 | 0.9 | 1.0 | 0.9 | 0.9 | 1.0 | 0.9 | 1.0 | 0.9 | 1.0 | 0.9 | 1.0 | 1.2 | 1.1 | 1.1 | 1.0 | 1.1 | 1.1 | 0.8 |
| X | 1 | 0.8 | 0.9 | 1.0 | 1.0 | 1.0 | 1.0 | 0.9 | 0.9 | 1.0 | 0.9 | 1.0 | 0.9 | 1.0 | 1.0 | 1.0 | 1.0 | 1.0 | 1.0 | 1.0 | 1.0 | 1.0 | 1.6 |
| XI | 1 | 0.8 | 0.9 | 1.0 | 1.1 | 1.0 | 2.0 | 0.9 | 0.9 | 1.0 | 0.9 | 0.9 | 0.9 | 0.9 | 1.0 | 1.0 | 1.0 | 1.0 | 1.0 | 1.0 | 1.0 | 1.0 | 0.8 |
| XII | 1 | 0.8 | 0.8 | 0.7 | 0.9 | 0.8 | 0.8 | 0.9 | 0.7 | 0.7 | 0.7 | 0.7 | 0.7 | 0.7 | 1.1 | 0.7 | 0.8 | 0.8 | 0.8 | 0.9 | 0.7 | 0.8 | 1.0 |
| XIII | 1 | 0.8 | 0.9 | 1.0 | 1.0 | 1.9 | 1.0 | 0.9 | 0.9 | 1.9 | 1.8 | 1.8 | 1.8 | 1.8 | 1.0 | 1.0 | 1.8 | 1.0 | 1.0 | 1.9 | 1.9 | 1.0 | 1.6 |
| XIV | 1 | 0.8 | 0.9 | 1.1 | 1.0 | 1.0 | 1.0 | 0.9 | 1.8 | 1.0 | 1.8 | 1.8 | 1.8 | 1.8 | 1.0 | 1.0 | 1.0 | 1.0 | 1.0 | 1.0 | 1.0 | 1.0 | 0.8 |
| XV | 1 | 0.8 | 0.8 | 1.0 | 1.0 | 1.0 | 1.0 | 0.9 | 0.9 | 1.0 | 1.0 | 0.9 | 0.9 | 0.9 | 1.0 | 1.0 | 0.9 | 0.9 | 0.9 | 1.0 | 0.9 | 1.0 | 0.8 |
| XVI | 1 | 0.8 | 0.9 | 1.1 | 1.0 | 1.0 | 1.0 | 0.9 | 0.9 | 1.0 | 1.0 | 0.9 | 0.9 | 0.9 | 1.0 | 1.0 | 0.9 | 0.9 | 1.0 | 1.0 | 1.0 | 1.0 | 0.8 |

B

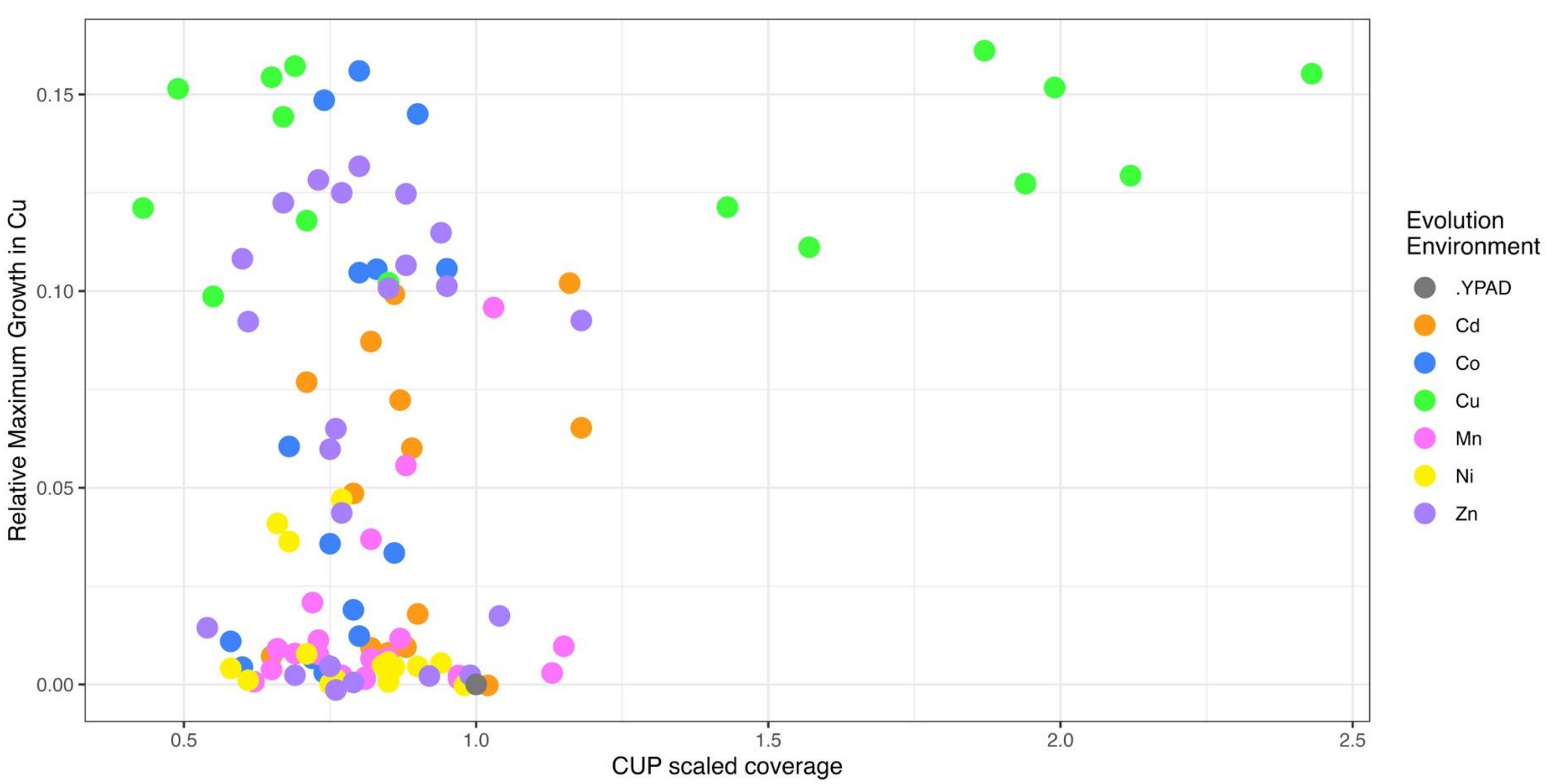

### Distribution on the deep-well plate of copper mutations

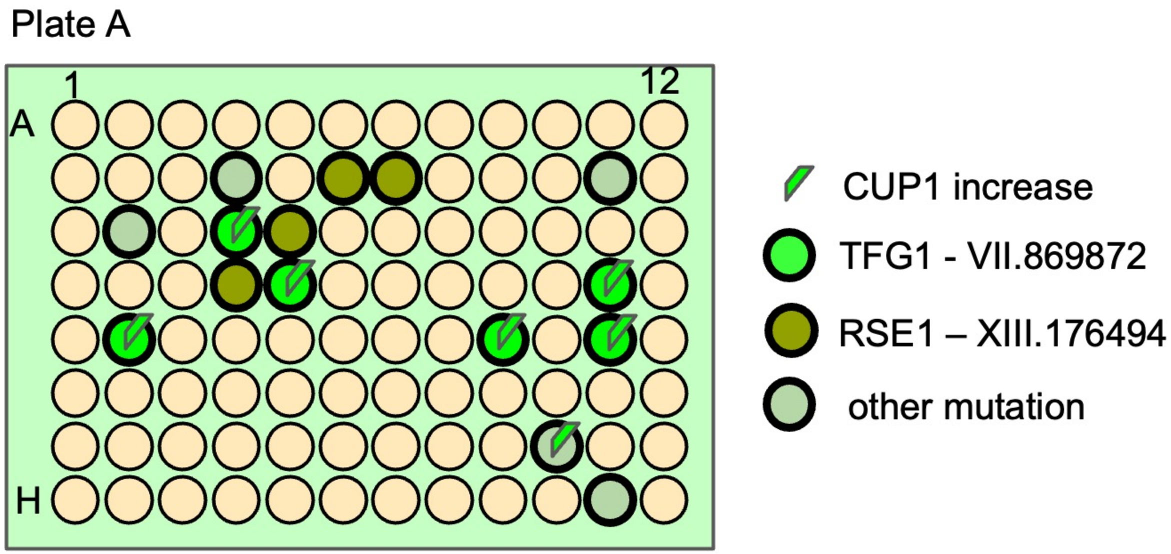


| | | |
|---|---|---|
| CuBM10 | 4d | RSE1, NGG1 |
| CuBM11 | 5d | TFG1, NGG1 |
| CuBM12 | 6b | RSE1, NGG1 |
| CuBM13 | 7b | RSE1, NGG1 |
| CuBM14 | 9e | TFG1, BUL1, FIG4, PYK2, NGG1 |
| CuBM15 | 10g | COQ1 |
| CuBM17 | 11d | TFG1, NGG1 |
| CuBM18 | 11e | ABP1, ROG1, TFG1, PRP2, NGG1 |
| CuBM3 | 11h | PMA1 |
| CuBM4 | 5c | RSC1, NGG1, RSE1 |
| CuBM6 | 2c | DNF1, ATG2 |
| CuBM7 | 2e | BLM10, TFG1, NGG1 |
| CuBM9 | 4c | MMS4, TFG1, KSP1, NGG1 |

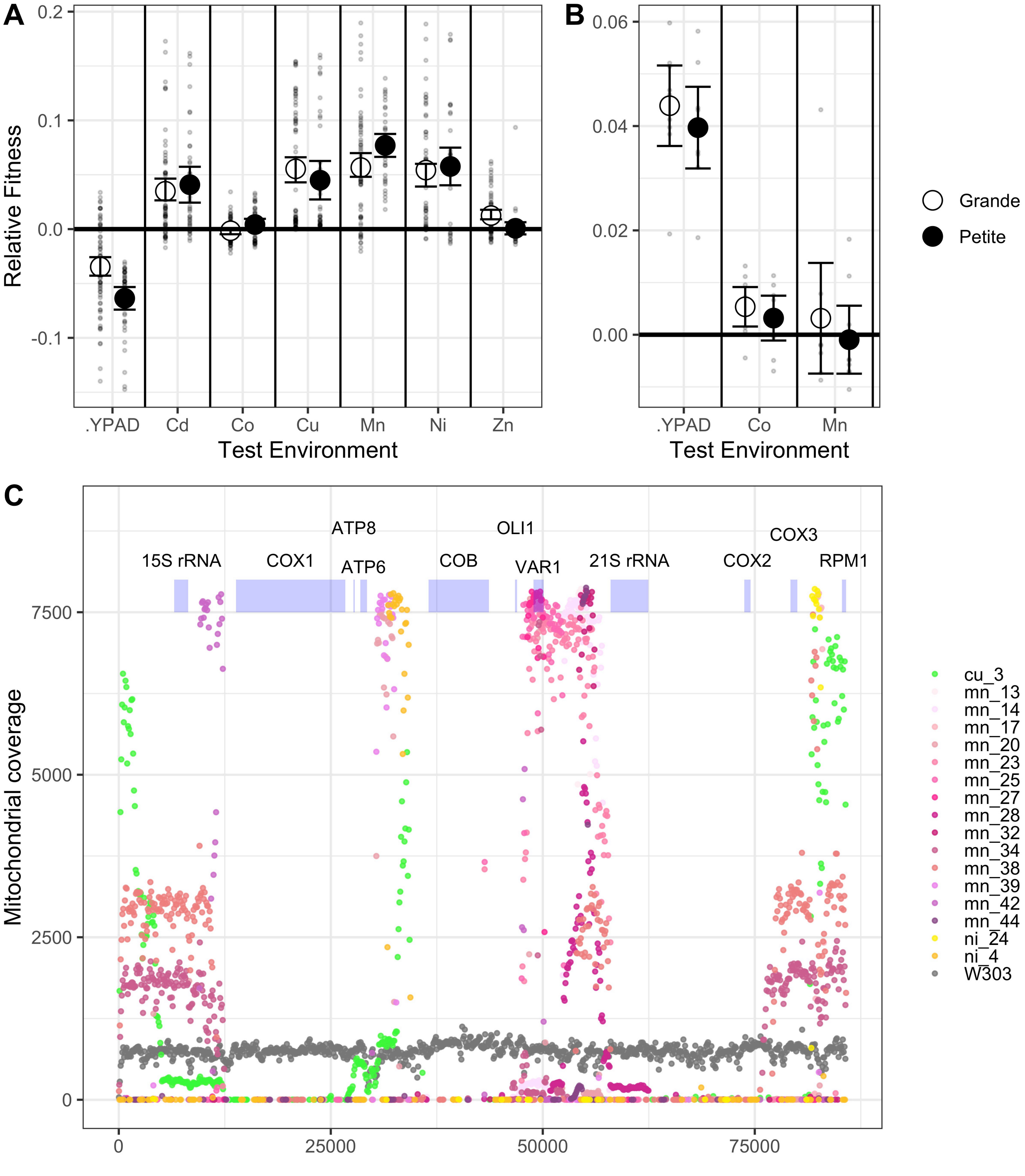
A
B
C
Relative Fitness
Test Environment
.YPAD
Cd
Co
Cu
Mn
Ni
Zn
Grande
Petite
Mitochondrial coverage
15S rRNA
COX1
ATP8
ATP6
COB
OLI1
VAR1
21S rRNA
COX2
COX3
RPM1
cu_3
mn_13
mn_14
mn_17
mn_20
mn_23
mn_25
mn_27
mn_28
mn_32
mn_34
mn_38
mn_39
mn_42
mn_44
ni_24
ni_4
W303
