## Appendix 1 for "Evolution of cross-tolerance to metals in yeast"

GITHUB address: https://github.com/joelkcamp/crossTolerance_bazzicalupoEtAl

#### 1 Tolerance curves

To decide which concentrations to use in our experiment we characterized the tolerance curves of the yeast strain W303 and the BY4741. We used W303 in our evolution experiment. BY4741 had been used previously and we characterized its tolerance curves to several metals.

R script to extract growth curve information: 1_shape_clean.R. This script calculates the spline using the max_growth_rate.R and bioscreen_functions.R, and summarizes the findings into tolerance curves.

#### 2 Reciprocal Transplant Growth Curves

Once the yeast lines were evolved, we performed a reciprocal transplant experiment. Similar to the tolerance curves, this script extracts the information for all the bioscreen OD readings:

2_evo_test.Rmd

We inspected growth curves and matched the expects ancestor in YPAD to perform and match our expectation and controls were clean.

*Compromised batch* folder contains a machine (2 bioscreen plates) that did not show growth in the ancestor in permissive media and we excluded it from further analyses because it's not clear if it worked properly.

#### 3 Reciprocal Transplant Cross Tolerance

The maximum growth measurements were then subtracted from the ancestral maximum growth rate in each environment to have a relative growth rate across the whole experiment. We also subtracted the maximum growth rate of the yeast tested in each environment relative to their evolution environment. Script: 3_cross_tol.Rmd

We also tested the relationship between genotype quality and environment quality including and excluding the evolution environment to explore the possibility that the harshness of the environment has an impact on the evolvability of cross-tolerance and the relationship between generalist and specialists.

For each pair of environments we also ran ANOVAs and show if the evolution vs. test environments are significantly different and if the interaction term is significant. These statistics help us summarize the level of local adaptation and which environments are more likely to lead to local adaptation.

### 4 Genome Analysis

We sequenced the genomes of 109 lines + ancestor. Steps from .fastq files to .vcf variant calling are included in folder: /4_bash

### 5 Petite Analysis

We recorded petite phenotype and we looked at the performance of the lines in each metal. There was some hint that perhaps it may be advantageous and adaptive to be petite in cobalt and manganese where almost all lines evolved a petite phenotype. We tested petite and grande lines from a different experiment using the same genetic background strain and showed that it is not the case and being petite in itself does not help in those metals.

We also report the coverage of the mitochondrial genome for these lines. We used the mpileup program on bam files (see bash folder for mpileup script). A custom made perl script calculates the coverage for 100 base pair windows (see depth folder for the windows.pl script to calculate the average coverage in windows). The Rmd script uses the output of the perl script to plot the coverage. Most of the manganese and one copper and two nickel lines showed increased coverage in previously described breakpoints. All cobalt lines and most of the copper petite lines completely lost the mitochondrial genome.

Script 5_metal_petite.Rmd

### 6 Chromosome Duplication

We found whole chromosome duplications by estimating coverage with our custom-made perl script calculates the coverage for 1000 base pair windows (see depth folder for the windows.pl script to calculate the average coverage in windows). The Rmd script uses the output of the perl script to plot the coverage of chromosomes. The results are summarized in a table.

We also estimated the relative copy number of the gene CUP by following Gerstein et al. (2015)'s protocol. In folder 'depth'.

Script: 6_aneuploidy.Rmd

### 7 SNP cleaning

We cleaned the vcf file from low quality calls and sites with uncertain alignments. After we systemically cleaned the vcf file (see: 7_snp_analysis.Rmd), we also manually inspected individual SNPs in IGV to confirm they were not ancestral or low coverage. We summarized all the types of mutations: SNP, whole chromosome duplications, CUP coverage, petite into one heat-map showing the amount of cross-tolerance per evolved line.

### 8 GO Terms

The gene annotations for the SNP mutations we are confident in were used to predict cellular component and biological process GO terms in the SGD. We then compared the proportion of each of the GO terms between the genes we found and the whole genome using a Fisher's Exact test to find terms significantly over-represented.

Script: 8_GO_analysis.Rmd

### 9 Predictions

To find ways to predict cross tolerance we used different information about the environment and genes known to matter for metal tolerance to run a random effects model. We first analyzed the potentially predictive data, so the scripts in the following subsections were used first.

Script: 9_predictions.Rmd

#### 9.1 SGD

We calculated an expected amount of cross-tolerance based on the overlap of known genetic mutations conferring cross-tolerance between pairs of metals.

Script: predictions/SGD_metal_resistance_genes_clean.Rmd

#### 9.2 Electropotential and Parts Per Million

We measured the Oxidative Reductive Potential of the media we used in the experiment and the amount in parts per million of the metals to predict how similar the stress between metals is and how much metal the yeast are exposed to in the regular growth medium.

Script: predictions/electrode_potential.Rmd

#### 9.3 US Geological Survey

We downloaded ppm of metals in soils and computed correlations for all pairs of metals to obtain a measure of how often they co-occur in the environment.

Script: predictions/soil_metals.Rmd

### 10 snpEff

SNP mutations were annotated with snpEff and the output of the programme was manipulated with this script. Script: snpEff.Rmd

### 11 Spectra

We compared the types of mutations in our experiment with the mutational spectra measured in Lynch et al. (Table 1 in their manuscript) from their mutation accumulation experiment. We also included results from Gerstein et al. as a control and repeat their test. The mutational spectra were extracted for each metal from the snpEff annotation output. Script: mutationtypes.Rmd
