## Appendix 2 for "Evolution of cross-tolerance to metals in yeast"

### SUP MAT: Evolution of cross-tolerance to metals in yeast

#### Calculating the chance of hitting the same gene once or twice

Given the 6607 genes in the yeast genome, the chance that all genic mutations observed in a metal occur in different genes:

In[1]:=
```
pr1[mutations_] := Product[(6607 - (i - 1))/6607, {i, 1, mutations}] // N
```

The chance that one gene is hit twice (but all other mutations occur in unique genes):

In[2]:=
```
pr2[mutations_] := Total[Table[Product[(6607 - (i - 1))/6607, {i, 1, j - 1}] * ((j - 1)/6607) *
    Product[(6607 - (i - 2))/6607, {i, j + 1, mutations}] // N, {j, 2, mutations}]]
```

The first mutation occurs anywhere. This sums together the chance that the second, third, fourth etc. mutation occurs in one of the previous j-1 genes. Even though the jth gene hits a gene for the second time, all other mutations hit unique genes (contributing to the products).

For some environments, there are so many mutations that it becomes reasonably likely that at least one gene would be hit twice and so we must consider also the probability that there are two genes hit twice:

In[3]:=
```
pr22[mutations_] := Total[
  Flatten[Table[Product[(6607 - (i - 1))/6607, {i, 1, j - 1}] * ((j - 1)/6607) * Product[(6607 - (i - 2))/6607,
      {i, j + 1, k - 1}] * ((k - 1)/6607) * Product[(6607 - (i - 3))/6607, {i, k + 1, mutations}] // N,
    {j, 2, mutations - 1}, {k, j + 1, mutations}]]]
```

We also randomize mutations across the genome and compare the observed distribution of hits per gene (>0) to the randomized distributions.

#### Data

Within the analysis for a particular metal (or for all metals combined), identical mutations (same exact SNP) were eliminated, as were mutations in MnBM14 and MnBM42. Rows do not sum to the “all” column because identical mutations observed in different metals were also removed when considering the dataset as a whole (“all”), but not from each metal considered in isolation. To assess parallelism across metals (but not within a metal), “difmetal” counts genes repeatedly hit ONLY in different metals (i.e., mutations in the same gene that occur within a metal are counted only once).

In[4]:=
```
data = {{"#hit", "all", "cd", "co", "cu", "mn", "ni", "zn", "difmetal"},
   {1, 186, 43, 63, 18, 47, 17, 20, 200}, {2, 15, 2, 5, 0, 2, 2, 2, 9},
   {3, 0, 1, 1, 0, 1, 1, 2, 1}, {4, 3, 0, 1, 0, 1, 0, 0, 0}, {5, 3, 0, 0, 0, 0, 0, 1, 0},
   {6, 1, 0, 1, 0, 0, 0, 0, 0}, {7, 0, 0, 0, 0, 0, 0, 0, 0}, {8, 1, 0, 0, 0, 0, 0, 0, 0},
   {9, 0, 0, 0, 0, 0, 0, 0, 0}, {10, 0, 0, 0, 0, 0, 0, 0, 0}, {11, 0, 0, 0, 0, 0, 0, 0, 0},
   {12, 0, 0, 0, 0, 0, 0, 0, 0}, {13, 1, 0, 0, 0, 1, 0, 0, 0}};
MatrixForm[data]
```

Out[5]//MatrixForm=

| #hit | all | cd | co | cu | mn | ni | zn | difmetal |
|---|---|---|---|---|---|---|---|---|
| 1 | 186 | 43 | 63 | 18 | 47 | 17 | 20 | 200 |
| 2 | 15 | 2 | 5 | 0 | 2 | 2 | 2 | 9 |
| 3 | 0 | 1 | 1 | 0 | 1 | 1 | 2 | 1 |
| 4 | 3 | 0 | 1 | 0 | 1 | 0 | 0 | 0 |
| 5 | 3 | 0 | 0 | 0 | 0 | 0 | 1 | 0 |
| 6 | 1 | 0 | 1 | 0 | 0 | 0 | 0 | 0 |
| 7 | 0 | 0 | 0 | 0 | 0 | 0 | 0 | 0 |
| 8 | 1 | 0 | 0 | 0 | 0 | 0 | 0 | 0 |
| 9 | 0 | 0 | 0 | 0 | 0 | 0 | 0 | 0 |
| 10 | 0 | 0 | 0 | 0 | 0 | 0 | 0 | 0 |
| 11 | 0 | 0 | 0 | 0 | 0 | 0 | 0 | 0 |
| 12 | 0 | 0 | 0 | 0 | 0 | 0 | 0 | 0 |
| 13 | 1 | 0 | 0 | 0 | 1 | 0 | 0 | 0 |

The number of independent mutations (leading to different SNPs) across all metals:

In[6]:=
```
Drop[data⟦All, 2⟧, 1].Drop[data⟦All, 1⟧, 1]
```

Out[6]= 270

The number of independent mutations (leading to different SNPs) across all metals, lumping together any mutations in the same gene that occur in the same metal (for testing parallelism across metals):

In[7]:=
```
Drop[data⟦All, 9⟧, 1].Drop[data⟦All, 1⟧, 1]
```

Out[7]= 221

#### Analyses

##### Cadmium - significant parallelism

In[8]:=
```
data⟦All, 3⟧⟦1⟧
```

Out[8]= cd

In[9]:=
```
obsdata = Drop[data⟦All, 3⟧, 1]
```

Out[9]= {43, 2, 1, 0, 0, 0, 0, 0, 0, 0, 0, 0, 0}

In[10]:=
```
nummut = obsdata.Drop[data[[All, 1]], 1]
```
Out[10]=
```
50
```

Of these, the number of multiply hit genes were:

In[11]:=
```
nummultiple = Total[Drop[data[[All, 3]], 2]]
```
Out[11]=
```
3
```

The probability that all mutations hit separate genes is:

In[12]:=
```
pr1[nummut]
```
Out[12]=
```
0.830378
```

The probability that either all mutations hit separate genes or at most one gene is hit twice is:

In[13]:=
```
pr1[nummut] + pr2[nummut]
```
Out[13]=
```
0.985488
```

Thus, p<0.02 for seeing more than one gene hit multiple times (there were three for cadmium), let alone having a gene hit more than two times (one gene hit three times).

Alternatively, we could switch to simulating data draws from a multinomial.

In[14]:=
```
SeedRandom[129 831]
```
Out[14]=
```
RandomGeneratorState[ Method: ExtendedCA
                      State hash: −4 493 126 053 975 581 468 ]
```

Generating random draws of nummut mutations, each of which could occur in any one of 6607 possible genes (with an equal probability of mutating), repeating this 1000 times:

In[15]:=
```
tab = Table[1 / 6607, {i, 1, 6607}];
rantab = Table[BinCounts[
    RandomInteger[MultinomialDistribution[nummut, tab]], {0, 14, 1}], {i, 1, 1000}];
```

The mean number of hits observed (j counts the number of mutations, accounting for genes hit 1, 2, ...13 times):

In[16]:=
```
obsdata.Table[j (j), {j, 1, 13}]
-------------------------------- // N
  obsdata.Table[j, {j, 1, 13}]
```
Out[16]=
```
1.2
```

Is higher than the mean numbers of all of 1000 randomizations:

In[17]:=
```
meantab = Table[rantab[[i]].Table[j^2, {j, 0, 13}] / rantab[[i]].Table[j, {j, 0, 13}], {i, 1, 1000}] // N;
Max[meantab]
```
Out[18]=
```
1.12
```

The 95% quantile for mean number of genes hit:

In[19]:=
```
{Quantile[meantab, 0.025], Quantile[meantab, 0.5], Quantile[meantab, 0.975]}
```
Out[19]=
```
{1., 1., 1.04}
```

Thus both the mean number of hits per gene is significant (p<0.001), as is the number of genes hit multiple times.

#### Cobalt - significant parallelism

In[20]:=
```
data[[All, 4]][[1]]
```
Out[20]=
```
co
```

In[21]:=
```
obsdata = Drop[data[[All, 4]], 1]
```
Out[21]=
```
{63, 5, 1, 1, 0, 1, 0, 0, 0, 0, 0, 0, 0}
```

In[22]:=
```
nummut = obsdata.Drop[data[[All, 1]], 1]
```
Out[22]=
```
86
```

Of these, the number of multiply hit genes were:

In[23]:=
```
nummultiple = Total[Drop[data[[All, 4]], 2]]
```
Out[23]=
```
8
```

The probability that all mutations hit separate genes is:

In[24]:=
```
pr1[nummut]
```
Out[24]=
```
0.573726
```

The probability that either all mutations hit separate genes or at most one gene is hit twice is:

In[25]:=
```
pr1[nummut] + pr2[nummut]
```
Out[25]=
```
0.895249
```

Thus, there are so many hits to cobalt that it’s still reasonably likely (p>0.01) that we would see one double hit.  Allowing two double hits though, the observed data is unlikely to be seen.

In[26]:=
```
pr1[nummut] + pr2[nummut] + pr22[nummut]
```
Out[26]=
```
0.983922
```

That is, p<0.02 for seeing more than two genes hit multiple times (there were eight for cobalt).

Alternatively, we could switch to simulating data draws from a multinomial.

In[27]:=
```
SeedRandom[54 219]
```
Out[27]=
```
RandomGeneratorState[ Method: ExtendedCA
                      State hash: 466 435 363 971 759 394 ]
```

Generating random draws of nummut mutations, each of which could occur in any one of 6607 possible genes (with an equal probability of mutating), repeating this 1000 times:

In[28]:=
```
tab = Table[1 / 6607, {i, 1, 6607}];
rantab = Table[BinCounts[
    RandomInteger[MultinomialDistribution[nummut, tab]], {0, 14, 1}], {i, 1, 1000}];
```

The mean number of hits observed (j counts the number of mutations, accounting for genes hit 1, 2, ...13 times):

In[29]:=
$$\frac{\text{obsdata.Table[j (j), \{j, 1, 13\}]}}{\text{obsdata.Table[j, \{j, 1, 13\}]}} \text{ // N}$$

Out[29]=
```
1.67442
```

Is higher than the mean numbers of all of 1000 randomizations:

In[30]:=
$$\text{meantab} = \text{Table}\left[\frac{\text{rantab[[i]].Table}\left[j^2, \{j, 0, 13\}\right]}{\text{rantab[[i]].Table[j, \{j, 0, 13\}]}}, \{i, 1, 1000\}\right] \text{ // N;}$$

```
Max[meantab]
```
Out[31]=
```
1.09302
```

The 95% quantile for mean number of genes hit:

In[32]:=
```
{Quantile[meantab, 0.025], Quantile[meantab, 0.5], Quantile[meantab, 0.975]}
```
Out[32]=
```
{1., 1., 1.04651}
```

Thus both the mean number of hits per gene is significant (p<0.001), as is the number of genes hit multiple times.

#### Copper - no parallelism

In[33]:=
```
data⟦All, 5⟧⟦1⟧
```
Out[33]=
```
cu
```
In[34]:=
```
obsdata = Drop[data⟦All, 5⟧, 1]
```
Out[34]=
```
{18, 0, 0, 0, 0, 0, 0, 0, 0, 0, 0, 0, 0}
```
In[35]:=
```
nummut = obsdata.Drop[data⟦All, 1⟧, 1]
```
Out[35]=
```
18
```
Of these, the number of multiply hit genes were:

In[36]:=
```
nummultiple = Total[Drop[data⟦All, 5⟧, 2]]
```
Out[36]=
```
0
```
The probability that all mutations hit separate genes is:

In[37]:=
```
pr1[nummut]
```
Out[37]=
```
0.977089
```
Thus, it is very likely (p>0.98) to see no multiply-hit genes (none were observed for copper).

#### Manganese - significant parallelism

For manganese, we drop the putative mutators (MnBM14 and MnBM42)

In[38]:=
```
data⟦All, 6⟧⟦1⟧
```
Out[38]=
```
mn
```
In[39]:=
```
obsdata = Drop[data⟦All, 6⟧, 1]
```
Out[39]=
```
{47, 2, 1, 1, 0, 0, 0, 0, 0, 0, 0, 0, 1}
```
In[40]:=
```
nummut = obsdata.Drop[data⟦All, 1⟧, 1]
```
Out[40]=
```
71
```
Of these, the number of multiply hit genes were:

In[41]:=
```
nummultiple = Total[Drop[data⟦All, 6⟧, 2]]
```
Out[41]=
```
5
```

The probability that all mutations hit separate genes is:

In[42]:=
```
pr1[nummut]
```
Out[42]=
```
0.6856
```

The probability that either all mutations hit separate genes or at most one gene is hit twice is:

In[43]:=
```
pr1[nummut] + pr2[nummut]
```
Out[43]=
```
0.946226
```

Thus, p~0.05 for seeing more than one gene hit multiple times (there were five for manganese).

Considering one or two genes hit twice (as well as all genes hit only once):

In[44]:=
```
pr1[nummut] + pr2[nummut] + pr22[nummut]
```
Out[44]=
```
0.99482
```

That is, p<0.01 for seeing more than two genes hit multiple times (there were five for manganese).

Alternatively, we could switch to simulating data draws from a multinomial.

In[45]:=
```
SeedRandom[77127]
```
Out[45]=
```
RandomGeneratorState[ Method: ExtendedCA
                      State hash: 8628400527046742293 ]
```

Generating random draws of nummut mutations, each of which could occur in any one of 6607 possible genes (with an equal probability of mutating), repeating this 1000 times:

In[46]:=
```
tab = Table[1 / 6607, {i, 1, 6607}];
rantab = Table[BinCounts[
    RandomInteger[MultinomialDistribution[nummut, tab]], {0, 14, 1}], {i, 1, 1000}];
```

The mean number of hits observed (j counts the number of mutations, accounting for genes hit 1, 2, ...13 times):

In[47]:=
$$\frac{\texttt{obsdata.Table[j (j), \{j, 1, 13\}]}}{\texttt{obsdata.Table[j, \{j, 1, 13\}]}} \texttt{ // N}$$
Out[47]=
```
3.50704
```

Is higher than the mean numbers of all of 1000 randomizations:

In[48]:=
```
meantab = Table[(rantab⟦i⟧.Table[j^2, {j, 0, 13}])/(rantab⟦i⟧.Table[j, {j, 0, 13}]), {i, 1, 1000}] // N;
Max[meantab]
```
Out[49]=
```
1.08451
```

The 95% quantile for mean number of genes hit:

In[50]:=
```
{Quantile[meantab, 0.025], Quantile[meantab, 0.5], Quantile[meantab, 0.975]}
```
Out[50]=
```
{1., 1., 1.05634}
```

Even if we drop the gene hit 13 times in manganese (*CDC25*), the result is highly significant and outside the range of all 1000 randomizations:

In[51]:=
```
(Drop[obsdata, -1].Table[j (j), {j, 1, 12}])/(Drop[obsdata, -1].Table[j, {j, 1, 12}]) // N
```
Out[51]=
```
1.37931
```

Thus both the mean number of hits per gene is significant (p<0.001), as is the number of genes hit multiple times.

#### Nickle - significant parallelism

In[71]:=
```
data⟦All, 7⟧⟦1⟧
```
Out[71]=
```
ni
```

In[72]:=
```
obsdata = Drop[data⟦All, 7⟧, 1]
```
Out[72]=
```
{17, 2, 1, 0, 0, 0, 0, 0, 0, 0, 0, 0, 0}
```

In[73]:=
```
nummut = obsdata.Drop[data⟦All, 1⟧, 1]
```
Out[73]=
```
24
```

Of these, the number of multiply hit genes were:

In[74]:=
```
nummultiple = Total[Drop[data⟦All, 7⟧, 2]]
```
Out[74]=
```
3
```

The probability that all mutations hit separate genes is:

In[75]:=
```
pr1[nummut]
```
Out[75]=
```
0.959039
```

The probability that either all mutations hit separate genes or at most one gene is hit twice is:

In[76]:=
```
pr1[nummut] + pr2[nummut]
```
Out[76]=
```
0.999242
```

Thus, p<0.001 for seeing more than one gene hit multiple times (there were three for nickel), let alone having genes hit more than two times.

Alternatively, we could switch to simulating data draws from a multinomial.

In[77]:=
```
SeedRandom[32412]
```
Out[77]=
```
RandomGeneratorState[ Method: ExtendedCA
                      State hash: −8692798250509348668 ]
```

Generating random draws of nummut mutations, each of which could occur in any one of 6607 possible genes (with an equal probability of mutating), repeating this 1000 times:

In[78]:=
```
tab = Table[1 / 6607, {i, 1, 6607}];
rantab = Table[BinCounts[
    RandomInteger[MultinomialDistribution[nummut, tab]], {0, 14, 1}], {i, 1, 1000}];
```

The mean number of hits observed (j counts the number of mutations, accounting for genes hit 1, 2, ...13 times):

In[79]:=
$$\frac{\text{obsdata.Table[j (j), \{j, 1, 13\}]}}{\text{obsdata.Table[j, \{j, 1, 13\}]}} \text{ // N}$$

Out[79]=
```
1.41667
```

Is higher than the mean numbers of all of 1000 randomizations:

$$\text{meantab = Table}\left[\frac{\text{rantab[[i]].Table}\left[j^2, \{j, 0, 13\}\right]}{\text{rantab[[i]].Table[j, \{j, 0, 13\}]}}, \{i, 1, 1000\}\right] \text{ // N;}$$

```
Max[meantab]
```
Out[ ]=
```
1.09302
```

The 95% quantile for mean number of genes hit:

```
{Quantile[meantab, 0.025], Quantile[meantab, 0.5], Quantile[meantab, 0.975]}
```
Out[ ]=
```
{1., 1., 1.04651}
```

Thus both the mean number of hits per gene is significant (p<0.001), as is the number of genes hit multiple times.

#### Zinc - significant parallelism

In[83]:=
```
data⟦All, 8⟧⟦1⟧
```
Out[83]=
```
zn
```
In[84]:=
```
obsdata = Drop[data⟦All, 8⟧, 1]
```
Out[84]=
```
{20, 2, 2, 0, 1, 0, 0, 0, 0, 0, 0, 0, 0}
```
In[85]:=
```
nummut = obsdata.Drop[data⟦All, 1⟧, 1]
```
Out[85]=
```
35
```
Of these, the number of multiply hit genes were:

In[86]:=
```
nummultiple = Total[Drop[data⟦All, 8⟧, 2]]
```
Out[86]=
```
5
```
The probability that all mutations hit separate genes is:

In[87]:=
```
pr1[nummut]
```
Out[87]=
```
0.913736
```
The probability that either all mutations hit separate genes or at most one gene is hit twice is:

In[88]:=
```
pr1[nummut] + pr2[nummut]
```
Out[88]=
```
0.996449
```
Thus, $p<0.01$ for seeing more than one gene hit multiple times (there were five for zinc), let alone having genes hit more than two times.

Alternatively, we could switch to simulating data draws from a multinomial.

In[89]:=
```
SeedRandom[82 712]
```
Out[89]=
```
RandomGeneratorState[ Method: ExtendedCA
                      State hash: −1 951 316 623 039 499 653 ]
```
Generating random draws of nummut mutations, each of which could occur in any one of 6607 possible genes (with an equal probability of mutating), repeating this 1000 times:

In[90]:=
```
tab = Table[1 / 6607, {i, 1, 6607}];
rantab = Table[BinCounts[
    RandomInteger[MultinomialDistribution[nummut, tab]], {0, 14, 1}], {i, 1, 1000}];
```

The mean number of hits observed (j counts the number of mutations, accounting for genes hit 1, 2, ...13 times):

In[91]:=
```
obsdata.Table[j (j), {j, 1, 13}] / obsdata.Table[j, {j, 1, 13}] // N
```
Out[91]=
```
2.02857
```

Is higher than the mean numbers of all of 1000 randomizations:

In[92]:=
```
meantab = Table[rantab⟦i⟧.Table[j^2, {j, 0, 13}] / rantab⟦i⟧.Table[j, {j, 0, 13}], {i, 1, 1000}] // N;
Max[meantab]
```
Out[93]=
```
1.11429
```

The 95% quantile for mean number of genes hit:

In[94]:=
```
{Quantile[meantab, 0.025], Quantile[meantab, 0.5], Quantile[meantab, 0.975]}
```
Out[94]=
```
{1., 1., 1.05714}
```

Thus both the mean number of hits per gene is significant ($p<0.001$), as is the number of genes hit multiple times.

#### All metals together - significant parallelism

In[64]:=
```
data⟦All, 2⟧⟦1⟧
```
Out[64]=
```
all
```

In[65]:=
```
obsdata = Drop[data⟦All, 2⟧, 1]
```
Out[65]=
```
{186, 15, 0, 3, 3, 1, 0, 1, 0, 0, 0, 0, 1}
```

In[66]:=
```
nummut = obsdata.Drop[data⟦All, 1⟧, 1]
```
Out[66]=
```
270
```

Of these, the number of multiply hit genes were:

In[67]:=
```
nummultiple = Total[Drop[data⟦All, 2⟧, 2]]
```
Out[67]=
```
24
```

The probability that all mutations hit separate genes is:

In[68]:=
```
pr1[nummut]
```
Out[68]=
```
0.00380006
```

The probability that either all mutations hit separate genes or at most one gene is hit twice is:

In[69]:=
```
pr1[nummut] + pr2[nummut]
```
Out[69]=
```
0.0255733
```

Thus, there are so many hits overall that it's still reasonably likely (p>0.01) that we would see one double hit.  Allowing two double hits isn't even enough:

In[70]:=
```
pr1[nummut] + pr2[nummut] + pr22[nummut]
```
Out[70]=
```
$Aborted
```

We thus switch to simulating data draws from a multinomial.

In[ ]:=
```
SeedRandom[2120]
```
Out[ ]=
```
RandomGeneratorState[ Method: ExtendedCA
                      State hash: 8 852 878 573 749 685 000 ]
```

Generating random draws of nummut mutations, each of which could occur in any one of 6607 possible genes (with an equal probability of mutating), repeating this 1000 times:

In[ ]:=
```
tab = Table[1 / 6607, {i, 1, 6607}];
rantab = Table[BinCounts[
    RandomInteger[MultinomialDistribution[nummut, tab]], {0, 14, 1}], {i, 1, 1000}];
```

The mean number of hits observed (j counts the number of mutations, accounting for genes hit 1, 2, ...13 times):

In[ ]:=
```
obsdata.Table[j (j), {j, 1, 13}] / obsdata.Table[j, {j, 1, 13}] // N
```
Out[ ]=
```
2.36296
```

Is higher than the mean numbers of all of 1000 randomizations:

In[ ]:=
```
meantab = Table[rantab[[i]].Table[j^2, {j, 0, 13}] / rantab[[i]].Table[j, {j, 0, 13}], {i, 1, 1000}] // N;
Max[meantab]
```
Out[ ]=
```
1.11111
```

The 95% quantile for mean number of genes hit:

In[∘]:=
```
{Quantile[meantab, 0.025], Quantile[meantab, 0.5], Quantile[meantab, 0.975]}
```
Out[∘]=
```
{1.00741, 1.03704, 1.07407}
```

Even if we drop the gene hit 13 times in manganese (*CDC25*), the result is highly significant and outside the range of all 1000 randomizations:

In[∘]:=
$$\frac{\text{Drop[obsdata, -1].Table[j (j), \{j, 1, 12\}]}}{\text{Drop[obsdata, -1].Table[j, \{j, 1, 12\}]}} \text{ // N}$$
Out[∘]=
```
1.8249
```

**Double hits:** The 95% quantile for expected number of double hit genes is 1-10 (median of 5), whereas 15 were observed:

In[∘]:=
```
{Quantile[rantab⟦All, 3⟧, 0.025],
 Quantile[rantab⟦All, 3⟧, 0.5], Quantile[rantab⟦All, 3⟧, 0.975]}
```
Out[∘]=
```
{1, 5, 10}
```

**More than two hits:** The 95% quantile for expected number of triple-plus hit genes is 0-1 (median of 0), whereas 9 were observed:

In[∘]:=
```
{Quantile[Sum[rantab⟦All, i⟧, {i, 4, 10}], 0.025],
 Quantile[Sum[rantab⟦All, i⟧, {i, 4, 10}], 0.5],
 Quantile[Sum[rantab⟦All, i⟧, {i, 4, 10}], 0.975]}
```
Out[∘]=
```
{0, 0, 1}
```

Number of hits per gene and count of the maximum time that # of hits was observed:

In[∘]:=
```
Table[{i - 1, Max[rantab⟦All, i⟧]}, {i, 1, 10}]
```
Out[∘]=
```
{{0, 6351}, {1, 270}, {2, 12}, {3, 2}, {4, 1}, {5, 0}, {6, 0}, {7, 0}, {8, 0}, {9, 0}}
```

Only 6.7% of simulations had any genes hit more than twice, whereas 9 were observed:

In[∘]:=
```
Total[Sum[rantab⟦All, i⟧, {i, 4, 10}]] / 1000.
```
Out[∘]=
```
0.067
```

Thus both the mean number of hits per gene (the main test) is significant (p<0.001), as is the number of genes hit twice or more than twice.

#### All metals together (dropping repeated hits in the same gene) - significant parallelism

In[ ]:=
```
data[[All, 9]][[1]]
```
Out[ ]=
```
difmetal
```
In[ ]:=
```
obsdata = Drop[data[[All, 9]], 1]
```
Out[ ]=
```
{200, 9, 1, 0, 0, 0, 0, 0, 0, 0, 0, 0, 0}
```
In[ ]:=
```
nummut = obsdata.Drop[data[[All, 1]], 1]
```
Out[ ]=
```
221
```
Of these, the number of multiply hit genes were:

In[ ]:=
```
nummultiple = Total[Drop[data[[All, 9]], 2]]
```
Out[ ]=
```
10
```
The probability that all mutations hit separate genes is:

In[ ]:=
```
pr1[nummut]
```
Out[ ]=
```
0.0242083
```
The probability that either all mutations hit separate genes or at most one gene is hit twice is:

In[ ]:=
```
pr1[nummut] + pr2[nummut]
```
Out[ ]=
```
0.116349
```
Thus, there are so many hits overall that it’s still reasonably likely (p>0.01) that we would see one double hit.  Allowing two double hits isn’t even enough:

In[ ]:=
```
pr1[nummut] + pr2[nummut] + pr22[nummut]
```
Out[ ]=
```
0.290613
```
We thus switch to simulating data draws from a multinomial.

In[ ]:=
```
SeedRandom[8329]
```
Out[ ]=
```
RandomGeneratorState[ Method: ExtendedCA  State hash: -4 294 397 200 794 781 781 ]
```
Generating random draws of nummut mutations, each of which could occur in any one of 6607 possible genes (with an equal probability of mutating), repeating this 1000 times:

In[∘]:=
```
tab = Table[1 / 6607, {i, 1, 6607}];
rantab = Table[BinCounts[
    RandomInteger[MultinomialDistribution[nummut, tab]], {0, 14, 1}], {i, 1, 1000}];
```

The mean number of hits observed (j counts the number of mutations, accounting for genes hit 1, 2, ...13 times):

In[∘]:=
```
obsdata.Table[j (j), {j, 1, 13}]
-------------------------------- // N
  obsdata.Table[j, {j, 1, 13}]
```

Out[∘]=
```
1.1086
```

Is above the highest of all 1000 randomizations:

In[∘]:=
```
meantab = Table[rantab⟦i⟧.Table[j^2, {j, 0, 13}] / rantab⟦i⟧.Table[j, {j, 0, 13}], {i, 1, 1000}] // N;
Max[meantab]
```

Out[∘]=
```
1.0905
```

There are 4/1000 (p~0.004) randomizations with such an extreme degree of parallelism as observed, even in this conservative test where all parallel mutations in the same gene were dropped as were metals that were hit but with a SNP seen in another metal:

In[∘]:=
```
Length[Select[meantab, # ≥ obsdata.Table[j (j), {j, 1, 13}] / obsdata.Table[j, {j, 1, 13}] &]]
```

Out[∘]=
```
0
```

The 95% quantile for mean number of genes hit:

In[∘]:=
```
{Quantile[meantab, 0.025], Quantile[meantab, 0.5], Quantile[meantab, 0.975]}
```

Out[∘]=
```
{1., 1.02715, 1.0724}
```

**Double hits:** The 95% quantile for expected number of double hit genes is 1-7 (median of 3) is consistent with the 7 observed:

In[∘]:=
```
{Quantile[rantab⟦All, 3⟧, 0.025],
  Quantile[rantab⟦All, 3⟧, 0.5], Quantile[rantab⟦All, 3⟧, 0.975]}
```

Out[∘]=
```
{0, 3, 7}
```

**More than two hits:** The 95% quantile for expected number of triple-plus hit genes is 0-1 (median of 0), whereas 1 was observed:

In[ ]:=
```
{Quantile[Sum[rantab[[All, i]], {i, 4, 10}], 0.025],
 Quantile[Sum[rantab[[All, i]], {i, 4, 10}], 0.5],
 Quantile[Sum[rantab[[All, i]], {i, 4, 10}], 0.975]}
```

Out[ ]=
```
{0, 0, 1}
```

Number of hits per gene and count of the maximum time that # of hits was observed:

In[ ]:=
```
Table[{i - 1, Max[rantab[[All, i]]]}, {i, 1, 10}]
```

Out[ ]=
```
{{0, 6396}, {1, 221}, {2, 10}, {3, 1}, {4, 0}, {5, 0}, {6, 0}, {7, 0}, {8, 0}, {9, 0}}
```

Only 4.1% of simulations had any genes hit more than twice, whereas one was observed:

In[ ]:=
```
Total[Sum[rantab[[All, i]], {i, 4, 10}]] / 1000.
```

Out[ ]=
```
0.033
```

Thus both the mean number of hits per gene (the main test) is significant (p~0.004), as is the number of genes hit more than twice (PMA1)
